# Design of an orally bioavailable small molecule that modulates the microtubule-associated protein tau’s pre-mRNA splicing

**DOI:** 10.1101/2025.04.30.651234

**Authors:** Peiyuan Zhang, Amirhossein Taghavi, Masahito Abe, Haruo Aikawa, Yoshihiro Akahori, Jonathan L. Chen, Yuquan Tong, Jared T. Baisden, Michael D. Cameron, Jessica L. Childs-Disney, Matthew D. Disney

**Author notes:** Author to whom correspondence is addressed.

## Abstract

Frontotemporal dementia with parkinsonism linked to chromosome 17 (FTDP-17) is caused by the aberrant alternative pre-mRNA splicing of microtubule-associated protein tau (*MAPT*) exon 10, the inclusion of which encodes for a toxic tau protein harboring four microtube domains (4R tau). Here, we describe the design of an RNA-targeted small molecule that thermodynamically stabilizes the structure of a pre-mRNA splicing regulator element in the *MAPT* pre-mRNA exon 10-intron junction to reduce the inclusion of exon 10 and hence 4R tau abundance. Structure-based drug design was used to obtain compounds that form a network of specific interactions to the RNA including multiple interactions between a one nucleotide A-bulge and the Hoogsteen face of a closing GC base pair, the latter of which was enabled by the design of base triple interactions. A battery of assays revealed that the compound binds the target in vitro and in cells and affects pre-mRNA splicing in various cellular models including primary neurons from a human tau (htau) knock-in mouse model. The orally bioavailable compound was administered *per os* (*p.o.*), where treatment diminished exon 10 inclusion, and reduced the 4R tau protein isoform. Further, the molecule mitigated cellular pathologies and behavioral phenotypes observed in the htau transgenic mouse model. This study provides a potentially general pipeline to design compounds that target RNAs and affect disease pathways and deliver compounds that have oral bioavailability and blood-brain barrier penetrance.

## INTRODUCTION

Over 90% of pre-mRNAs are alternatively spliced, the outcome of which can be affected by a variety of factors, including the sequence and structure of the pre-mRNA and the recognition thereof by *trans*-splicing factors such as the spliceosome and other proteins.^1, 2^ Many diseases are caused by aberrant alternative pre-mRNA splicing events including inherited genetic diseases, cancer, and diseases of aging.^3, 4^ The mechanistic underpinnings of disease-relevant alternative pre-mRNA splicing events can provide insights into the development of targeted therapeutics.

The aberrant alternative splicing of the microtubule-associated protein tau (*MAPT*) gene, which encodes the protein tau, causes the genetically defined disease frontotemporal dementia with parkinsonism linked to chromosome 17 (FTDP-17).^5^ The gene comprises 16 exons, including alternatively spliced exons 2, 3, and 10, yielding six isoforms containing eight constitutive exons that affect the number of microtubule binding repeat (MTBR) domains at the carboxy-terminus.^6^ Exclusion or inclusion of exon 10 leads to three- or four-repeat MTBR isoforms (3R tau or 4R tau), respectively. In the healthy adult brain, 3R and 4R tau proteins are expressed approximately equally,^7^ where 4R tau has additional phosphorylation sites that change protein solubility and ultimately lead to insoluble aggregates.^8^ In FTDP-17, a mutation in an RNA regulatory structure at the exon 10-intron junction dysregulates alternative splicing, tipping the balance of 3R:4R towards the aggregation-prone 4R isoform.^9^ In addition, an increase in 4R tau without a genetic disposition has been shown to contribute to Alzheimer’s disease (AD), as evidenced by aggregation of 4R in human brains.^8^ The contributions of 4R tau to disease are supported by the delivery of antisense oligonucleotides (ASOs) that switch splicing to increase 4R tau in a humanized tau (htau) mouse model, which causes behavioral defects observed in AD.^10^ Conversely, tipping the balance of 3R and 4R tau in the opposite direction (too much 3R) results in Pick disease, which is characterized by 3R tau aggregates (Pick bodies),^11^ neuronal loss, and cortical atrophy.^12^

Mechanistic studies used in conjunction with human genetics have deciphered the mechanism of FTDP-17. Various mutations, including a C-to-U intronic mutation (cytosine to uracil; C[+14]U; commonly referred to as disinhibition-dementia-parkinsonism-amyotrophy complex (DDPAC)), occur downstream of the 5′ splice site of tau exon 10 (Figure 1A).^13^ This mutation in the RNA’s structure, as well as others, decreases the thermodynamic stability of the hairpin structure formed at the exon 10- intron 10 junction, a splicing regulatory element (SRE).^14^ This destabilization increases U1 small nuclear ribonuclear protein (snRNP) binding and hence exon 10 inclusion and 4R tau expression (Figure 1A).^14^ The proportion of 4R to 3R tau is therefore greater in FTDP-17 patients (Figure 1A). Conversely, introduction of mutations that thermodynamically stabilize the SRE hairpin causes a reduction of the 4R/3R ratio,^10^ suggesting a potential therapeutic strategy for FTDP-17. That is, small molecules that bind to the RNA structure in *MAPT* pre-mRNA could limit U1 snRNP binding at the exon 10-intron junction and reduce the formation of toxic 4R tau protein.

**Figure 1.**
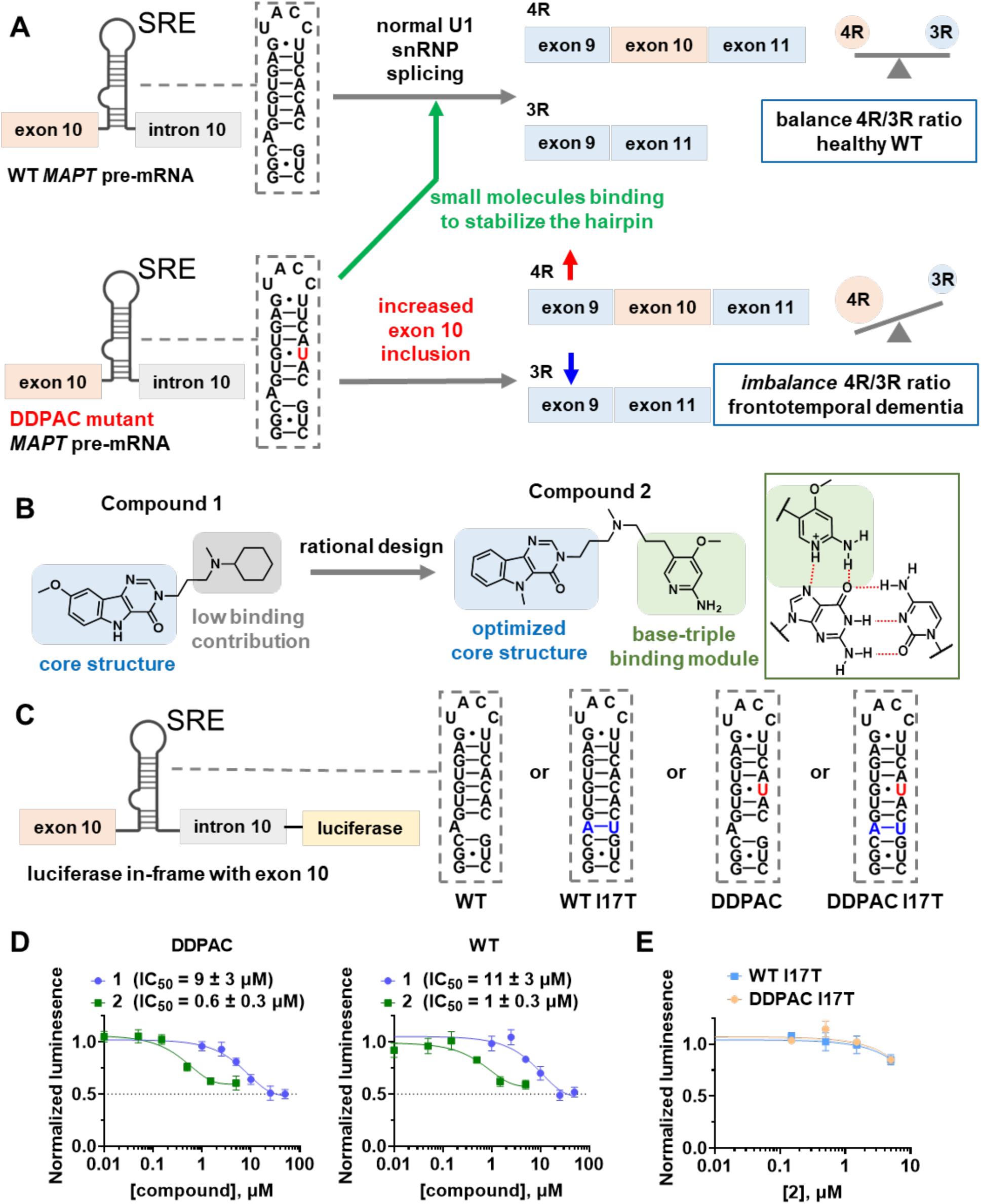
Rational design of 2, which modulates alternative splicing of *MAPT* exon 10 by binding to a splicing regulatory element (SRE). (A) Schematic of alternative splicing of exons 9, 10, and 11 of *MAPT* pre-mRNA. U1 snRNP binds and unfolds the *MAPT* SRE at the exon 10−intron 10 junction, leading to exon 10 inclusion. DDPAC *MAPT*, a mutation associated with FTDP-17, has **+14 C to U** mutation that destabilizes the hairpin, promoting U1 snRNP binding and hence exon 10 inclusion, producing higher levels of 4R tau and an imbalance of the 4R/3R ratio. The binding of small molecules to the A-bulge site within SRE stabilizes the hairpin, increasing exon 10 skipping and production of 3R tau. (B) Structure of parent compound **1**, which was previously identified.^17^ Compound **2** was rationally designed to form a base triple with a GC pair that closes the A-bulge, the binding site of **1**. (C) Secondary structures of various *MAPT* SRE RNAs used in this study. The mini-gene luciferase reporters express luciferase in-frame with exon 10. The I17T mutants abolish the small molecule binding site by converting the A-bulge to an AU base pair. (D) HeLa cells transfected with a DDPAC or WT *MAPT* mini-gene reporter treated with **1** or **2**, showing inhibition of luciferase activity in a dose dependent manner (n = 12 biological replicates for vehicle; n = 4 biological replicates for compound treatment). The highest concentration of compound **2** tested was 5 µM as cell viability was reduced by 17 ± 3% at the 12.5 µM dose (Figure S5). (E) Effect of **1** and **2** on the alternative splicing of WT I17T or DDPAC I17T mutant luciferase reporters (n = 4 biological replicates). Error bars indicate SD. Each biological replicate has two technical replicates.

Various small molecules have been discovered that bind to the SRE at the exon 10-intron junction, including the topoisomerase inhibitor mitoxantrone and aminoglycosides.^15, 16^ Previously, a series of small molecules, including **S1** (Figure S1), that binds a well-folded adenine-containing bulge (A-bulge) region of the *MAPT* SRE and modulate pre-mRNA splicing were identified from a database of experimentally derived RNA-small molecule interactions named Inforna.^17^ This method of lead identification searches for overlap between experimentally derived interactions and secondary structural elements present in cellular RNAs, outputting potential small molecule binders as well as associated metrics of selectivity and affinity. Using these small molecules as leads, a collection of compounds, including **1** (Figure S1), was later identified that promoted exon skipping by directly targeting *MAPT* pre-mRNA in primary neuron cells harvested from htau transgenic mice.^17^ Although the approach was promising in identifying drug-like chemical matter that bound the RNA, the IC_50_ of **1** for directing *MAPT* exon 10 splicing was around 10 µM in a cellular mini-gene splicing assay.^17^

A general and pervasive challenge in the development of RNA-targeted small molecule medicines is their lead optimization, particularly for potency, selectively, and in vivo efficacy. One such strategy relies on targeting multiple sites within an RNA with a single molecule, which has been effective for RNA repeat expansions where small molecules selectively alleviate disease-associated phenotypes in patient-derived cells and mouse models.^18–22^ In this work, we aimed to develop a general strategy to improve the potency and selectivity of small molecules that target non-repeating RNA targets utilizing *MAPT* pre-mRNA splicing as a proof-of-concept, while providing orally bioavailable compounds that cross the blood-brain barrier (BBB). In particular, we leveraged a base triple formation^23^ design strategy around **1**, affording **2**, which selectively binds the *MAPT* SRE A-bulge and a neighboring GC base pair simultaneously, with sub-micromolar affinity (Figure 1B). Our design strategy takes cues from nature where base triples are frequently occurring motifs formed by edge-to-edge hydrogen bonding, particularly in structured RNAs.^24^ A base triple was first observed between poly(A) and poly(U) in the late 1950s^25, 26^ and predicted to form in tRNAs in the late 1960s.^27^ They were later identified in group I introns,^28^ ribosomal (r)RNAs,^23^ and others,^29, 30^ including triple helices formed in MALAT1,^31^ riboswitches,^32^ telomerase,^33, 34^ and spliceosomal RNAs.^35, 36^ These base triples are key for proper tertiary folding, stability, and function. Our previously elucidated structure of **1** bound to the *MAPT* SRE suggested that a nearby GC base pair was available to form a base triple if the proper module was attached to **1**, here 2-aminopyridine where its protonation under physiological conditions increases its affinity for the negatively charged RNA. In particular, **2** is at least 6-fold more potent in a biochemical assay that measures inhibition of the binding of a model U1 snRNA and at least a 10-fold more potent in cells, as assessed by a cellular mini-gene pre-mRNA splicing assay, than **1**. Target validation studies by Chemical Cross-Linking and Isolation by Pull-down (Chem-CLIP)^37–39^ showed that **2** binds the SRE in cells. Furthermore, orally administered **2** modulated tau splicing in htau mice and improved an associated behavioral phenotype.

## RESULTS

### Design and synthesis of ligands with high affinity for the tau exon 10-intron junction hairpin structure

We previously reported a framework to design and optimize drug-like small molecules that bind tau pre-mRNA and modulate the alternative pre-mRNA splicing of exon 10,^17^ which encodes a microtubule binding domain. Initial molecules were designed from sequence to bind the A-bulge and its closing pairs present in the *MAPT* SRE^14^ using our lead identification strategy, Inforna.^17^ These molecules, with demonstrated binding to the target and cellular activity, informed a pharmacophore model that was used in a virtual screen to generate a small molecule library with potential to modulate exon 10 splicing. Iterative biophysical, biochemical, and cellular studies afforded drug-like compounds that bound to tau pre-mRNA, and the most potent compound from these studies modulated splicing in primary neurons harvested from tau mice.^17^

Three scaffold-diverse compounds, **M1**, **S1**, and **1** (Figure S1), emerged from these studies, and their structures in complex with the tau pre-mRNA target were elucidated by nuclear magnetic resonance (NMR)-restrained molecular dynamics (Protein Data Bank (PDB)^40^ IDs: 6VA3 for **M1**, 6VA2 for **S1** and 6VA4 for **1**).^17^ Each small molecule has a central core that forms a network of interactions to the tau pre-mRNA SRE hairpin structure (Figures S2-S4). Most of the interactions are hydrogen bonds to the closing GC base pairs of the A-bulge and stacking interactions with either one or both closing GC base pairs.

Further investigation of RNA-small molecule interaction patterns in the poses that comprise the NMR-restrained MD structures (20 structures for each small molecule) showed that **M1** was stabilized solely by hydrogen bonds (calculated binding energy = - 6.39 kcal/mol, Table S1). In contrast, the interactions between **S1** and **1** and the RNA were stabilized by stacking interactions and hydrogen bonds (with the phosphodiester backbone (**1** only) and bases), with calculated binding energies of -11.42 kcal/mol and - 8.91 kcal/mol, respectively (Table S1). To determine the likelihood that the small molecules were BBB penetrant, their Central Nervous System Multiparameter Optimization (CNS MPO) scores were calculated.^41, 42^ This analysis, based on the properties of CNS drugs, is calculated by assessing six physicochemical properties (ClogP, ClogD, molecular weight (MW), topological surface area (TPSA), number of hydrogen bond donors (HBD), and pK_a_) which are assigned on a scale from 0 (not CNS drug-like) – 1 (CNS drug-like).^42^ The scores for each property are then summed, with a maximum score of 6; small molecules with scores >4 are likely to be BBB penetrant.^42^ CNS MPO analysis of the small molecules studied herein suggest that **1** and **M1** are likely to be BBB penetrant (CNS MPO = 5.43 and 5.0, respectively) while **S1** is likely not (CNS MPO = 3.64). (See Table S1 for the properties of each small molecule and how each contributes to the CNS MPO score.) Due to its weak interactions (solely hydrogen bonds) with the A-bulge binding pocket, **M1** was eliminated from further consideration.^17^ As **S1** and **1** are different chemical scaffolds with favorable binding energies, they were advanced for further optimization, despite **S1**’s low CNS MPO score.

Careful inspection of the area occupied by **S1** and **1** showed any ligand growth along the X- or Y-axis would either cause steric clashes with the surrounding atoms or project into solvent without providing additional interactions (Figure 2A; right). Therefore, we explored adding functionalities that form additional stabilizing interactions with the RNA target along in the Z-axis, or a T-shaped molecule. Indeed, the structures of the compounds bound to the *MAPT* SRE suggested an opportunity to form additional stabilizing interactions to the Hoogsteen face of the guanosine residue in the A-bulge’s closing GC base pair, that is a base triple.^24^ We took cues from how nature evolves molecular recognition of RNA’s major and minor grooves and triplex-forming oligonucleotides.^24, 29, 30^ In particular, 2-aminopyridines form base triples with GC pairs,^23^ and its protonation under physiological conditions increases its affinity for the negatively charged RNA. Thus, 2-aminopyridine modules were appended onto the central cores of **S1** and **1** to investigate if precise positioning of the base triple module could improve molecular recognition (affinity and selectivity).

**Figure 2.**
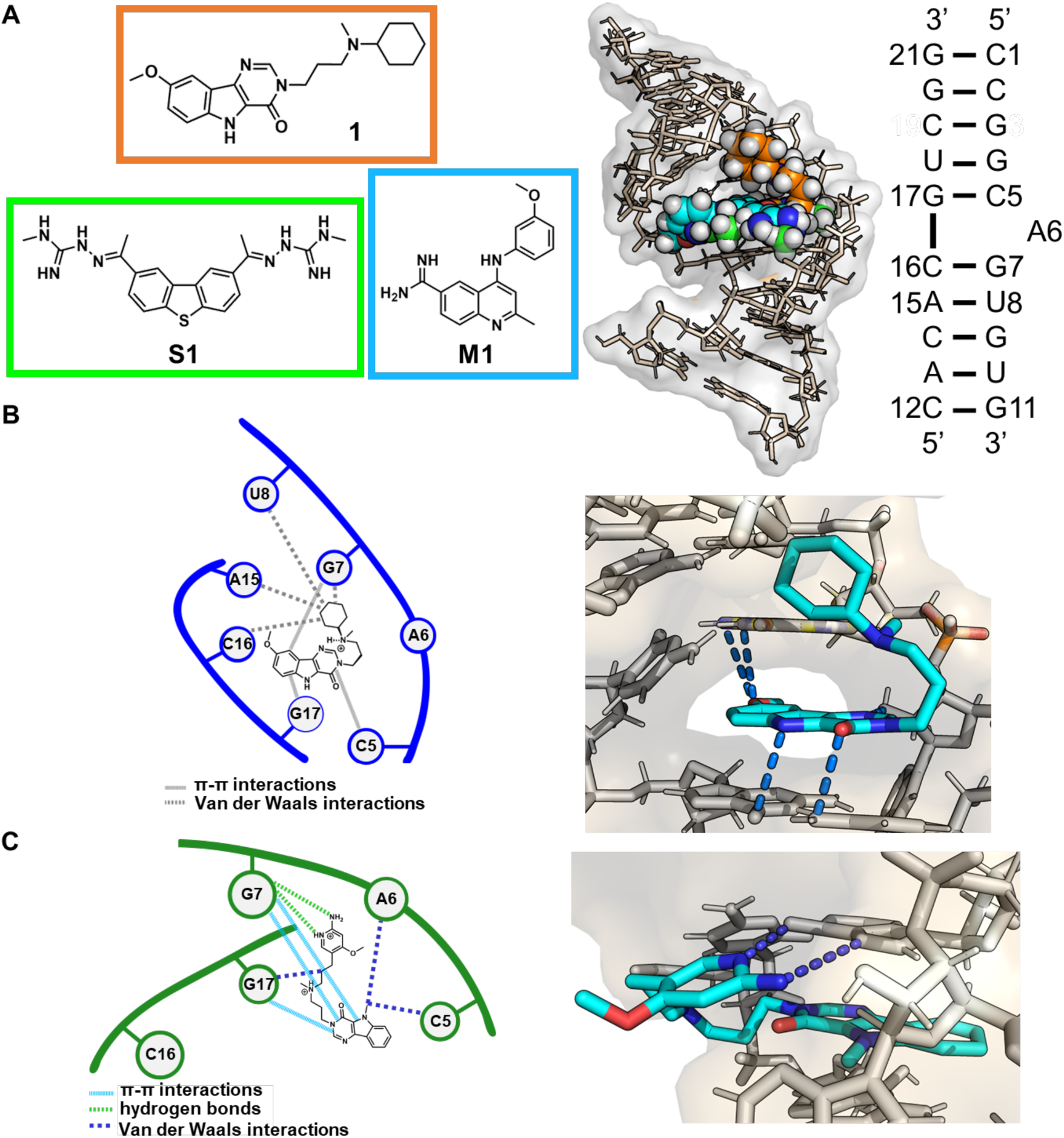
Structures of unbound and ligand-bound WT *MAPT* SRE. (A) Previously reported structures of the bound WT stem mimic duplex.^17^ (B) Structure of the interactions formed between **1** and a duplex model of the WT *MAPT* SRE that was previously reported.^17^ (C) Structure of the interactions formed between **2** and a duplex model of the WT *MAPT* SRE, as determined by NMR-restrained molecular dynamics (MD) simulations. One NOE was observed between the RNA and **2**, which was then used for a restrained based modeling. Light blue lines depict π-stacking interactions; green lines depict hydrogen bonds formed in the base triple; and dark blue lines depict van der Waals interactions. Van der Waals interactions are not shown in the 3D structure. For clarity, only the triple base hydrogen bonds are shown.

As a first step in the optimization process, we designed eight analogs (Figure S1, five based on **S1** and three based on **1**) guided by the previously elucidated NMR solution structures of the parent compounds in complex with the RNA.^17^ Notably, structures of the complexes revealed that two methyl groups on the guanidines present in **S1** and the cyclohexane moiety of **1** do not contribute to binding (Figure 2B). These observations suggest that these methyl groups and the cyclohexane moiety can be modified with 2-aminopyridine modules without affecting molecular recognition. The 2-aminopyridine modules were attached to both parent compounds with flexible linkers such that the linker lies within the major groove and the 2-aminopyridine is positioned so that it may form a base triple with the A-bulge’s closing GC pair (Figures 2 and S1). Three of the five **S1** analogs have a 2-aminopyridine module on one side of the **S1** core (**S2**, **S3,** and **S4**) while the other two have a 2-aminopyridine module on both sides (**S5** and **S6**) (**Figure S1**). Analogs **S2** and **S6** have core binders linked to amino groups of 2-aminopyridine while the other six analogs have core binders linked to aromatic ring carbon atom of 2-aminopyridine. All three analogs of **1** replace the cyclohexane moiety with a 2-aminopyridine module (**2**, **S7**, **S8**; **Figure S1**). These analogs differ by a methyl group present either in the **1** core or in the base triple module. CNS MPO scores were calculated for each analog (Table S1). While the analogs of **1** have CNS MPO scores over 4.0 (likely to be BBB penetrant), the analogs of **S1** gave CNS MPO scores below 3.0 (unsurprising as the CNS MPO for **S1** = 3.64), indicating lower likelihood of BBB penetrance (Table S1).

Small molecule docking was used to investigate the interactions formed between the *MAPT* SRE and the eight 2-aminopyridine conjugates. The RNA target was prepared by removing **1** from the RNA-ligand complex (6VA4); the binding pocket for **S1** (6VA2) is highly similar to that of **1** with an RMSD < 2Å. As expected, based on the binding energies of the parent compounds, derivatives of **S1** had more favorable binding energies than the analogs of **1** (Table S1). Inspection of the lowest energy docked poses of the **1**-base triple conjugates showed the appropriateness of the linker length (the same for all three analogs), placing the 2-aminopyridine ring in the vicinity of one of the GC base pairs that close the A-bulge. In contrast, small molecule docking indicated that the **S1** derivatives were unlikely to form base triple interactions between the 2-aminpyridine and the GC base pairs, mainly due to the longer linker length and thus more rotational degrees of freedom. It also appeared that the longer linkers in **S3** and **S6** formed non-specific interactions with the RNA backbone, pushing the small molecule’s core out of the binding pocket. These results suggest the importance of the linker length for base triple conjugates in particular and modularly assembled compounds in general.

### Bioactivity of 2-aminopyridine conjugates

The eight base triple conjugates (five for **S1** and three for **1**) were screened using a *MAPT* exon 10 splicing mini-gene reporter transfected into HeLa cells. The mini-gene comprises *MAPT* exons 9 – 11 with truncated introns and expresses firefly luciferase in-frame with exon 10, a proxy for expression of 4R tau (Figure 1C).^10^ Four reporters were used in these studies: (i) wild type (WT) *MAPT*; (ii) the DDPAC mutation; (iii) the single mutant, known as WT I17T, containing a U insertion opposite to the A-bulge (lacks the small molecule binding site); and (iv) the double mutant, known as DDPAC I17T (also lacks the small molecule binding site) (Figure 1C).

As an initial measure of bioactivity, the eight conjugates were evaluated for facilitating exon 10 splicing using the DDPAC mini-gene splicing assay at three or four different concentrations (0.1 µM, 0.5 µM, and 2.5 µM for **S1** and analogs thereof; 0.5 µM, 1.5 µM, 5 µM for **1** and analogs thereof; an additional concentration of 15 µM was also studied for **1**). As shown in Figure S6, **S1** and its aminopyridine conjugates had either no (**S2**) or modest activity in the assay. In particular, a ∼20-25% reduction was observed at the 2.5 μM (highest) dose evaluated for **S3**, **S4**, and **S6**. Note that the highest concentration tested was selected based on cell viability studies (**Figure S5**). Of all small molecules studied, **2** excluded exon 10 to the greatest extent, as measured by the reduction of luciferase activity, by 42 ± 6% at the 1.5 µM dose, similar to the reduction observed upon treatment with 15 µM of **1** (42 ± 4%). Because of its activity in the mini- gene assay and its superior CNS MPO score as compared to the **S1** series (CNS MPO scores ranging from 2.12 – 2.74), **2** (CNS MPO = 4.30) was selected for further evaluation.

An extended dose response afforded IC_50_s of 9 ± 3 µM and 0.6 ± 0.3 µM for **1** and **2**, respectively (Figure 1D). A similar dose-dependent reduction in luciferase activity by **2** was also observed with the WT reporter (Figure 1D). [Notably, the maximum reduction in luciferase activity observed for the compounds was ∼50%. A previously reported Vivo-Morpholino ASO (Gene Tools; uses an arginine-rich peptide to facilitate uptake) complementary to the SRE RNA that directs splicing to the 3R isoform (heretofore “tau ASO”)^10^ reduced the *MAPT* mRNA 4R/3R ratio from 2 ± 0.4 (scrambled ASO) to 0.3 ± 0.05 (∼85% reduction) in the DDPAC mini-gene reporter, as determined by RT-qPCR. However, firefly luciferase activity was only reduced by 55 ± 3% in DDPAC mini-gene splicing assay (Figures 3A & S6).] No effect on the viability of HeLa cells upon treatment with **2** was observed up to a concentration of 12.5 µM, 2.5 times higher than its highest dose (5 µM) used in the luciferase assay; at the 12.5 µM dose, cell viability was reduced by 17 ± 3% (Figure S5). In contrast, an attenuated response was observed for **2** for directing splicing of the mutant mini-genes, both WT I17T and DDPAC I17T which lack the A-bulge binding site; activity was only observed at the highest dose (5 µM) and the observed reduction was 15 ± 5% and 15 ± 2%, respectively (Figure 1E).

**Figure 3.**
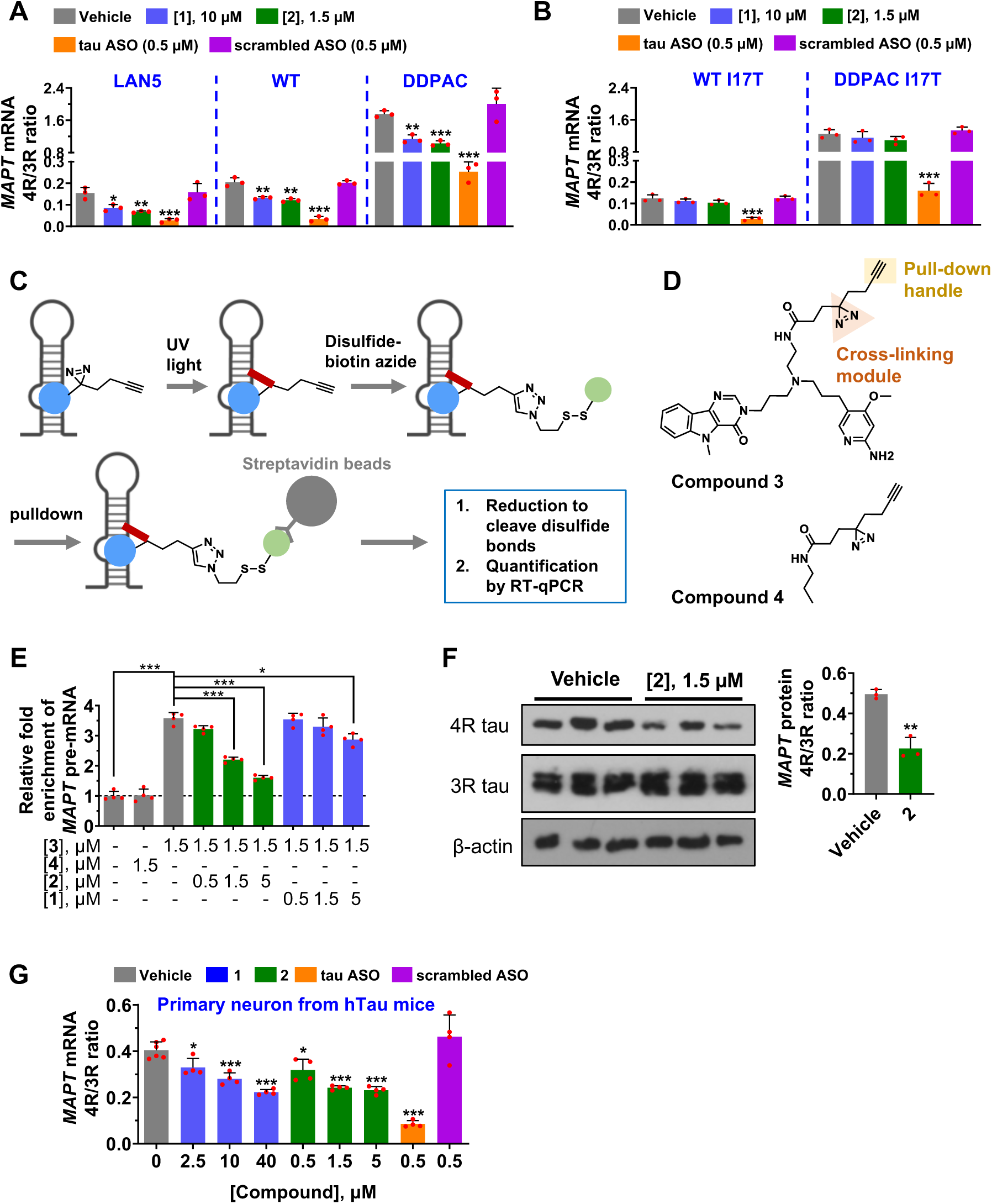
Target profiling in cells demonstrates direct target engagement of *MAPT* mRNA by 2 using Chemical Cross-Linking and Isolation by Pull-down (Chem-CLIP). (A) Effects of **1** and **2** on the ratio of 4R and 3R *MAPT* mRNA, as assessed by RT-qPCR in HeLa cells transfected with WT or DDPAC mini-genes, and in LAN5 neuroblastoma cells. Cells were treated with 10 µM of **1**, 1.5 µM of **2**, 0.5 µM of tau ASO, or 0.5 µM of a scrambled ASO control (n = 3 biological replicates). (B) Effects of **1** and **2** on the ratio of 4R and 3R *MAPT* mRNA, as assessed by RT-qPCR in HeLa cells transfected with mini-genes expressing the mutants WT I17T or DDPAC I17T, which abolishes the small molecule binding site. Cells were treated with 10 µM of **1**, 1.5 µM of **2**, 0.5 µM of the tau ASO, or 0.5 µM of a scrambled ASO control (n = 3 biological replicates). (C) Schematic of Chem-CLIP to cross-link *MAPT* pre-mRNA to small molecule binders in cells to validate target engagement. (D) Structure of probes used in Chem-CLIP and competitive Chem-CLIP (C-Chem-CLIP) studies. Chem-CLIP probe **3** comprises the RNA-binding module **2**, an alkyne purification module, and a cross-linking diazirine module. Control Chem-CLIP probe **4** lacks RNA-binding module **2**. (E) Target validation studies in LAN5 cells completed by Chem-CLIP using Chem-CLIP probe **3** and control probe **4** (n = 4 biolgical replicates). C-Chem-CLIP studies between **3** and increasing concentrations of **1** or **2**, which were preincubated with LAN5 cells prior to addition of **3**. (F) Effect of **2** (1.5 µM) on *MAPT* exon 10 alternative splicing in LAN5 cells at the protein level, as determined by Western blotting (n = 3 biological replicates). (G) Effects of **1**, **2**, tau ASO, or a scrambled ASO on *MAPT* exon 10 splicing outcomes in primary neurons harvested from htau mouse pups, as reported by the *MAPT* mRNA 4R/3R ratio (n = 6 biological replicates for vehicle; n = 4 biological replicates for treatment with compound or ASO). *, p < 0.05; **, p < 0.01; and ***, p < 0.001, as determined by one-way ANOVA. Error bars indicate SD.

To demonstrate that the observed reduction of firefly luciferase activity was due to facilitating exon 10 exclusion, RT-qPCR was employed to measure the amount of each isoform using primers specific for *MAPT* 3R and 4R mRNA. Both **1** (10 µM) and **2** (1.5 µM) reduced the 4R/3R ratio in HeLa cells transfected with WT or DDPAC mini-genes, as compared to vehicle (Figure 3A). In cells transfected with the WT mini-gene, the 4R/3R ratio was reduced from 0.20 ± 0.02 in vehicle-treated cells to 0.13 ± 0.01 (p = 0.004; 36 ± 2% reduction) and to 0.12 ± 0.01 (p = 0.003; 41 ± 3% reduction) in **1**- and **2**- treated cells, respectively. Likewise in cells transfected with the DDPAC mini-gene, the 4R/3R ratio was reduced from 1.8 ± 0.1 for vehicle to 1.1 ± 0.1 (p = 0.0012; 35 ± 5% reduction) and 1.0 ± 0.1 (p = 0.0003; 42 ± 3% reduction) for **1** and **2**, respectively. A very small and statistically insignificant change in the 4R/3R ratio was observed upon treatment of HeLa cells that expressed the WT I17T (from 0.12 ± 0.02 for vehicle to 0.11 ± 0.01 and 0.10 ± 0.01 for **1** and **2**, respectively) or the DDPAC I17T mutant (from 1.2 ± 0.1 for vehicle to 1.2 ± 0.2 and 1.1 ± 0.1 for **1** and **2**, respectively), both lacking the A-bulge binding site (Figure 3B). The percent spliced in index (PSI), defined as the ratio of the relative abundance of all isoforms containing a certain exon over the relative abundance of all isoforms of the gene containing the exon, was also calculated.^43^ PSI_10_ (the PSI of *MAPT* mRNA containing exon 10), calculated as 4R/(4R+3R) was reduced by **1** and **2**, as compared to vehicle, in HeLa cells transfected with WT or DDPAC mini-gene but not when transfected with WT I17T or DDPAC I17T mutant (Figure S7A-B).

### In vitro binding affinity of 1 and 2 for the *MAPT* SRE

The affinities of **1** and **2** for a model of the WT and DDPAC *MAPT* SRE were measured by monitoring the change in the inherent fluorescence of the small molecule as a function of RNA concentration. Both **1** and **2** bound to the WT RNA, with K_d_s of 3.3 ± 0.7 µM and 0.55 ± 0.11 µM, respectively, which were similar for a model of the DDPAC SRE (K_d_ = 3.6 ± 0.7 µM and 0.58 ± 0.1 µM for **1** and **2**, respectively) (Figure S8A). No saturable binding was observed to either the WT I17T or DDPAC I17T mutant, where the K_d_ is estimated to be >50 µM (Figure S8A). As the interaction of **2** with the *MAPT* SRE is ∼6-fold higher affinity than that for **1**, these data suggest that the 2-aminopyridine is forming favorable interactions with the RNA. To investigate whether the 2-aminopyridine module was indeed forming a base triple with the GC pair closing the A-bulge, the guanosine residue was replaced with 7-deaza-guanosine (Hoogsteen face; hydrogen bonds in a base triple; Figure S8B) in the WT model of the SRE. The affinity of **2** for the modified RNA decreased from 0.55 ± 0.11 to 3.4 ± 0.9 µM (Figure S8B). This replacement did not affect the binding affinity of **1**, in agreement with its limited interaction with the guanosine in the NMR solution structure (Figures 2C & S8B).

To verify target engagement in vitro, we employed Chemical Cross-Linking and Isolation by Pull-down (Chem-CLIP)^37–39^ (Figure 3C). In brief, **2** was appended with diazirine and alkyne functionalities to afford Chem-CLIP probe **3** (Figure 3D); the diazirine forms a cross-link upon irradiation with UV light while the alkyne is used as a chemical handle for pull-down. Compound **4,** which lacks the RNA-binding module, was used as a control Chem-CLIP probe (Figure 3D). We studied the ability of **3** to engage and pull-down radioactively labeled WT and DDPAC SRE RNA, affording similar IC_50_s, around 1 µM, in agreement with the similar affinity of **2** observed for the two RNAs (Figure S9A-B). The maximum percent pull-down observed for both RNAs was ∼40%, at the 10 and 50 µM concentrations (Figure S9A-B). For comparison, the Chem-CLIP probe derived from **1** has a previously reported IC_50_ around 10 µM.^17^ Notably, engagement of the two I17T mutants by **3** was not observed until a concentration of 50 µM (15 ± 0.9% for WT I17T and 9 ± 0.7% for DDPAC I17T; Figure S9A-B). Control Chem-CLIP probe **4** did not engage any of the RNAs tested as compared to vehicle. Chem-CLIP target validation can also be completed as a competition experiment between the Chem-CLIP probe and the parent compound (C-Chem-CLIP). C-Chem-CLIP experiments revealed that **2** competed the binding of **3** (1.5 µM) for WT and DDPAC SRE RNA similarly with IC_50_s of **∼**2 µM, indicating that **2** and **3** bind the same A-bulge binding site (Figure S9C).

### Compounds 1 and 2 reduce the binding of a model U1 snRNA by thermodynamically stabilizing the *MAPT* SRE in vitro

We previously reported an assay that assesses a compound’s ability to impede the binding of a model U1 snRNA to the *MAPT* SRE.^17^ A model of the DDPAC RNA hairpin was dually labeled at the 5’ and 3’ ends with fluorescein (FAM) and a black hole quencher (BHQ), respectively. This RNA can be unfolded by the addition of an oligonucleotide mimic of U1 snRNA, measured by an increase in FAM fluorescence as a function of time (Figure S10A). The addition of 10 µM of both **1** and **2** resulted in a significantly slower rate in the increase of FAM signal compared to vehicle (0.49 ± 0.03 min^-1^), where the rate of unfolding in the presence of **2** (0.37 ± 0.03 min^-1^; p = 0.002 compared to vehicle) was slower than that for **1** (0.41 ± 0.02 min^-1^; p = 0.009 compared to vehicle) (Figure S10B). Using an end-point measurement (5 min), we observed a dose dependent reduction in FAM fluorescence upon the addition of **1** or **2**, affording IC_50_s of 5.9 ± 1.7 µM and 0.9 ± 0.2 µM, respectively (Figure S10C). The results from both time- and dose-dependence studies reflect the higher affinity that **2** has for the *MAPT* SRE than **1** does.

The biochemical assays described above suggest that **1** and **2** thermodynamically stabilize the *MAPT* SRE RNA hairpin, which was verified using optical melting experiments of the DDPAC, WT, and DDPAC I17T model RNAs in the presence or absence of **1** (10 µM) or **2** (1.5 µM). Both compounds increased the melting temperature (T_m_) of the WT RNA (by 0.58 ± 0.06 °C for **1** and by 0.76 ± 0.1 °C for **2**) and DDPAC RNA (by 1.25 ± 0.06 °C for **1** and by 1.36 ± 0.11 °C for **2**) (Figure S11). Both compounds decreased ΔG°_37_ of the WT RNA (by -0.60 ± 0.26 kcal•mol^-1^ for **1** and by -1.07 ± 0.13 kcal•mol^-1^ for **2**) and DDPAC RNA (by -1.04 ± 0.15 kcal•mol^-1^ for **1** and by -1.59 ± 0.17 kcal•mol^-1^ for **2**) (Figure S11). Addition of either compound had no effect on the T_m_ or the ΔG°_37_ of the DDPAC I17T mutant, which lacks the small molecule binding site (Figure S11).

### Studying 2-*MAPT* SRE interactions by NMR spectroscopy and molecular modeling

As further evidence that **2** engages the *MAPT* DDPAC SRE in vitro, various NMR spectral studies were carried out, including WaterLOGSY^44^ (water Ligand Observed via Gradient SpectroscopY) and 1D imino proton NMR spectra. In WaterLOGSY studies with a duplex model of the WT SRE (Figure S12A), addition of the RNA changed the sign of the nuclear Overhauser effect (NOE), resulting in the appearance of negative peaks (Figure S12B-C). In complementary studies, addition of **2** to the RNA revealed dose dependent changes in the RNA’s 1D imino proton spectra. Imino proton resonances for G4 and U18, which form a G·U base pair nearby the A-bulge, and G20 shifted up-field upon the addition of 0.5 equivalents of **2**. U18 completely disappeared upon addition of 1.5 – 2.0 equivalents of **2** while G4 broadened from 0.5 – 2 equivalents. Resonances for G7 and G17, which form the GC base pairs closing the A-bulge, and G21 were broadened and disappeared as 0.5 – 1.0 equivalents of **2** were added. These data suggest that the complex, particularly the GC closing pair, is dynamic, and these observations are similar to those for the complex with **1**.^17^ Resonances for G3 (slightly), U8 (significantly), G9 [slightly and begins to merge with G7 (significantly) and G20 (significantly)], and U10 (slightly) were also broadened, but otherwise remain unchanged.

In addition, 2D ^1^H-^1^H D_2_O NMR spectra were acquired to assign nonexchangeable protons. In a 400 ms mixing time NOESY spectrum at 35 °C (Figure S13), a sequential H6/H8 to H1′ NOESY walk was assigned through all residues except for G20, which could not be assigned due to spectral overlap. A single NOE was observed from H5/H6 of **2** to G17H1′ (forms one of the A-bulge’s closing base pairs), which orients the benzene ring within the A-bulge site. Overall, the NOE patterns of the RNA and compound were similar to those observed in 2D NOESY spectra of the tau WT RNA with **1** (previously reported),^17^ but with broader signals likely due to intermediate exchange associated with higher ligand-RNA affinity or complex solubility.

To better understand the orientation adopted by **2** within the binding pocket provided by the A-bulge and a more detailed structural analysis, **2** was docked into the previously elucidated NMR structure of the *MAPT* WT RNA in complex with **1**; **1** was removed to create an apo RNA. Before studying the **2**-RNA complex, we first verified that AutoDock 4.0 (GPU accelerated version) could recapitulate the bound structure of **1**. The lowest energy bound state predicted by docking **1** into the *MAPT* WT RNA overlapped with the solution structure with an RMSD of 0.535 Å. AutoDock-GPU was then used for untargeted and unrestrained docking of **2** against the RNA. The resulting lowest energy state of **2** bound to the RNA yielded an orientation in accordance with the single NOE observed between H5/H6 of the compound to G17H1′ (Figure S13). A closer inspection of the bound structure revealed that the interaction between **2** and the RNA was stabilized by a combination of stacking interactions with the A-bulge’s closing base pairs, multiple hydrogen bonds (with bases as well as the phosphodiester backbone), and hydrophobic interactions, affording a binding energy of -9.91 kcal/mol (Table S1). This structure was then used to perform molecular dynamics (MD) simulations. The resultant clustering of the obtained trajectory and free energy calculations (Table S2) showed that the 2-aminopyridine group forms a base triple with the G7-C16 pair via two hydrogen bonds with the G7 base in the major groove (Figure 2C).

The topology of the binding site when **1** and **2** were bound to the WT *MAPT* SRE was compared using principles moment of inertia (PMI). In brief, PMI serves as a descriptor to assess the shape of a given molecule, whether rod, disc, or sphere. These calculations showed that the two topologies adopted by **1** and **2** are different, where the binding pocket of **2** adopted a rod-shape topology while **1** adopted a sphere-like topology, providing clues for molecular recognition and possible lead optimization (Figure S14). Similar to **1**, the ellipticine core of **2** displaced the A-bulge from the co-axial helical axis and stacked on G7 (one of the A-bulge’s closing base pairs) and the closing base pair formed by C5-G17A. Although **1** stacked on C5-G17 base pair, it did not form stacking interactions with G7. The H5/H6 proton of **2** laid in the A-bulge, consistent with its observed NOE with G17H1′. Furthermore, the 2-aminopyridine group in **2** formed two hydrogen bonds with the backbone and a hydrogen bond with C14. These additional interactions stabilized the **2**-RNA complex more so than the **1**-RNA complex. These differences are reflected in the binding energies of **1** and **2**, -8.91 kcal/mol and -9.91 kcal/mol, respectively. Unsurprisingly, **2** occupied a larger surface area within the binding pocket than **1** (∼723.8 Å^2^ vs. ∼330.1 Å^2^), also contributing to the stronger interaction of **2** with the *MAPT* RNA.

### Compound 2 inhibits endogenous *MAPT* exon 10 inclusion and generation of 4R tau protein in cells

The biophysical, biochemical, and cellular reporter data suggest that **2** may bind the endogenous *MAPT* SRE and direct splicing away from the 4R isoform. We previously studied the effect of **1** on the alternative splicing of exon 10 at the RNA and protein levels in the neuroblastoma cell line LAN5. At a 10 µM dose, **1** reduced the *MAPT* mRNA 4R/3R ratio from 0.15 ± 0.03 for vehicle to 0.09 ± 0.02 (p = 0.02; 44 ± 8% reduction) and protein isoforms from 0.50 ± 0.02 for vehicle to 0.32 ± 0.02 (p = 0.008; 37 ± 4% reduction).^17^ Here, **2** (1.5 µM) facilitated exon 10 exclusion and reduced the 4R/3R mRNA ratio to 0.07 ± 0.004 (p = 0.005; 56 ± 4% reduction), as determined by RT-qPCR (Figure 3A), and the 4R/3R protein ratio to 0.23 ± 0.05 (p = 0.002; 54 ± 9% reduction), as determined by Western blotting (Figure 3F). For comparison, treatment of LAN5 with the tau ASO (0.5 µM) reduced the 4R/3R *MAPT* mRNA ratio from 0.16 ± 0.04 for a scrambled ASO to 0.03 ± 0.01 (p = 0.006) (Figure 3A).

### Compound 2 binds endogenous *MAPT* pre-mRNA in LAN5 cells

In vitro Chem-CLIP studies demonstrated that **2** bound the *MAPT* SRE (Figures S9). We therefore studied whether **2** binds the SRE in the context of native, endogenous *MAPT* in LAN5 neuroblastoma cells. A 3.6 ± 0.2-fold enrichment of *MAPT* pre-mRNA was observed upon treatment with 1.5 µM of **3**; no enrichment was observed upon treatment with **4** (Figure 3E). Further, C-Chem-CLIP studies revealed that the cross-linking and hence binding of **3**, was competed by **1** or **2** in dose dependent fashion (Figure 3E). Notably, the concentrations of **1** required to compete **3** were greater than those required by **2**; 5 µM of **1** competed **3** to a lesser extent than 1.5 µM of **2** competed.

### Mapping the binding site of 2 in LAN5 cells using Chem-CLIP-Map-Seq

The precise binding site of **2** within *MAPT* pre-mRNA in LAN5 cells was mapped using a method named Chem-CLIP to Map Small Molecule-RNA Binding Sites (Chem-CLIP- Map).^45^ Cross-linking impedes the processivity of reverse transcriptase, resulting in an “RT stop”. Thus, the nucleotide where truncation of the complementary (c)DNA occurs indicates the binding and cross-linking site. After the pull-down of *MAPT* RNA by **3** in LAN5 cells, reverse transcription was completed with a gene-specific primer followed by ligation of single stranded (ss)DNA adaptors and cloning (Figure S15A). Sequencing analysis showed that cross-linking occurred to the G in the 5’ CG closing base pair of the A-bulge (Figure S15B). These results are consistent with the formation of base triple as the cross-linking module is positioned on the opposite strand from where 2-aminopyridine forms a base triple (Figures 2C and S15B).

### Effect of 2 transcriptome-wide in LAN5 cells

To study the effect of **2** transcriptome-wide, RNA-seq analysis was completed. Differentially expressed genes were identified using quantification and analyses from the Kallisto-Sleuth packages in R.^46^ For gene abundance, genes having an absolute fold change >2 with an adjusted P value < 0.05 were considered to be significantly altered in expression. Among the 15,404 genes commonly detected in all samples, very few changes were observed upon **2**-treatment (15,362/15,404 (99.73%) genes were unchanged 99.73%) or tau ASO treatment (15,399/15,404 (99.97%) genes were unchanged) (Dataset S1 & Figure S16). Notably, neither **2** nor the tau ASO altered the expression of *MAPT* gene (Dataset S1 & Figure S16).

### Compound 2 directs exon 10 alternative splicing in primary neurons from htau mice

We next evaluated **2** in cultured primary neurons extracted from the cortex of htau transgenic mice. The humanized tau mice lack the endogenous murine *MAPT* gene and instead express all six isoforms (including both 3R and 4R forms) of human *MAPT*,^47^ and exhibit age-dependent impairment of cognitive and synaptic function.^48^ Brains from htau mice pups were isolated, and the neurons from the cortex were collected and cultured. These neurons require 15 days to express 3R and 4R *MAPT* RNA at measurable levels.^17^ Therefore, after 15 days, the neurons were treated with **1**, **2**, tau ASO, or scrambled ASO^10^ for 48 h. The compounds were well tolerated at the doses studied (up to 40 μM of **1** and 5 μM of **2**), with no significant changes in morphology or cell density observed upon treatment. The *MAPT* mRNA 4R/3R ratio was measured by RT-qPCR, revealing dose dependent reduction of the 4R/3R ratio as well as PSI_10_ by the tau ASO, **1** and **2**, where **2** was more potent than **1** (Figures 3G & S7C).

### Drug metabolism and pharmacokinetic analysis of 2

To determine whether **2** can direct *MAPT* alternative splicing in vivo, in vitro and in vivo drug metabolism and pharmacokinetics (DMPK) studies were completed. We first evaluated the brain exposure of **2** upon a single dose of 100 mg/kg delivered orally (*p.o.*) by measuring the concentration of the small molecule in both brain and plasma (Figure S17A). At the 2 h time point, the concentrations of **2** in brain and plasma were 7.0 ± 1.1 µM and 5.4 ± 2.7 µM, respectively. The differences in concentration in the brain and plasma increase over time: 3.6-fold difference at 12 h, 4.3-fold difference at 24 h, and 5.8-fold at 48 h. At 48 h post-administration, the concentrations of **2** were 2.9 ± 1.6 µM in brain and 0.5 ± 0.3 µM in plasma (Figure S17A). These concentrations are in the range where they directed *MAPT* exon 10 splicing in primary neurons isolated from htau mice (ex vivo studies). [For comparison, the percent free for Ritonavir (a protease inhibitor used for the treatment of HIV) and Carbamazepine (an oral drug used to treat seizures) was 0.18 ± 0.02% and 37.7 ± 4.2%, respectively.]

Because of the observations in the brain exposure studies above, additional in vitro and in vivo PK studies were carried out. Drug distribution in the brain is often evaluated using a simple readout of brain concentration vs. plasma concentration and reporting the partition coefficient (K_p_). Because this uses total drug instead of free drug, which represents the portion of the drug able to diffuse into cells and modulate the target, a more rigorous investigation converts to free plasma and brain concentrations to calculate K_p,uu_.^49^ Typically, K_p,uu_ values approximate 1 when passive diffusion drives the equilibration between the brain and plasma. Deviation often predicts transporter involvement. Thus, the K_p,uu_ and other parameters were measured for **2**.

The pharmacokinetics of **2** were evaluated under single dose conditions as an intravenous dose at 5 mg/kg and by oral gavage at 100 mg/kg (Figure S17A). Compound **2** had a long half-life of approximately 16 h and a C_max_ of 1.9 µM after the 100 mg/kg oral dose. Because the efficacy studies were completed using repeat dosing conditions (*vide infra*), the oral PK study was repeated to determine concentrations under steady-state dosing conditions. C_max_, 3.5 µM, and total drug exposure (AUC) were approximately doubled with daily oral dosing of **2** at 100 mg/kg.

The free fraction in both plasma and brain were evaluated using equilibrium dialysis and were found to be 8.9 ± 0.9% and 4.4 ± 0.3%, respectively. These data were used in concert with plasma and brain concentration data from mice orally dosed with 100 mg/kg of **2**. The equilibrium dialysis data were used to calculate both total and free concentrations in brain and plasma (Figure S17A). While not tested, comparison of single dose and daily dosing at 100 mg/kg doubled plasma C_max_ and AUC, and a similar doubling of brain concentration is anticipated. The total brain-to-plasma ratio (K_p_) was 4.0 ± 1.8, indicating significant brain penetration. When adjusted for non-specific binding, the free brain-to-plasma partition coefficient (K_p,uu_) varied across samples, with an average of 2.7 ± 1.2. This may indicate an additional factor influencing drug uptake into the brain. Alternatively, this discrepancy may reflect compound binding to mRNA in the brain, shifting equilibrium and leading to higher in vivo brain concentrations that were not recapitulated in the equilibrium dialysis experiment that uses brain homogenate and likely has degraded RNA.

The determined K_p,uu_ suggests that beyond the normal factors such as non-specific lipid interactions, additional interactions, such as target binding, may increase the intracellular drug concentrations. This discrepancy underscores the importance of intracellular binding components in CNS pharmacokinetics and highlights the limitations of equilibrium dialysis in fully capturing equilibrium. The RNA that binds compound **2** and promotes its retention within the cell are quickly degraded upon brain homogenization, which impacts binding predictions.

### Compound 2 directs exon 10 alternative splicing and alleviates aberrant behavioral phenotypes in htau mice

Oral administration of 100 mg/kg of **2** *quaque altera die* (q.o.d.) to htau mice was well tolerated as no change in the weight of WT or htau mice was observed throughout the 20-day treatment period (Figure S17). After the treatment period, the abundance of 3R and 4R MAPT isoforms was assessed by RT-qPCR, where the 4R/3R ratio was reduced from 0.91 ± 0.17 for vehicle to 0.56 ± 0.05 (p = 0.007; 38 ± 5% reduction) upon **2**-treatment, corresponding to a reduction of PSI_10_ from 47 ± 4% to 36 ± 2% (p = 0.007) (Figure 4A). Total human *MAPT* transcript levels were unaffected (Figure S18).

**Figure 4.**
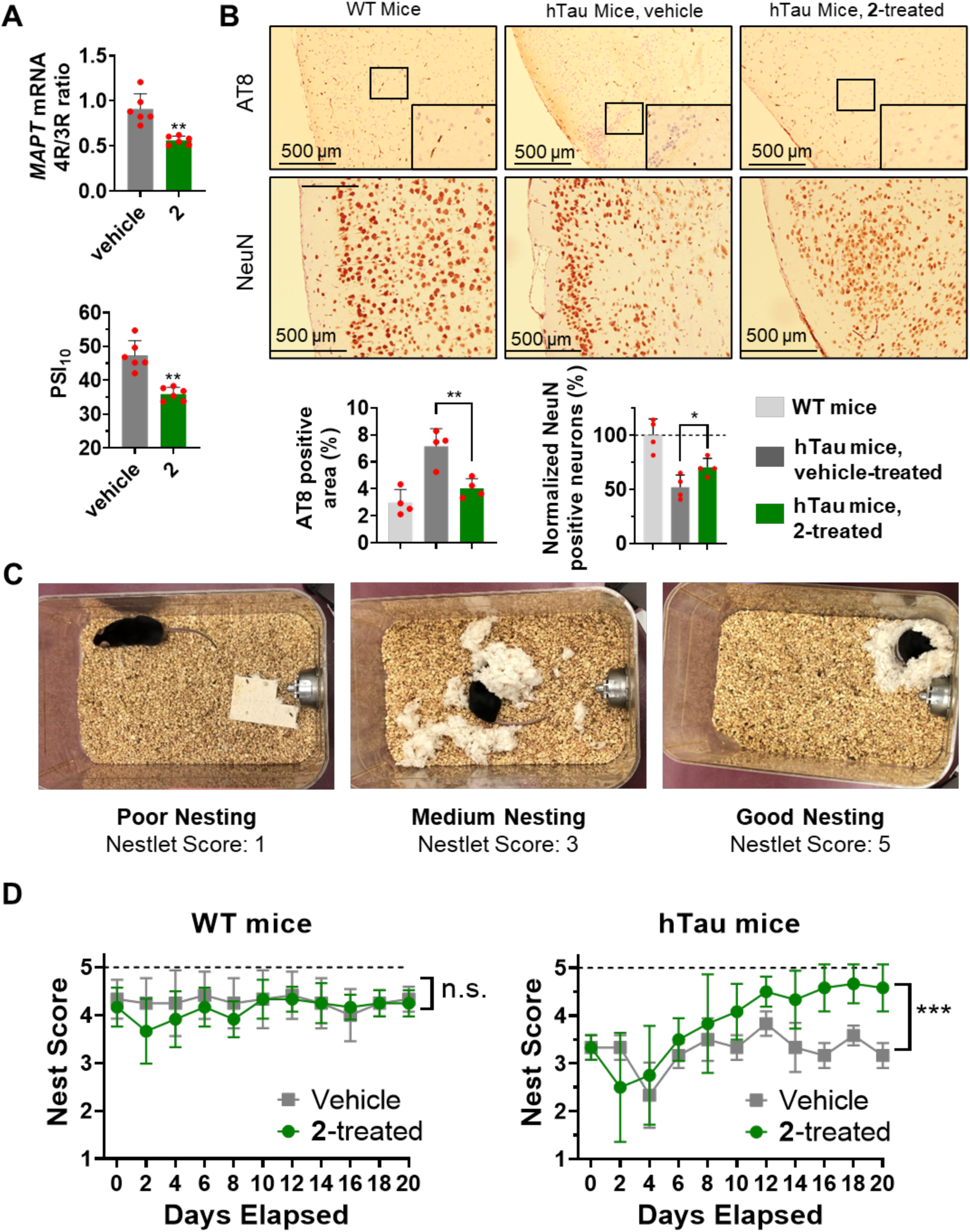
Compound 2 mitigates disease-associated pathologies and rescues aberrant behavioral phenotypes in htau mice when orally administrated at 100 mg/kg q.o.d. (A) Effect of **2** on the *MAPT* mRNA 4R/3R ratio (top) or PSI_10_ (PSI of *MAPT* exon 10, bottom) in htau mice, as compared to vehicle-treated mice (measured by RT-qPCR; n = 6 biological replicates, each from a different mouse). (B) Top, representative histological images of cortex from htau mice treated with vehicle or **2** and quantification thereof. Immunohistochemical staining was used to measure levels of phosphorylated tau (AT8 antibody; maker of tauopathy) and NeuN (marker of neuron viability) (n = 4 biological replicates, each from a different mouse). Bottom, quantification of AT8 positive area and NeuN positive neurons. (C) Representative images of nests built by htau mice. Nesting activity was assessed using scoring criteria to rate the quality of nest construction and the amount of torn nestlet material on a scale of 1 – 5 as previously described.^53^ (D) Nesting scores upon treatment of WT or htau mice treated with vehicle or **2** (n = 6). *, p < 0.05; **, p < 0.01; and ***, p < 0.001, as determined by one-way ANOVA. Error bars indicate SD.

Aberrant tau protein phosphorylation is frequently associated with tauopathies caused by an imbalanced 4R/3R ratio^50, 51^ and is also observed in the htau mouse model.^51^ Thus, sagittal brain sections were assessed for tau phosphorylation by immunohistochemistry (IHC) using AT8 antibody, which detects S202/T205 phosphorylation events.^50^ Treatment of **2** reduced antibody immunoreactivity throughout the cortex, indicative of decreased pathological tau burden with a lower 4R/3R ratio (Figure 4B). NeuN, a neuron-specific protein, was used as a marker to quantify neuron viability, which was enhanced in **2**-treated mice (Figure 4B).

One of behavioral deficits observed in htau mice is impaired nesting behavior,^51, 52^ which can be quantified based on published criteria (Figure 4C).^53^ From Day 1 to Day 20, mice were individually housed and nesting material from the previous day was removed. An intact 3.0 g nestlet was placed within each cage. Approximately 15 h later, the resulting nest was photographed for classification, and the untorn nestlet, if present, was weighed. Nesting is scored on a rating scale of 1–5^53^ where if the nestlet is over 90% intact, it is given a nesting score of 1 (Poor; Figure 4C). When the nestlet is partially torn but >50% remains, the nesting score is 2; a score of 3 is assigned when 50-90% of the nestlet is torn but spread around the cage and no identifiable nest site is found (Medium; Figure 4C). When more than 90% of the nestlet is torn and the nest is identifiable but flat, the nesting score is 4. When a near perfect nest is built and the wall is higher than the mouse’s body, it is assigned a nesting score of 5 (Good; Figure 4C).

WT mice showed no significant difference in nesting behavior without or with **2**-treatment after 20 days, and the average nesting score was consistent over time. On average, the nesting score at Day 20 for untreated and **2**-treated WT mice was 4.3 ± 0.1 and 4.1 ± 0.2, respectively (Figure 4D). The average nesting score for untreated htau mice was consistently below 3.8 and was 3.2 ± 0.2 after 20 days. In contrast, the nesting scores for **2**-treated mice improved over the treatment period, on average 2.6 ± 1.1 at Day 2 and 4.5 ± 0.4 at Day 20 (Figure 4D). Collectively, these in vivo studies demonstrated that **2** is orally bioavailable, well tolerated, and able to mitigate cellular and behavioral phenotypes observed in htau mice.

## DISCUSSION

Aberrant alternative splicing causes a wide range of diseases, from β-thalassemia^54^ to Duchenne muscular dystrophy^55^ to Myelodysplastic syndrome.^56, 57^ The notion that alterative splicing could be directed for therapeutic benefit was first demonstrated using oligonucleotides^58, 59^ and later small molecules.^60^ Indeed, two have been approved for the treatment of spinal muscular atrophy (SMA), nusinersen (Spinraza)^61, 62^ and risdiplam.^63–65^ Interestingly, risdiplam functions as a molecular glue to stabilize an RNA helix formed by the 5’ splice site of SMN2 exon 7 and the 5′ terminus of U1 snRNA/U1 snRNP to facilitate exon *inclusion*.^63–65^ The results collected herein suggest that **2** has a simpler mode of action where it binds the *MAPT* SRE and thermally stabilizes its structure to impede U1 snRNA binding and facilitate exon *exclusion*. Thus, the two approaches for directing alternative splicing toward inclusion or exclusion could be complementary.

The design principles outlined here emphasize the importance of shape complementarity between the small molecule and RNA binding pocket, beyond topological properties, as a means to increase the favorable interactions, and hence affinity, selectivity, and potency. RNA-binding pockets are quite different than protein-binding pockets and thus different design strategies must be developed and implemented.^60^ The molecule designed herein exploits spatial proximity to extend the binding pocket beyond what is typically perceived as the boundaries of said pocket. We have shown that T-shaped small molecules, grown vertically, can have sub-micromolar binding affinity. This T-shaped geomorphic compound design principles could be a general strategy to generate small molecules targeting RNA with favorable physicochemical properties, bio-distribution, and oral bioavailability as exemplified in this study. Interestingly, PMI calculations show that the majority of compounds in DrugBank are rod-like molecules with fewer disc-like or sphere-like molecules (Figure S19), in agreement with a previous report.^66^ Very few T-shaped molecules are observed, which is a restraint imposed by the binding pocket topology of proteins that prevents longitudinal growth. This suggests that a T-shape topology could be a strategy for targeting RNAs specifically over proteins.

Targeting RNAs using a base triple formation strategy is an area of ongoing investigation in the peptide-nucleic acid space.^23, 67–70^ Emerging datasets described here could provide additional modules that could be appended to small molecule RNA binders to increase affinity and specificity. We envision this base triple formation strategy could be applied to optimize compounds that target other RNAs that cause diseases, including those by aberrant pre-mRNA splicing. Notably, the distance between the binding pocket and a base pair that can form a base triple will be different for each target. Further, most targets will lack available structural data, such as the NMR solution structures used herein, to guide design. Although a 2-aminopyridine base triple-forming module was successfully employed, the module could be optimized to increase its affinity and specificity while additional modules could be developed for AU or GU base pairs.

## Acknowledgements

The authors thank Xiangming Kong (The Herbert Wertheim UF Scripps Institute for Biomedical Innovation and Technology) for assistance with NMR experiments and Tanya Khan for help with mouse studies. The authors also acknowledge University of Florida Research Computing for providing computational resources and support that have contributed to the research results reported in this publication (http://www.rc.ufl.edu) and The Scripps Research Institute (now UF Scripps) Histology Core and Animal Behavior Core.

## Author contributions

Conceptualization: M.D.D. Methodology: M.D.D., J.L.C.-D., P.Z. Investigation: P.Z., Y.T., A.T., J.T.B., Y.A., H.A., M.A., J.L.C. Formal Analysis: P.Z., Y.T., A.T., J.T.B., Y.A., H.A., M.A., J.L.C. Visualization: P.Z., Y.T., A.T. Supervision: M.D.D. Writing – original draft: P.Z., A.T., J.L.C.-D.

## Data and materials availability

All data associated with this study are present in the paper or the Supplementary Materials.

## ToC Graphic

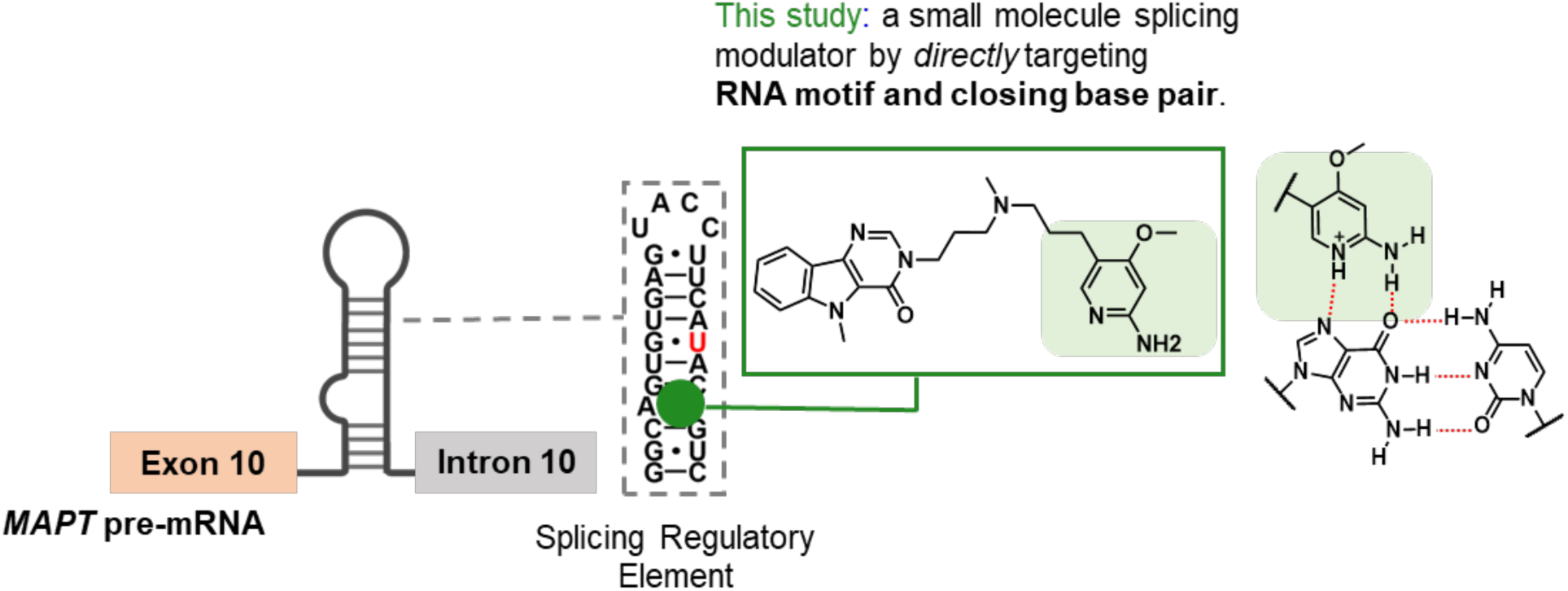

