## Supplementary Information for "Design of an orally bioavailable small molecule that modulates the microtubule-associated protein tau’s pre-mRNA splicing"

**This PDF file includes:**

Materials and Methods
Supplementary Figures S1 – S19
Tables S1 – S3
Synthetic Experimental Procedures
Compound Characterization

### Materials and Methods

**General Nucleic Acids Methods.** All DNA oligonucleotides used in these studies were purchased from Integrated DNA Technologies, Inc. (IDT). When received, they were dissolved in nanopure water and used without further purification. A Vivo-Morpholino antisense oligonucleotide (ASO) targeting the *MAPT* exon-10-intron junction hairpin (tau ASO: 5'-TGAAGGTACTCACACTGCCGC-3') and a scrambled ASO (5'-CTGTCTGACGTTCTTTGT-3') were purchased from Gene Tools, LLC. The ASO was dissolved in water to afford a final concentration of 1 mM and stored at 4 °C as recommended by the manufacturer.

All RNA constructs were purchased from GE Healthcare Dharmacon, Inc. When received, they were deprotected according to the manufacturer's recommended protocol and desalted with PD-10 columns (GE Healthcare), also per the manufacturer's recommended protocol. RNA oligonucleotide concentrations were determined by their absorbance at 260 nm at 90 °C using a Beckman Coulter DU800 UV/vis spectrophotometer and the corresponding extinction coefficient provided by the manufacturer.

**Compounds.** All compounds were synthesized as described below in **Synthetic Experimental Procedures**.

**Preparation of NMR Samples.** Two duplex-forming oligoribonucleotides r(5′-CCGGCAGUGUG-3′) + r(5′CACACGUCGG-3′) were designed to mimic the stem of the tau wild type (WT) hairpin. Individual RNA oligonucleotides were combined in equimolar amounts in 400 μL of NMR Buffer [10 mM $KH_2PO_4$/$K_2HPO_4$, pH 6.0 and 0.5 mM EDTA] prepared in $H_2O$ or 100% $D_2O$. The final RNA duplex concentrations were 15 μM for WaterLOGSY experiments, 100 μM for 1D $^1$400 μM for 2D $^1D_2O$ was added to 5% (v/v) to provide a lock signal. RNA samples were annealed by heating to 95 °C for 3 min, followed by slow cooling to room temperature before being added to Shigemi NMR tubes (Shigemi, Inc.). Compound of interest was added to a final compound:RNA molar ratio of 20.0 for WaterLOGSY experiments, 2.0 for 1D $^1H$ $H_2O$ experiments, and 1.5 for 2D $^1$NOESY experiments.

**NMR Spectroscopy.** NMR spectra were acquired on Bruker Avance III 600 and 700 MHz spectrometers equipped with cryoprobes. WaterLOGSY spectra were acquired at 298 K; 1D NMR spectra of samples in 95% $H_2O$/5% $D_2O$ were acquired at 283 K; and 2D NOESY spectra in 100% $D_2O$ were acquired at 308 K. For 1D spectra of samples in 95% $H_2O$/5% $D_2O$, an excitation sculpting sequence during acquisition suppressed the water signal. Proton chemical shifts were referenced internally to the frequency of water. WaterLOGSY and 1D $^1H$ spectra were processed using Bruker TopSpin and 2D NMR spectra were processed with NMRPipe[1] and assigned with SPARKY[2].

**Simulation Protocols.** Simulations were carried out with the AMBER 18 simulation package[3] using the PARM99 force field[4] with revised χ[5] and α/γ[6] torsional parameters. Each system was first neutralized with $Na^+$ ions,[7] which then was solvated with TIP3P[8] water molecules in a truncated octahedral box with periodic boundary conditions extended to 10 Å using the LEAP module of Amber 16[4]. Each system was then added with 13 $Na^+$ and 13 $Cl^-$ ions to mimic physiological conditions, where after equilibration each system had a $Na^+$ concentration of 0.16 M. The number of $Na^+$ and $Cl^-$ ions were chosen as an approximate calculation based on the number of water molecules in the simulation box and the box size. The structures were minimized with the sander module[3] in two steps. A positional restraint (10 kcal $mol^{-1}$ Å$^{-2}$) was applied to the tau RNA in the first step of minimization with 5000 steps of the steepest-descent algorithm and subsequently followed with the second round of minimization with 5000 steps of the conjugate-gradient algorithm and no restraints.

Minimization was followed by an equilibration protocol first in constant volume with restraints on the RNA molecule (10 kcal $mol^{-1}$ $Å^{-2}$) and gradually increasing the temperature up to 300 K for several nanoseconds using the Langevin thermostat.[9] A second round of equilibration was performed at constant pressure (1 atm) and constant temperature (300 K), and pressure coupling[10] of 1.0 $ps^{-1}$, gradually removing the constraints on the solute. After minimization and equilibration, a 3 µsec MD simulation with a 2 ps time step was performed using NPT (constant temperature and constant pressure) dynamics with isotropic positional scaling. The reference pressure was set to 1 atm with a pressure relaxation time of 2 ps. SHAKE[11] was turned on for constraining bonds involving hydrogen atoms. An atom-based long-range cutoff of 8.0 Å was used in the production run. The reference temperature was set to 300 K. Particle Mesh Ewald (PME) was used to handle the electrostatics[12] and the Langevin thermostat[13] was applied with a coupling constant $\gamma$ = 1.0 $ps^{-1}$. Simulations were performed using the PMEMD.CUDA implementation of AMBER18.[12]

**Parametrization of compound 2.** A hand-built model of **2** was geometry-optimized at the HF/6-31G* quantum mechanical (QM) level consistent with AMBER ff10 force fields. Atomic charges were determined using the restrained electrostatic potential (RESP)[14] charge fitting. Bonds, torsions, angles, improper torsions, and Lennard-Jones parameters were assigned using the general Amber force field[15] (GAFF) using the Antechamber programs.[16] QM calculations were performed using Gaussian 09.[17]

**Molecular Docking.**

**Docking**. A previously published NMR solution structure (PDB:6VA3) of the tau RNA in complex with a small molecule was used for docking studies after removal of the small molecule.[18] Prior to grid calculations, polar hydrogen atoms and Gasteiger charges were added to RNA using AutoDock Tools 1.5.6 and MG Tools of AutoDock Vina[19] and saved in pdbqt format. The 3D structures of small molecules were created with OBabel[20] from the corresponding SMILES input file and geometry-optimized with general AMBER force field (GAFF)[16] in 5000 cycles prior to further processing for docking. Polar hydrogen atoms and Gasteiger[21] charges (partial charges of each atom) were then added to the small molecules as described above. The protonation state of the compounds was calculated at the physiological pH (7.4) prior to docking with OBabel. The Grid file was generated from ligand and receptor pdbqt files, applying prepare_gpf4.py, autogrid4 and prepare_dpf4.py script to prepare the docking parameter file. AUTODOCK-GPU[22] was then used to dock the ligands against the receptor. Twenty different structures were computed with AutoDock-GPU for each molecule using the Solis Wets[23] method of search.

**Calculation of accessible surface area (ASA).** dr-sasa was used to calculate solvent accessible surface area.[24]

**Calculation of interactions.** Interactions between RNA and compounds were extracted with fingeRNAt.[25]

**Calculation of CNS-MPO.** Calculator Plugins from Marvin were used to calculate the CNS-MPO score. ChemAxon (http://www.chemaxon.com).

**3D-RISM.** 3D-RISM (three-dimensional reference interaction site model) provides the 3D map of the density distribution of the solvent around the solute.[26-27] The probability density $\rho_\gamma g_\gamma(r)$ of finding the interaction sites (γ) of solvent molecules positioned around the solute, in the 3D space (r) was used to represent the solvent structure. 3D-RISM calculations were performed using the AMBER18 implementation applying the Kovalenko−Hirata (KH) closure[28] to calculate the binding free energy along with MMGB(PB)SA.[29]

**Binding Affinity Measurements.** Binding assays were completed as previously described.[30] Briefly, the RNA of interest was folded in 1× Assay Buffer-1 (AB1: 8 mM $Na_2HPO_4$, pH 7.0, 185 mM NaCl, and 1 mM EDTA) by heating at 95 °C for 30 s followed by slowly cooling to room temperature. Bovine serum albumin (BSA) was then added to a final concentration of 40 μg/mL to afford 1× Assay Buffer-2 (AB2: 8 mM $Na_2HPO_4$, pH 7.0, 185 mM NaCl, 1 mM EDTA, and 40 μg/mL BSA). The RNA solution was serial diluted with 1× AB2. Then, compound of interest was added to each RNA solution to a final concentration of 0.5 μM. The samples were incubated at room temperature in the dark for 15 min and then transferred to a black 384-well plate (Greiner Low-Volume 784076). Fluorescence intensity was measured using a Tecan Plate Reader (Gain: 100, Integration time: 40 μs). The excitation wavelength and emission wavelengths for compound **1** are 335 nm/430 nm, respectively, and for compound **2** are 330 nm/390 nm, respectively. The change in fluorescence of the small molecule was calculated by its comparison to samples lacking RNA. Binding affinity was calculated using Equation 1, as previously described:[30]

$$\mathrm{I} = \mathrm{I_0} + 0.5\Delta\varepsilon\{([\mathrm{SM}]_0 + [\mathrm{RNA}]_0 + \mathrm{K_d}) - (([\mathrm{SM}]_0 + [\mathrm{RNA}]_0 + \mathrm{K_d})^2 - 4[\mathrm{SM}]_0[\mathrm{RNA}]_0)^{0.5}\} \qquad \textbf{(Eq.1)}$$

where $\mathrm{I}$ and $\mathrm{I_0}$ are the measured fluorescence intensity of the compound with or without RNA, respectively, Δε is the difference between the fluorescence intensity with the infinite concentration of RNA and the fluorescence intensity without RNA, $[SM]_0$ and $[RNA]_0$ are the concentrations of the small molecule (SM) and RNA, respectively, and $K_d$ is the dissociation constant.

**U1 snRNA Mimic Quenching Assay.** Two different types of experiments were used to explore the ability of small molecules to inhibit U1 snRNA binding: (i) time course studies completed at a single dose; and (ii) concentration dependent studies completed as an end-point measurement.

*Time course studies*. A 200 nM aliquot of a dually labeled model of the DDPAC RNA (5'-FAM-GGCAGUGUGAGUACCUUCAUACGUC-BHQ-1-3'; where FAM is 5(6)-Carboxyfluorescein and BHQ-1 is black hole quencher) was folded in 1× Assay Buffer as described in **Fluorescent Binding Affinity Measurements.** Compound **1** or **2** was added to a final concentration of 10 μM, and the samples were incubated at room temperature in the dark for 15 min. A U1 snRNA mimic with a sequence of 5′-GUCCACUCAUGGA-3′ was then added to the samples to a final concentration of 20 μM. The fluorescence intensity of FAM was measured as a function of time for 1 h using a Biomek FLx800 plate reader, with excitation and emission wavelengths of 485 nm and 525 nm, respectively. The kinetic rate was calculated using Equation 2, as previously described.[18]

$$[\mathrm{Y}] = 0.5 \times [b - \mathrm{K_0} - [\mathrm{X}] + \sqrt{([\mathrm{X}] + \mathrm{K_0} - b)^2 + 4b \times \mathrm{K_0}}\,] \qquad \textbf{(Eq. 2)}$$

where [Y] is the fold change of fluorescence intensity of FAM, [X] is the time in min, $K_0$ is the kinetic rate to be fitted by the equation, and b is the maximal possible fold change of fluorescence intensity with infinite time.

*Concentration dependent studies (end-point measurement).* A 200 nM aliquot of a dually labeled model of the DDPAC RNA (5'-FAM-GGCAGUGUGAGUACCUUCAUACGUC-BHQ-1-3') was folded in 1× Assay Buffer as described in **Fluorescent Binding Affinity Measurements.**

Compound **1** or **2** was added at the indicated concentration (**1** from 200 µM to 0.1 µM and **2** from 50 µM to 0.1 µM), and the samples were incubated at room temperature in the dark for 15 min. The highest concentration of compound **2** in this assay was 50 µM based on its solubility in 1× Assay Buffer. The U1 snRNA mimic was then added to the samples to a final concentration of 20 µM. After 5 min of incubation with U1 snRNA mimic at room temperature, the fluorescence intensity of FAM was measured using a Biomek FLx800 plate reader, with excitation and emission wavelengths of 485 and 525 nm, respectively. $IC_{50}$s were calculated using Equation 3:

$$\mathrm{I} = \mathrm{I}_0 + 0.5 \times \mathrm{b}\,\{([\mathrm{SM}]_0 + [\mathrm{FC}]_0 + \mathrm{IC}_{50}) - (([\mathrm{SM}]_0 + [\mathrm{FC}]_0 + \mathrm{IC}_{50})^2 - 4[\mathrm{SM}]_0[\mathrm{FC}]_0)^{0.5}\} \qquad \textbf{(Eq.3)}$$

where $\mathrm{I}$ and $\mathrm{I}_0$ are the fold change of fluorescence intensity with or without compound, respectively, b is the maximal possible fold change of fluorescence intensity with the infinite time, $[SM]_0$ and $[FC]_0$ are the concentrations of the compound and the fold change of fluorescence intensity, respectively, and $IC_{50}$ is to be fitted by the equation.

**Optical Melting Experiments.** The thermal stabilities of WT, DDPAC, and DDPAC+I17T model RNAs were analyzed by optical melting in the presence or absence of compound **1** or **2** as previously described.[18] The RNAs (2 µM) were heated to 95 °C in 1× PBS buffer and slowly cooled to room temperature. Compound **1** (10 µM; 0.1% (v/v) DMSO), **2** (1.5 µM; 0.1% (v/v) DMSO) or vehicle (0.1% (v/v) DMSO) were then added to the RNA solution to generate samples. The samples were then further cooled to 15 °C and heated to 85 °C at a rate of 1 °C per min. Absorbance as a function of temperature was measured at 260 nm using a Beckman Coulter DU800 UV/vis spectrophotometer. Melting curves were fit to two-state model with MeltWin 3.5.[31]

**In Vitro Chemical Cross-linking and Isolation by Pull Down (Chem-CLIP) and Competitive Chem-CLIP (C-Chem-CLIP).** Growth medium [Dulbecco's modified eagle medium (DMEM; Corning, catalog # 10-013-CM) with 1× Glutagro (Corning)] and 10% (v/v) fetal bovine serum (FBS; Sigma, catalog # 12003C)] was inactivated by heating at 95 °C for 15 min and slowly cooling to room temperature. Approximately 10,000 counts per min (cpm) of $^{32}$P 5′-end labeled tau WT, DDPAC, WT+I17T, or DDPAC+I17T RNA were folded in 10 µL growth medium by heating at 65 °C for 5 min and slowly cooling to room temperature. Chem-CLIP probe **3** (final concentrations of 0.1 – 50 µM) or negative control probe **4** (final concentrations of 0.1 – 50 µM) was then added into the folded RNA in a total volume of 20 µL, and the mixtures were incubated at 37 °C for 1 h. For C-Chem-CLIP, dilutions of competing compound **2** (final concentrations of 0.1 – 50 µM) were incubated with RNA for 30 min before adding **3** (final concentrations of 1.5 µM). The samples were then cross-linked by UV irradiation with 365 nm light in a UV Stratalinker 2400 (Stratagene) for 10 min (lids of the microcentrifuge tube open).

The Chem-CLIP probe **3** or control probe **4** was then clicked to disulfide biotin azide (0.2 µL of 10 mM solution prepared in DMSO solution per sample; Click Chemistry Tools, catalog # 1168-10). After addition of 20 µL of 25 mM HEPES, pH 7, a mixture of 30 µL (v/v/v = 1:1:1) of sodium ascorbate (250 mM in $H_2O$), $CuSO_4$ (10 mM in $H_2O$) and THPTA (50 mM in $H_2O$) was added, and the samples were incubated for 2 h at 37 °C. Then, Dynabeads MyOne Streptavidin C1 Beads (20 µL slurry per sample;, Invitrogen) were added to the mixtures for pull-down of the cross-linked RNA. (The slurry was used directly from the manufacturer; that is the beads were not washed prior to use.) The samples were shaken for 20 min at room temperature and then washed three times with 1× PBST (phosphate buffered saline containing 0.1% (v/v) Tween-20). After the incubation period and for each wash, the beads were collected by placing reaction tubes on a magnetic separation rack, and the buffer was collected and pooled. The radioactive signal from bound RNAs (on the beads) and unbound RNAs (in the wash buffer) was measured by using a Beckman Coulter LS6500 Liquid Scintillation Counter as $R_{bound}$ and $R_{unbound}$. Percent capture was calculated as the percentage of radiolabeled RNA captured by beads calculated as $R_{bound}/(R_{bound}+R_{unbound})$.

**Cell Culture.** All cells were grown at 37 °C with 5% $CO_2$. HeLa cells were grown in growth medium [Dulbecco's modified eagle medium (DMEM; Corning, catalog # 10-013-CM) with 1× Glutagro (Corning, catalog # 25-015-CI)] and 10% (v/v) fetal bovine serum (FBS; Sigma, catalog # 12003C) supplemented with 1× Antibiotic/Antimycotic solution (Gibco, catalog # 15240062). HeLa cells were discarded after 30 passages. LAN5 neuroblastoma cells were grown in RPMI 1640 medium (Corning, catalog # 10-041-CV) supplemented with 1× penicillin/streptomycin solution (Corning, catalog # 30-002-CI) and 20% (v/v) FBS. LAN5 cells were discarded after 20 passages. The protocol for culturing primary neurons from htau mice is described in **htau Mice Primary Neuron Experiments**.

**Cell Viability.** HeLa cells were grown to ~60% confluency in 96-well plates (Corning; cat. # 3598). The growth medium was then replaced with freshly prepared 100 µL medium containing DMSO [0.1% (v/v)] as vehicle, compound **2** (final concentration of DMSO is 0.1% (v/v)), tau ASO (0.5 µM) and scrambled ASO (0.5 µM) for 48 h at 37 °C. The medium was then removed, and the cells were washed with 1× DPBS. After removing the DPBS, the cells were incubated in 100 µL of a mixture containing 10 µL WST-1 (Roche) reagent and 90 µL of growth medium for 40 min. The absorbance of the solution was measured at 450 and 690 nm with a SpectraMax M5 plate reader (Molecular Devices). $A_{690}$ (background) was subtracted from $A_{450}$ to obtain $A_{450}$' for analysis. $A_{450}$' of the cells treated with compounds or ASO were normalized to vehicle-treated cells to determine the effect of the compound on cell viability.

**Luciferase Reporter Assay to Study Exon 10 Alternative Splicing.** The exon 10 mini-genes, both WT and DDPAC and mutant thereof, fused to firefly luciferase, have been previously described.[18] HeLa cells were seeded in 60 mm diameter dishes and incubated for 12-16 h until they reached ~70% confluency. They were then transfected with WT, WT+I17T, DDPAC, or DDPAC+I17T luciferase reporter minigenes using JetPrime (Polyplus Transfection) per the manufacturer's recommended protocol and incubated at 37 °C for 4 h. [The transfection cocktail was added directly to the wells containing growth medium.] After transfection, the cells were collected by trypsinization and re-seeded in 96-well plates at $0.2 \times 10^5$ cells per well. After allowing the cells to adhere to the plate for 4 h at 37 °C, they were treated with compound at the indicated concentration for 48 h. In brief, the transfection cocktail was removed from the cells and replaced with compound-containing growth medium prepared as follows. The compound of interest (stocks prepared in DMSO) was diluted in growth medium at the appropriate concentration (final DMSO concentration at 0.1% (v/v)) was then added (10 µL per well).

After a 48-h treatment period, the cells were washed with 1× DPBS. To each well was added 100 µL of a mixture containing 10 µL WST-1 reagent and 90 µL of growth medium. After incubation at 37 °C for 40 min, absorbance was measured at 450 nm and 690 nm. $A_{690}$ (background) was subtracted from $A_{450}$ to obtain $A_{450}$' for analysis. After removal of WST-1/DMEM from wells, the cells were washed again with 1× DPBS and treated with 100 µL of One-Glo EX reagent (Promega, E8110) per well. The plate was incubated at room temperature for 20 min and the luminescence was measured with a Biomek FLx800 plate reader (1000 ms integration time). The resulting luminescence was normalized to $A_{450}$'. Finally, all viability-normalized luminescence values were normalized to vehicle, set to 1.

**RNA Isolation and Real-Time Quantitative PCR (RT-qPCR).** HeLa cells were transfected as described in the **Luciferase Reporter Assay to Study Exon 10 Alternative Splicing**. After transfection, the cells were collected by trypsinization and re-seeded in 24-well plates ($1 \times 10^5$

cells per well) to afford ~50% confluency. After allowing the HeLa cells to adhere to the plate for 4 h at 37 °C, the transfection cocktail was removed and replaced with growth medium containing the compound of interest or ASO (500 µL per well) for 48 h.

LAN5 cells were seeded in 12-well plates ($0.3\times10^6$ cells per well). After allowing the LAN5 cells to adhere to the plate for 12 h at 37 °C (~60% confluent), they were treated with vehicle, compound, or ASO, and incubated for 48 h. (Compound stocks were prepared in DMSO, and ASOs were prepared in water. Both modalities were diluted in growth medium to the appropriate concentration and added to the cells (1000 µL per well).)

Total RNA was extracted from both cell types and analyzed via RT-qPCR to determine levels of 3R and 4R tau mRNA. In brief, the cells were washed with 1× DPBS and then lysed in the well using 300 µL of Lysis Buffer provided in a Quick-RNA MiniPrep Kit (Zymo Research). Total RNA was extracted using the same kit per the manufacturer's protocol, including the on-column DNase I digestion. The concentration of total RNA was measured by NanoDrop 2000 (Thermo Fisher Scientific). Only samples with $OD_{260}/OD_{280}$ >1.8 were carried forward for analysis.

Reverse transcription (RT) was carried out on 200 ng of total RNA using a qScript cDNA Synthesis Kit (QuantaBio) in a total volume of 10 µL according to the manufacturer's protocol. An Applied Biosystems QS5 384-well PCR system was used to complete qPCR reactions with Power SYBR Green Master Mix (Life Technologies), 600 nM each primer (Table S3), 20 ng of cDNA (1 µL of the RT reaction), in a total volume of 33 µL. Technical triplicates (10 µL each) were aliquoted into 384-well qPCR plates (Applied Biosystems, catalog # 4483285). Subsequent qPCR analysis was performed using an Applied Biosystems QS5 384-well PCR system (software v.1.3.0). The 4R/3R ratio was determined using $\Delta\Delta C_t$ method using Equation 4:

$$\text{4R/3R ratio} = 2^{-(C_t\,4R-\,C_t\,3R)} \qquad \textbf{(Eq. 4)}$$

where $C_t\ 4R$ or $C_t\ 3R$ is the $C_t$ values for the 4R or 3R *MAPT* mRNA in total RNA from cells.

**Measuring Cellular Occupancy by Chem-CLIP and C-Chem-CLIP.** LAN5 cells were grown to ~80% confluency in 100 mm diameter dishes. The cells were then treated with 1.5 µM of **3** or 1.5 µM of **4** for 12 h. For C-Chem-CLIP, dilutions of competing compounds **1** or **2** were pre-incubated with cells for 12 h prior to adding **3**. After addition of **3**, the cells were incubated for an additional 12 h. After the treatment period, whether for Chem-CLIP or C-Chem-CLIP studies, the cells were washed with 10 mL of 1× DPBS. Ice-cold 1× DPBS (10 mL) was then added to the plate followed by irradiation with 365 nm light in a UV Stratalinker 2400 (Stratagene). Cells were scraped into the 1× DPBS, transferred to a 15 mL conical vial, and collected by centrifugation. Total RNA was extracted using TRIzol LS Reagent (500 µL; Invitrogen) per the manufacturer's protocol.

Approximately 20 µg of total RNA was then subjected to the click reaction with biotin azide and pull-down with magnetic streptavidin beads as described in **In Vitro Chemical Cross-linking and Isolation by Pull Down (Chem-CLIP) and Competitive Chem-CLIP (C-Chem-CLIP)**. After the washing steps, the cross-linked RNA captured by the beads was cleaved using a 1:1 mixture of TCEP (200 mM) and $K_2CO_3$ (600 mM) by incubating at 37 °C for 30 min with shaking. The reaction was quenched with 1 volume of Iodoacetamide (400 mM) by shaking at room temperature for 30 min. The isolated RNA was cleaned up by using RNA Clean XP beads (Beckman Coulter) according to manufacturer's protocol.

RT-qPCR was performed on RNA samples before and after pull-down as described in **Real-Time Quantitative PCR** to enable calculation of fold enrichment in the pulled down samples. Relative fold enrichment of tau pre-mRNA was measured using Equation 5:

$$\text{Relative Fold Enrichment} = 2^{-(\Delta C_t \text{ before pull-down} - \Delta C_t \text{ after pull-down})}$$

**(Eq. 5)**

where "$\Delta C_t$ before pull-down" is the difference between the $C_t$ values for the *MAPT* mRNA and 18S rRNA in total RNA from cells and "$\Delta C_t$ after pull-down" is the difference between the $C_t$ values for the *MAPT* pre-mRNA and 18S rRNA in RNA in the pulled down fractions.

**Chem-CLIP-Map.** Chem-CLIP-Map was performed as previously described.[32] Briefly, LAN5 cells were grown to ~80% confluency in 100 mm diameter dishes and treated with 1.5 μM of **3** for 12 h. Total RNA was extracted and pulled down as described in **Measuring Cellular Occupancy by Chem-CLIP and C-Chem-CLIP**. RT was carried out using 1 μg of RNA pulled down by **3**, 2 pmol of a *MAPT* gene-specific reverse primer (5'-CAGACGTGTGCTCTTCCGATCTGACACTCCAGTCCCACAGT-3'), and 1 unit of SuperScript III (Invitrogen, 18080400) in a total volume of 10 μL per the manufacturer's protocol. Please see Fig. S15A for a schematic of the gene-specific primer binding site. The blue sequence binds nt 1790-1809 in *MAPT* mRNA, ~1437 nt downstream of the cross-linked site.

After digestion with RNase A and RNase H (part of the Superscript III protocol), the cDNA from the RT reaction was cleaned up using RNA Clean XP beads (Beckman Coulter) according to the manufacturer's protocol. The purified cDNA was ligated to a 3' ssDNA adaptor (5'-Phosphate-NNNAGATCGGAAGAGCGTCGTGTAG-3C spacer) by incubating 2 μL 10× T4 RNA Ligase Buffer, 1 μL of 1 mM ATP, 10 μL 50% (w/v) PEG 8000, 5 μL cDNA, 1 μL of 20 μM ssDNA adaptor, and 1 μL of T4 RNA Ligase (New England BioLabs, catalog # M0204S). The ssDNA adaptor is the red sequence of the ligated cDNA in Fig.S15A. The ligated cDNA was purified again with RNAClean XP beads as described above.

PCR amplification was performed with the purified cDNA ligated with a 3' adaptor by using Phusion polymerase (NEB) with 25 cycles of 98 °C for 30 s, 65 °C for 20 s, and 72 °C for 60 s. The forward and reverse primers were 5'–CAGACGTGTGCTCTTCCGATC-3' and 5'–CTACACGACGCTCTTCCGATCT-3', respectively. The forward primer is nt 1-21 of the *MAPT* gene-specific reverse primer shown above (green sequence). The reverse primer binds to the ssDNA adaptor (red sequence) corresponding to nt 1461-1482 of the ligated cDNA. The PCR product (~1.5k bp) was excised from the gel and ethanol precipitated with 20 μg of glycogen (Thermo Scientific, catalog # R0561) per sample. A second round PCR was performed to amplify the purified DNA. The DNA from second round PCR was used without further purification. Approximately 100 ng of this DNA was ligated directly into a vector provided with NEB's PCR Cloning Kit (E1202S) per the manufacturer's protocol. Ampicillin-resistant colonies were selected and subjected to Sanger sequencing, completed by Eton Biosciences.

**Western Blotting.** LAN5 cells were seeded in 6-well plates at $0.7 \times 10^6$ cells per well. After allowing the LAN5 cells to adhere to the plate for 12 h at 37 °C (reaching ~60% confluency), they were treated with compound of interest and incubated for 48 h. After removing the growth medium, the cells were washed with 1× DPBS and harvested by trypsinization. Total protein was extracted with Mammalian Protein Extraction Reagent (M-PER, Thermo Scientific) per the manufacturer's protocol. Protein concentrations were determined using a Pierce Micro BCA Protein Assay Kit per the manufacturer's recommended protocol. Approximately 20 μg of total protein from each biological sample was separated on a sodium dodecyl sulfate (SDS)-

polyacrylamide gel (5% (w/v) polyacrylamide stacking layer; 12% (w/v) polyacrylamide separating layer), followed by transfer to a polyvinylidene fluoride (PVDF; 0.45 μm) membrane. After blocking the membrane in 1× TBST [Tris-Buffered Saline with 0.1% (v/v) Tween-20] with 5% (w/v) nonfat milk for 1 h, the membrane was incubated with 1× TBST containing 5% (w/v) milk and primary antibodies for 3R and 4R tau simultaneously as follows: 4R tau antibody (MilliporeSigma, catalog # 05-804; 1:2000 dilution) or 3R tau antibody (MilliporeSigma, catalog # 05-803; 1:5000 dilution) and incubated at 4 °C overnight. The membrane was washed three times with 1× TBST for 10 min each and then incubated with a horseradish peroxidase (HRP)-conjugated secondary antibody (Cell Signaling Technology, catalog # 7076; 1:5000 dilution) in 1× TBST containing 5% (w/v) milk for about 1 h at room temperature. After three more washes with 1× TBST for 15 min each, 4R and 3R tau protein were detected using SuperSignal West Pico Chemiluminescent Substrate (Pierce Biotechnology).

After imaging, the membrane was stripped with 1× Stripping Buffer (200 mM glycine with 0.1% SDS, pH 2.2) for 0.5 h at room temperature, and β-actin levels were detected as described above, except using a β-actin primary antibody (Cell Signaling Technology, 4970S; 1:5000 dilution). ImageJ software was used to quantify the protein bands.[33]

**Isolation and Treatment of Primary Neurons from Humanized tau (hTau) Transgenic Mice.** All animal studies were completed as approved by the Scripps Florida Institutional Animal Care and Use Committee. All the required Dissection Medium, Plating Medium, Feeding Medium, and miscellaneous reagents were prepared as previously described.[18] The Dissection Medium consisted: 430 mL of culture-grade water (Fisher Scientific, SH3052902), 50 mL of 10× HBSS without $Ca^{2+}$ and $Mg^{2+}$ (Invitrogen, 14185052), 10 mL of HEPES (Invitrogen, 15630080), 5 mL of pyruvate (Invitrogen, 11360070), 5 mL of glucose solution (Thermo, A2494001), and 100 μL of gentamicin (Invitrogen, 15710064). Plating Medium is composed of 465 mL of Neurobasal (Invitrogen, 21103049), 25 mL of FBS, heat-inactivated (Invitrogen, 10082139), 10 mL of Glutamax-I (Invitrogen, 35050061), and 100 μL of gentamicin (Invitrogen, 15710064). Feeding Medium contains the following: 490 mL of Neurobasal-A (Invitrogen, 10888022), 10 mL of Glutamax-I (Invitrogen, 35050061), and 100 μL of gentamicin (Invitrogen, 15710064).

Culture plates (Corning CellBIND 24-well Surface Microplates, catalog # 3337) were pre-coated with 0.1 mg/mL Poly-D-Lysine (PDL), which was dissolved in 10 mM Tris, pH = 7.4, for 2 h at 37 °C. The amounts of media required for isolation and culture of primary neurons are as follows: 1 mL per pup of Dissection Medium, placed on ice; another 1 mL per pup of Dissection Medium supplemented with 1% (v/v) Papain pre-warmed at 37 °C; 5 mL per pup of Plating Medium for the dissociation step, and 25 mL of Feeding Medium per plate pre-warmed at 37 °C.

P0 htau mouse pups were obtained from breeding of [B6.Cg-Mapttm1(EGFP)Klt Tg(MAPT)8cPdav/J] mice, which acquired from Jackson Labs. Genotype was confirmed by PCR amplification as previously described.[18] For one 24-well plate, the cortex of three P0 htau mouse pups were removed and each placed in the 15 mL tubes containing 1 mL of ice-cold Dissection Medium for dissection and removal of meninges. After removal of the meninges as previously described,[34] the cold Dissection Medium was aspirated, and 1 mL of Dissection Medium containing Papain solution pre-warmed to 37 °C was added into each tube. The samples were incubated at 37 °C for 20 min, and then 1 mL of Plating Medium was added to each tube to quench Papain activity. During the Papain quenching, a PDL pre-coated 24-well culture plate described above was washed with 1× DPBS, 12 mL of Feeding Medium was added (0.5 mL per well), and the plate was incubated at 37 °C.

The Dissection and Plating Media in each 15 mL tube was aspirated, and the remaining tissue was washed three times with 1 mL of pre-warmed Plating Medium. After the washing steps, 2 mL of freshly prepared Plating Medium was added, and the tissue was dissociated by gently triturating through a 5 mL serological pipette three times and then a P-1000 pipet three times. Next, 1 mL of the supernatant was transferred to a new 15 mL tube, to which was added 1 mL of freshly prepared Plating Medium. Trituration was repeated three times using 1 mL of Plating Medium and a P-1000 pipet until the tissue was homogenized and all chunks of tissue were dissociated; 1 mL of supernatant was transferred each time from each 15 mL tube and collected in a new 15 mL conical vial separately.

The cell mixtures were diluted to 10 mL with Plating Medium and strained through a 40 µm cell strainer into a 50 mL tube. The cells were centrifuged at 1900 rpm for 4 min, and the three pellets were combined and resuspended in 1 mL freshly prepared Plating Medium for cell counting. After cell counting, the cells were diluted with 12 mL of Feeding Medium supplemented with 2% (v/v) B-27 to afford to $0.6\times10^{6}$ /mL. The cells were then aliquoted (0.5 mL per well) into pre-warmed PDL pre-coated 24-well culture plate. Half of the Feeding Medium (0.25 mL) supplemented with 2% (v/v) B-27 was replaced every 3-4 days.

After 15 days, the time period required for detectable levels of 3R and 4R tau,[18] neurons were treated with **1**, **2**, tau ASO, or scrambled ASO for 48 h at 37°C. In brief, compound or ASO was prepared as 2× final concentration in Feeding Medium. In the 24-well culture plates, half of the Feeding Medium was removed and then replaced with the 2× compound or ASO solution. After 48 h, Total RNA was extracted, and RT-qPCR was performed as described in **RNA Isolation and Real-Time Quantitative PCR (RT-qPCR)**.

**Plasma and brain protein binding.** Plasma protein binding and non-specific brain binding were determined using equilibrium dialysis. All samples were tested in triplicate using the RED Rapid Equilibrium Dialysis Device (Thermo Scientific). The initial drug concentration in the plasma or brain homogenate (25% brain) chamber was 2 µM, and phosphate buffered saline was added to the receiver chamber. The plate was covered and allowed to shake in a 37 ºC incubator for 6 h. A 25 µL aliquot was sampled from the plasma/brain and PBS chambers, diluting with either blank PBS or plasma/brain to achieve a 1:1 ratio for all samples. The concentration of **2** was determined by LC-MS/MS. The plasma fraction bound was calculated as ([plasma] – [PBS]) / [plasma]. For non-specific brain binding, the directly measured value is for a 25% brain homogenate, and this is converted to 100% tissue using the following equation: Free Fraction as % = 100*(1/dilution factor)/((1/Observed Free-1)+1/dilution factor).

**Quantification of 2 in Mouse Plasma and Brain**. Male C57BL/6J mice (n = 3 per time point; 5-7 weeks) were orally administered **2** (100 mg/kg) in a formulation of DMSO/Tween-80/$H_2O$ (5/5/90). After 2, 12, 24, and 48 h, mice were euthanized, and blood and brain were collected. Brains were immediately frozen. Blood was centrifuged to generate plasma and immediately frozen. On the day of analysis, the plasma and brain homogenate were deprotonated by adding acetonitrile carbamazepine as an internal standard at a ratio of 1 part plasma or brain homogenate to 5 parts acetonitrile in a 96-well Multiscreen Solvinter 0.45 µm low binding PTFE hydrophilic filter plate (Millipore, catalog # MSRLN0450) and let sit on ice for 15 minutes with occasional shaking to precipitate proteins. The filtrate was collected into a capture 96-well plate (Corning; cat. # 3598) upon centrifugation and directly analyzed using mass spectrometry. Parallel samples were processed using the same conditions to generate standard curves between 2 and 2000 ng/ml. The analyte was spiked into blank plasma or blank brain homogenate and processed alongside the samples. Drug levels were determined by mass spectrometry using an ABSciex 5500 mass spectrometer with multiple reaction monitoring. Compound **2** was detected using the mass transition m/z 435→236. The first mass corresponds to **2** in positive ion mode and the second mass is the fragment ion that was followed after fragmenting the compound in the collision cell.

**Pharmacokinetic profiling of 2 in plasma.** To understand steady state levels of **2** in plasma, reflective of the multiple dosing carried out in therapeutic efficacy studies with htau mice (below), three experiments were carried out: (i) steady state dosing was achieved by dosing male C57BL/6J mice *p.o.* with 100 mg/kg of **2** as follows: Day 1 at 7 am and 5 pm; Day 2 at 7 am; and Day 3 at 7 am, followed by collection of data at the indicated time points; (ii) upon a single dose delivered by *p.o* of 100 mg/kg of **2** on Day 3 at 7 am, followed by collection of data at the indicated time points; and (iii) upon a single dose delivered by *i.v.* of 5 mg/kg of **2** on Day 3 at 7 am, followed by collection of data at the indicated time points. The concentration of **2** in plasma was measured as described in "**Quantification of 2 in Mouse Plasma and Brain**".

**htau Mice Studies.** htau mice and WT mice (C57BL/6J) were purchased from Jackson Laboratories. htau mice were maintained in-house through breeding between htau+/- mice. Genotype was confirmed by PCR amplification as previously described.[18] Only homozygous mice were used in in vivo studies. Mice were randomly assigned to a treatment group, which were age- and gender-matched (n = 6). In these studies, mice were >9 months of age since cognitive and physiological impairments in htau mice is age-dependent and were not presented in young mice (<4 months) with early-stage tau pathology.[35] Mice were orally administered vehicle (5/5/90 DMSO/Tween-80/$H_2O$) or 100 mg/kg of **2** in the same formulation every other day (q.o.d.). During the treatment period, nesting activity was assessed, as describe in **Nesting Activity** below. After 20 days, the mice were euthanized (in accordance with guidelines provided by the American Veterinarian Medical Association), and the brain was harvested for analysis. One hemisphere of the brain was frozen at -80 °C for RT-qPCR analyses while the other was used for histological studies.

**Nesting Activity.** From Day 1 to Day 20, mice were individually housed, and each day the old nesting material was collected and replaced with a new nestlet (3.0 g). Around 15 h later, the resulting nest was photographed for qualitative scoring, and the untorn nestlet was weighed if present. All nest images and weights were analyzed by a blinded individual and based on published criteria.[36] In brief, nesting is scored on a rating scale of 1–5 where if the nestlet is over 90% intact, it is given a nesting score is 1 (Poor; Figure 4C). When the nestlet is partially torn but >50% remains, the nesting score is 2; a score of 3 is assigned when 50-90% of the nestlet is torn but spread around the cage and no identifiable nest site is found (Medium; Figure 4C). When more than 90% of the nestlet is torn and the nest is identifiable but flat, the nesting score is 4. When a near perfect nest is built and the wall is higher than the mouse's body, it is assigned a nesting score of 5 (Good; Figure 4C).

**Brain Tissue Histology.** Left brain hemispheres of htau mice were harvested for total RNA. Right brain hemispheres (unfrozen) of htau mice were stored in 10% neutral buffered formalin (VWR) for 48 h. Tissue processing, embedding, and sectioning were performed and generated by the Scripps Florida Histology Core. Briefly, tissue was embedded in paraffin using a Sakura Tissue-Tek VIP5 paraffin processor, sectioned at 4 μm, and then mounted on positively charged slides for further immunostaining. The slides were stained on a Leica BondMax Immunostaining platform with the following primary antibodies: AT8 (MN1020, Invitrogen; 1:100 dilution) and NeuN (MAB377B, Chemicon; 1:500 dilution). After the slides were washed three times with 1× DPBS, a DAB Substrate Kit (Vector Laboratories, Inc., cat #) was used to detect the primary antibodies per the manufacturer's protocol. After staining was complete, the slides were dehydrated, cover-slipped with Cytoseal 60 (Thermo Scientific), and imaged with a Leica DMI3000 B upright fluorescent microscope using 20× or 60× objective. Quantification was performed in a blinded fashion using ImageJ. Within a representative part of an image, AT8 positive area was calculated as area of the aggregates (purple staining of AT8) with diameter over 20 μm and staining signal over 185 divided by total area of the slide, and NeuN positive neurons were the aggregates (brown staining of NeuN) with diameter over 20 μm and staining signal over 200.

**Statistical Analysis.** All data are reported as the mean ± standard deviation (SD). Data were plotted and analyzed using commercially available software (Perseus, GraphPad Prism, and ImageJ). Statistical significance between experimental groups was analyzed either by two-tailed Student's t-test or one-way ANOVA followed by Bonferroni's multiple-comparison test. In all cases, p-values of less than 0.05 were considered statistically significant.

### Supplementary Figures and Tables

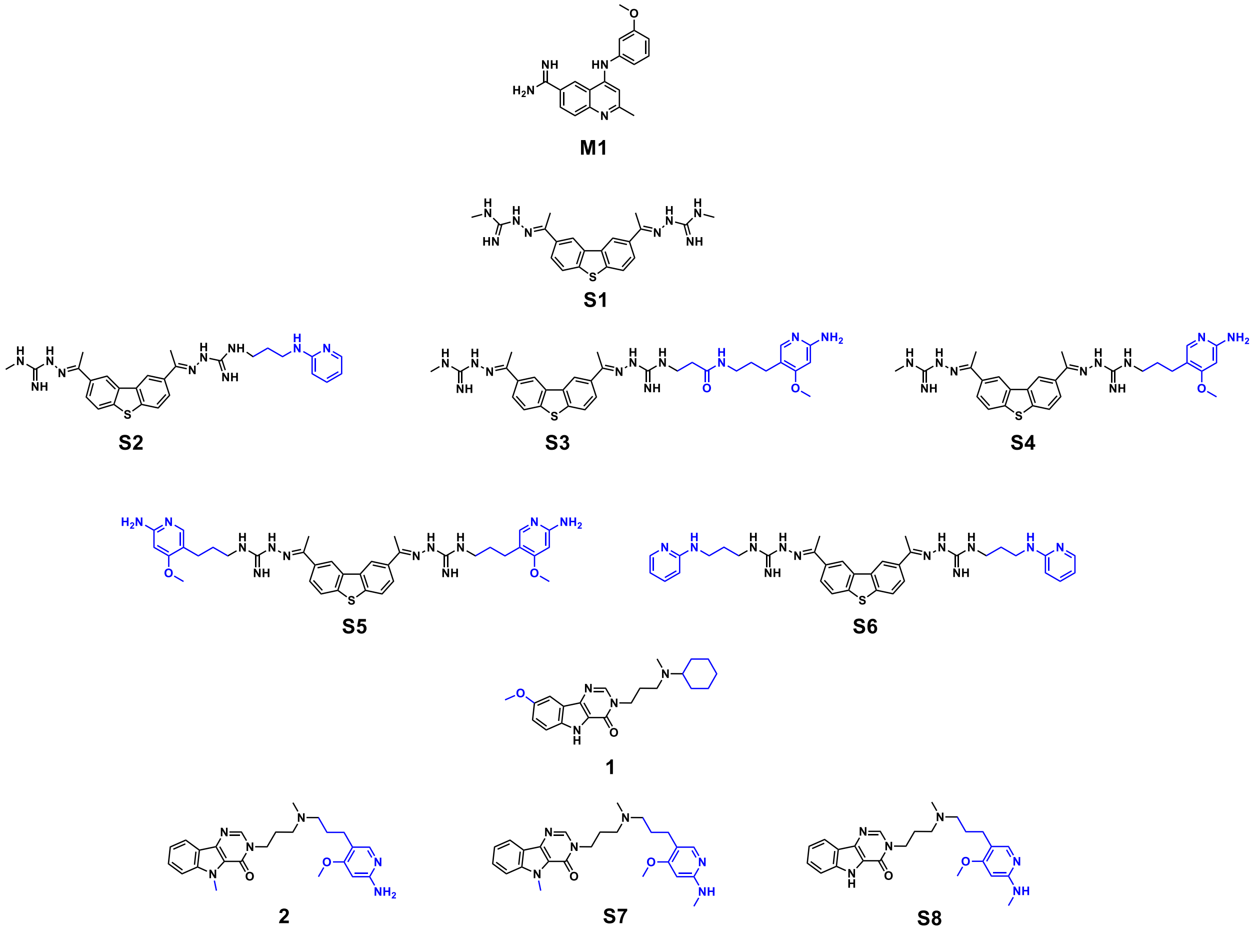

**Figure S1. Structures of S1, 1, and the eight 2-aminopyridine analogs that were screened in a cell-based luciferase assay.** **S1** and **1** were previously described and designed by an iterative process of file mining, in vitro and cellular assays, and structure-based design.[18] The structural differences of **S1**, **1** and their analogs are highlighted in blue. **M1** was not pursued for derivatization since **M1** has a low calculated binding energy (-6.39 kcal/mol, Table S1) due to its weak interactions (solely hydrogen bonds) with the A-bulge binding pocket.[18]

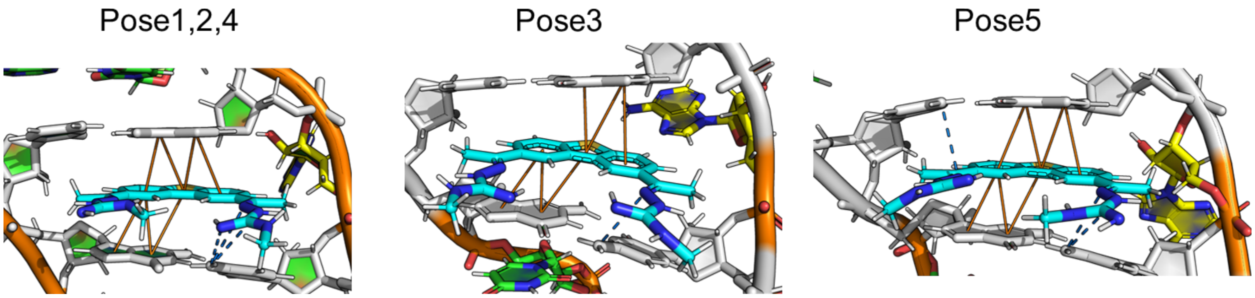

**Figure S2. Interaction network of the five structures with the lowest free energies extracted from the NMR ensemble of the RNA-S1 complex (PDB:6VA2[18]).** Stacking interactions with the neighboring GC base pair of the A-bulge (solid line) as well as hydrogen bonds (dashed lines) stabilize the bound state of **S1**. The RNA backbone is shown in cartoon and base pairs in stick representation. **S1** is shown in stick representation.

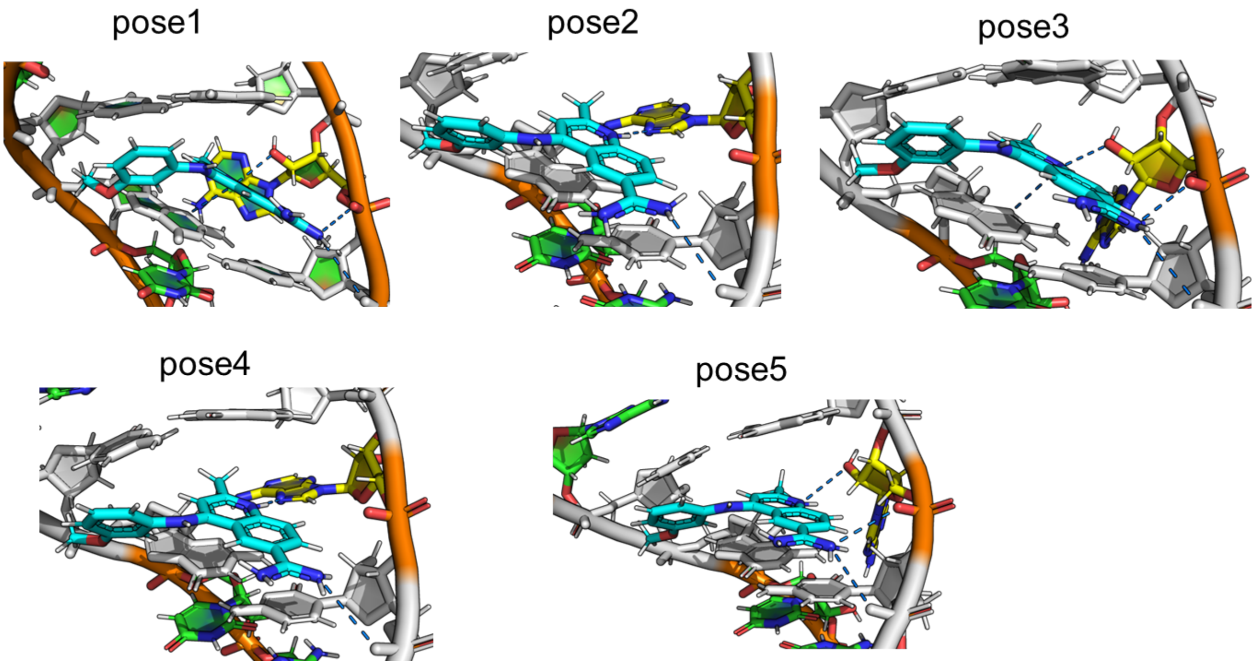

**Figure S3. Interaction network of the five structures with the lowest free energies extracted from the NMR ensemble of the RNA-M1 complex (PDB:6VA3[18]).** Hydrogen bonds with the neighboring GC base pair of the A-bulge (dashed lines) as well as with backbone stabilize the bound state of **M1**. The RNA backbone is shown in cartoon and base pairs in stick representation. **M1** is shown in stick representation.

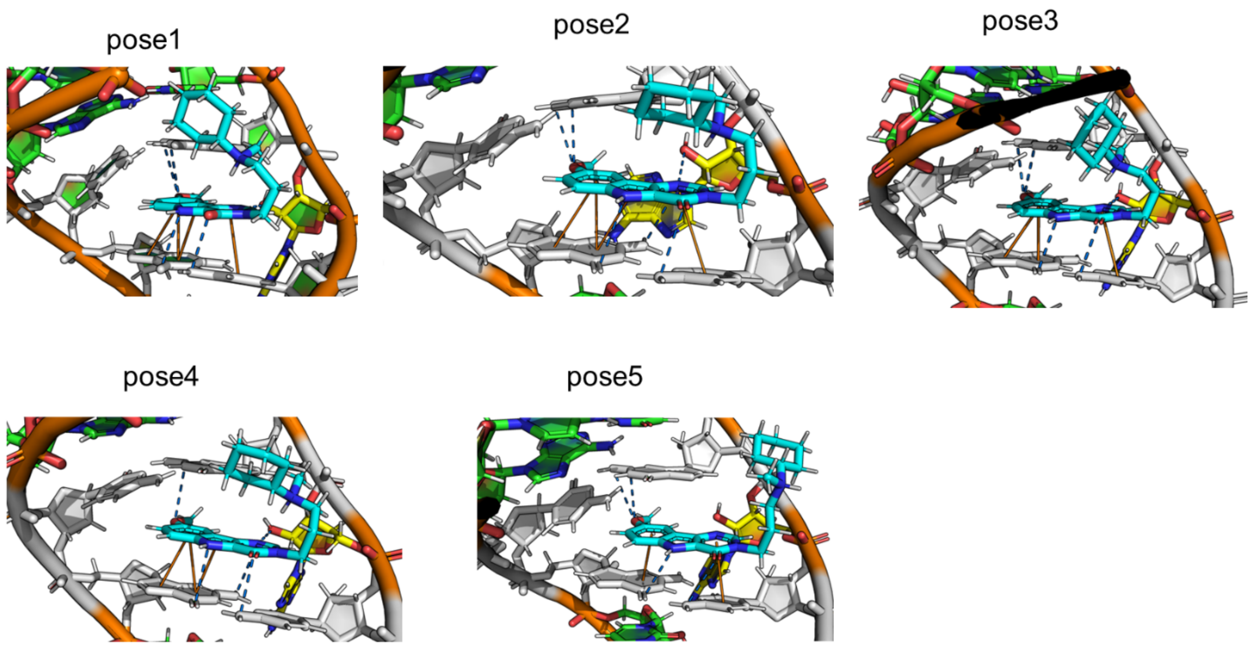

**Figure S4. Interaction network of the five structures with the lowest free energies extracted from the NMR ensemble of the RNA-1 complex (PDB:6VA4[18]).** Stacking interactions with the neighboring GC base pair of the A-bulge (solid lines) as well as hydrogen bonds (dashed lines) stabilize the bound state of **1**. The RNA backbone is shown in cartoon and base pairs in stick representation. **1** is shown in stick representation.

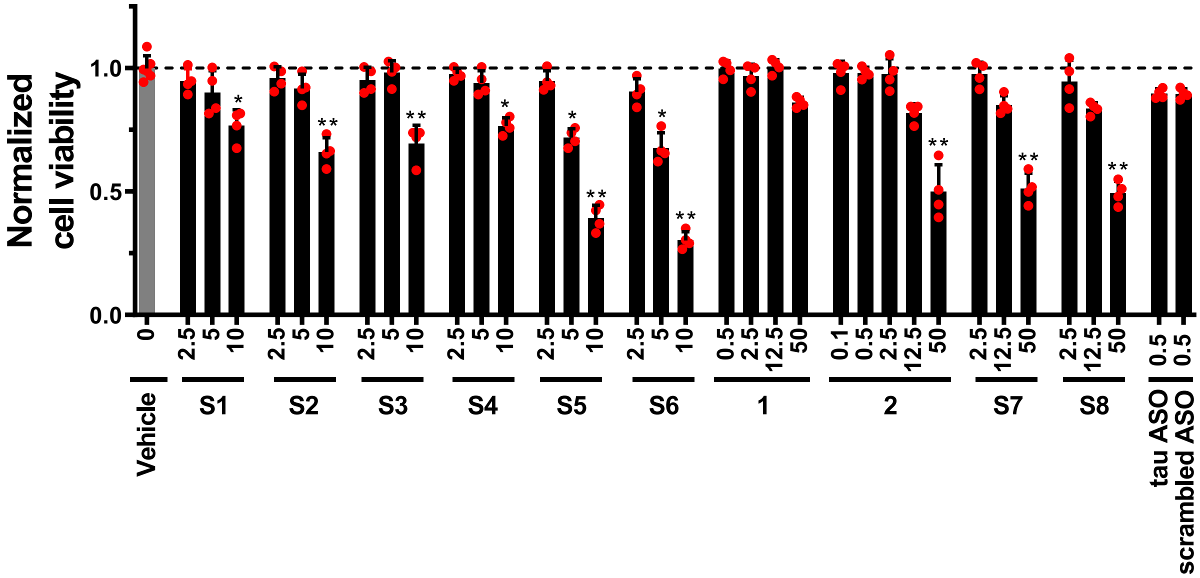

**Figure S5. Effect of 2 on the viability of wild type (untransfected) HeLa cells, assessed by WST-1 cell proliferation reagent (n = 6 biological replicates for DMSO vehicle; n = 4 biological replicates for compound or ASO).** *P < 0.05, **P < 0.01, as determined by one-way ANOVA.

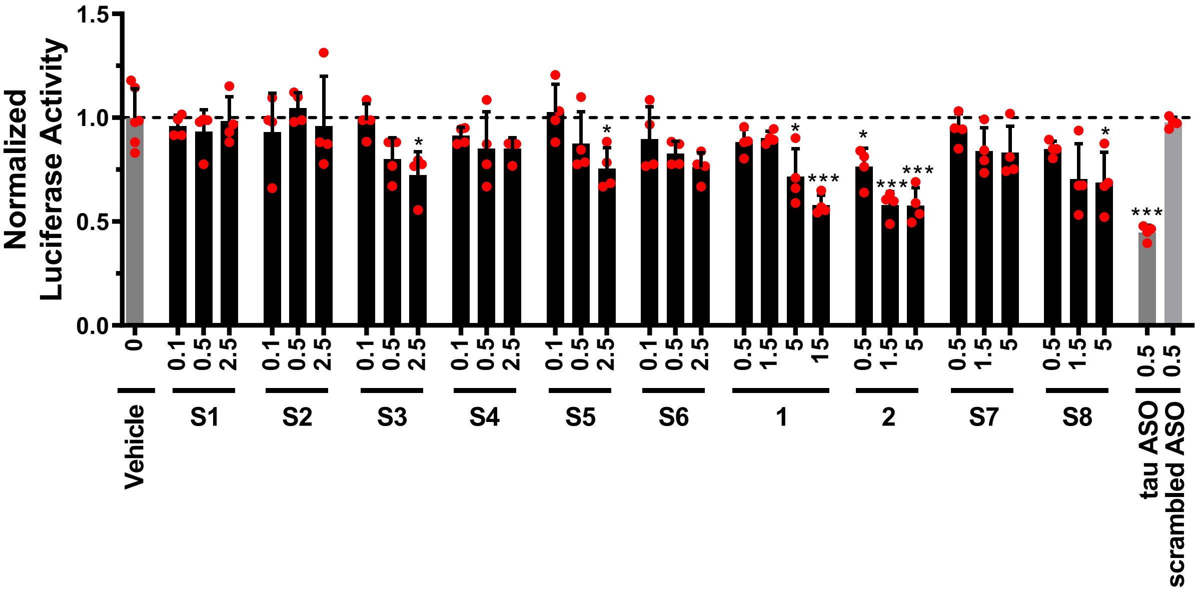

**Figure S6. Effect of parent molecules and base triple conjugates thereof on the alternative splicing of DDPAC *MAPT* exon 10 mini-gene.** The DDPAC mini-gene reporter was previously described,[37] where firefly luciferase is in-frame with exon 10. Thus, compounds that direct alternative splicing such that exon 10 is excluded reduce firefly luciferase activity. The mini-gene was transfected into HeLa cells, followed by treatment with vehicle or compound (n = 6 biological replicates for DMSO vehicle; n = 4 biological replicates for compound or ASO). Note that **1** was tested at concentrations of 0.5, 1.5, 5, and 15 μM while its 2-aminopyridine conjugates were evaluated at 0.5, 1.5, and 5 μM; **S1** and its 2-aminopyridine conjugates were treated at 0.1, 0.5, and 2.5 μM; Tau and scrambled ASO were tested at 0.5 μM; Top concentration of each compound was determined based on their effect on cell viability (Figure S5). "Normalized Luciferase Activity" indicates that the luciferase data were normalized to cell viability. $^{*}P < 0.05$, $^{**}P < 0.01$, $^{***}P < 0.001$, as determined by one-way ANOVA.

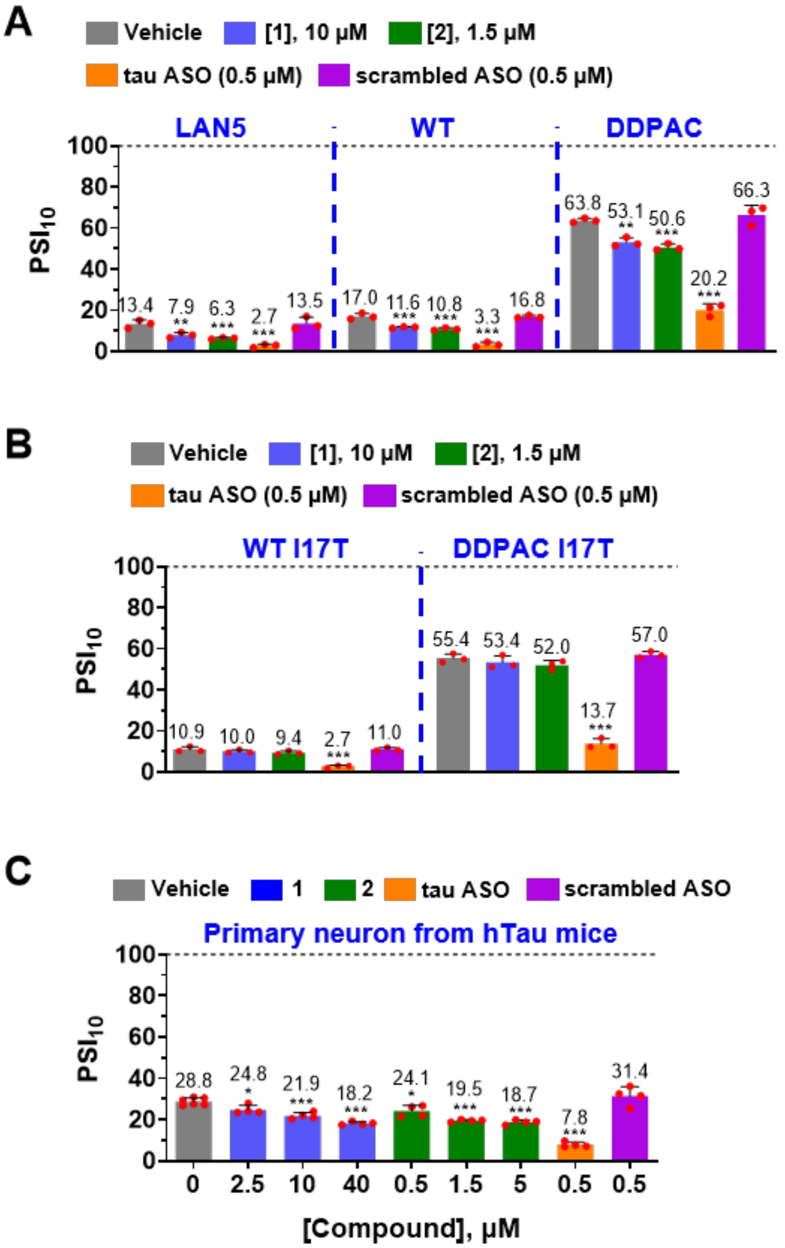

**Figure S7. Percent spliced in index (PSI) values of *MAPT* exon 10 are reduced by compound- or ASO-treatment.** Effect of **1**-, **2**-, or ASO treatment on $PSI_{10}$ (PSI of *MAPT* exon 10), as compared to vehicle in: (A) LAN5 cells and HeLa cells transfected with WT or DDPAC mini-genes (n = 3 biological replicates); (B) WT I17T or DDPAC I17T mutants (n = 3 biological replicates) , and (C) primary neurons harvested from htau mouse pups (n = 6 biological replicates for vehicle; n = 4 biological replicates for treated neurons). *, $p < 0.05$; **, $p < 0.01$; ***, $p < 0.001$, as determined by one-way ANOVA.

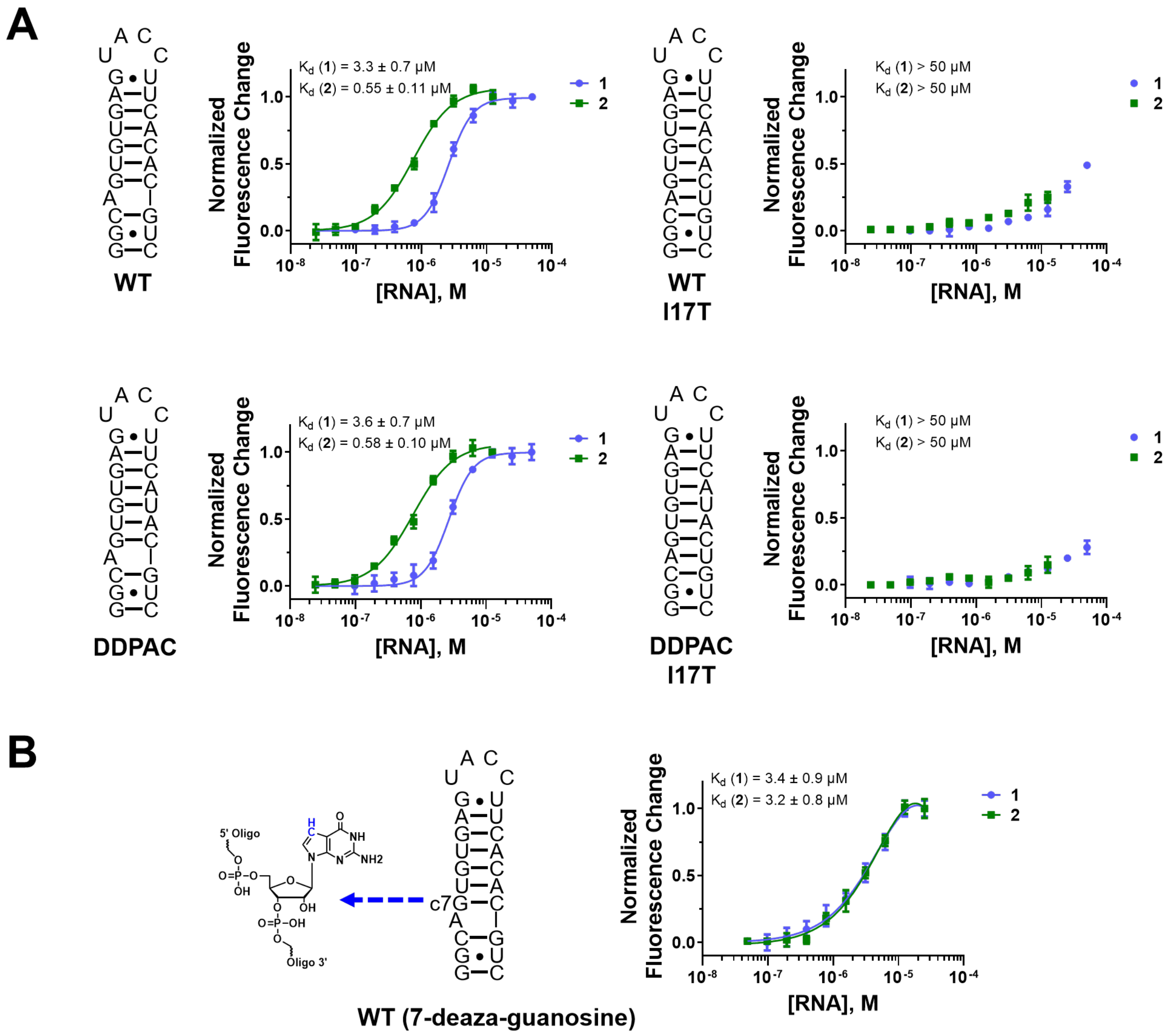

**Figure S8. In vitro binding measurements for 2 and models of various tau SRE RNAs.** (A) Representative binding curves for **1** or **2** and models of the WT, WT I17T, DDPAC, or DDPAC I17T *MAPT* SRE, where the intrinsic fluorescence of the small molecule was measured as a function of RNA concentration (n = 3 independent replicates). No saturable binding was observed with the addition of up to 50 µM of the I17T mutant RNAs. (B) Representative binding curves for **1** or **2** and a 7-deaza-guanosine-modified WT *MAPT* SRE RNA (n = 3 independent replicates).

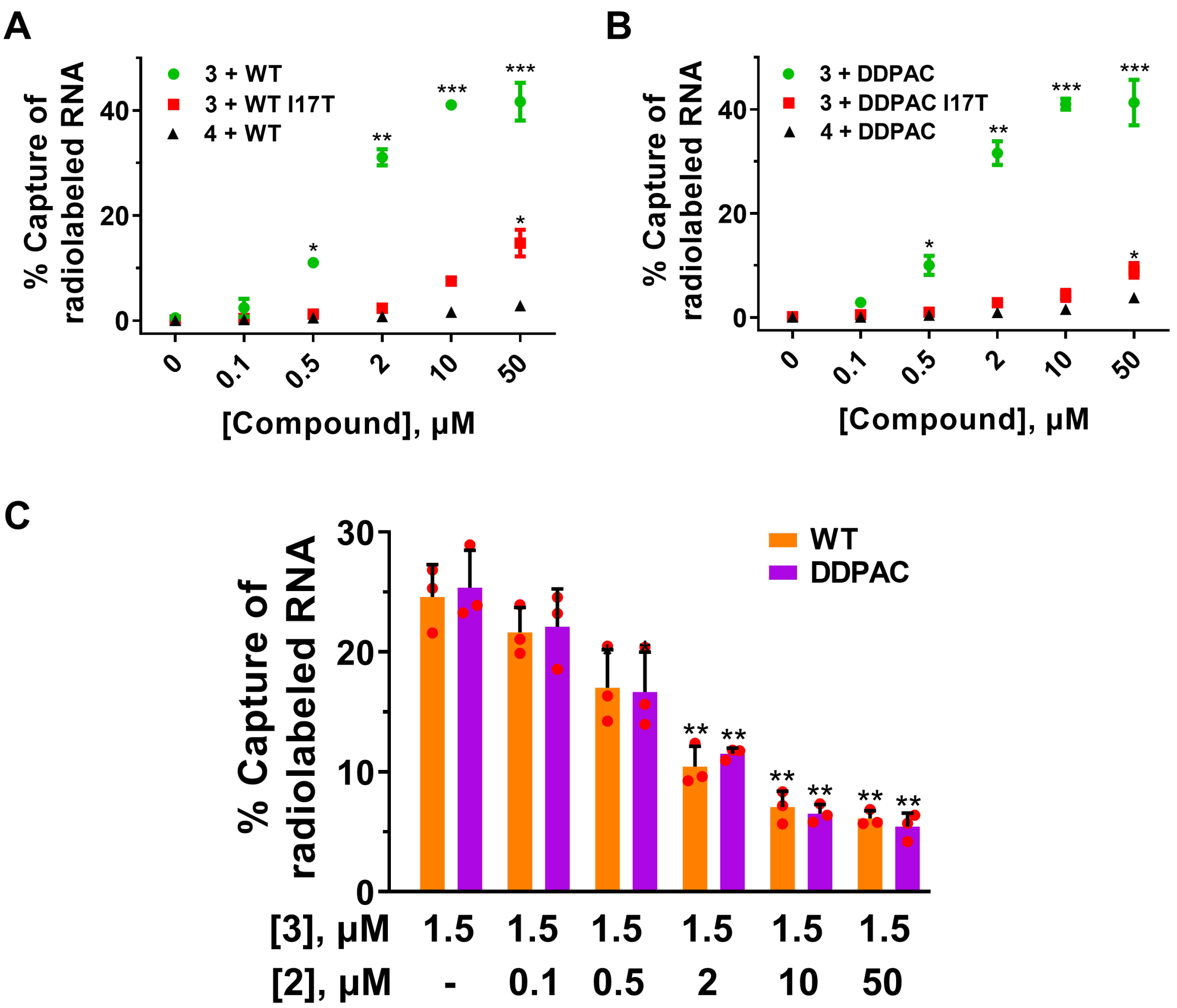

**Figure S9. In vitro Chem-CLIP experiments to study direct target engagement of tau SRE RNA by 2.** (A) Percentage of WT and WT+I17T mutant RNA pulled down in Chem-CLIP experiments as function of probe concentration, whether **3** and **4** (n = 2 independent replicates). (B) Percentage of DDPAC and DDPAC+I17T mutant RNA pulled down in Chem-CLIP experiments as function of probe concentration, whether **3** and **4** (n = 3 independent replicates). (C) Results of in vitro C-Chem-CLIP experiments with WT and DDPAC RNAs and **2** and **3** (n = 3 independent replicates). * $P < 0.05$, ** $P < 0.01$, and *** $P < 0.001$, as determined by one-way ANOVA. Data are reported as the mean ± SD, and in some cases the error bars are smaller than the data points.

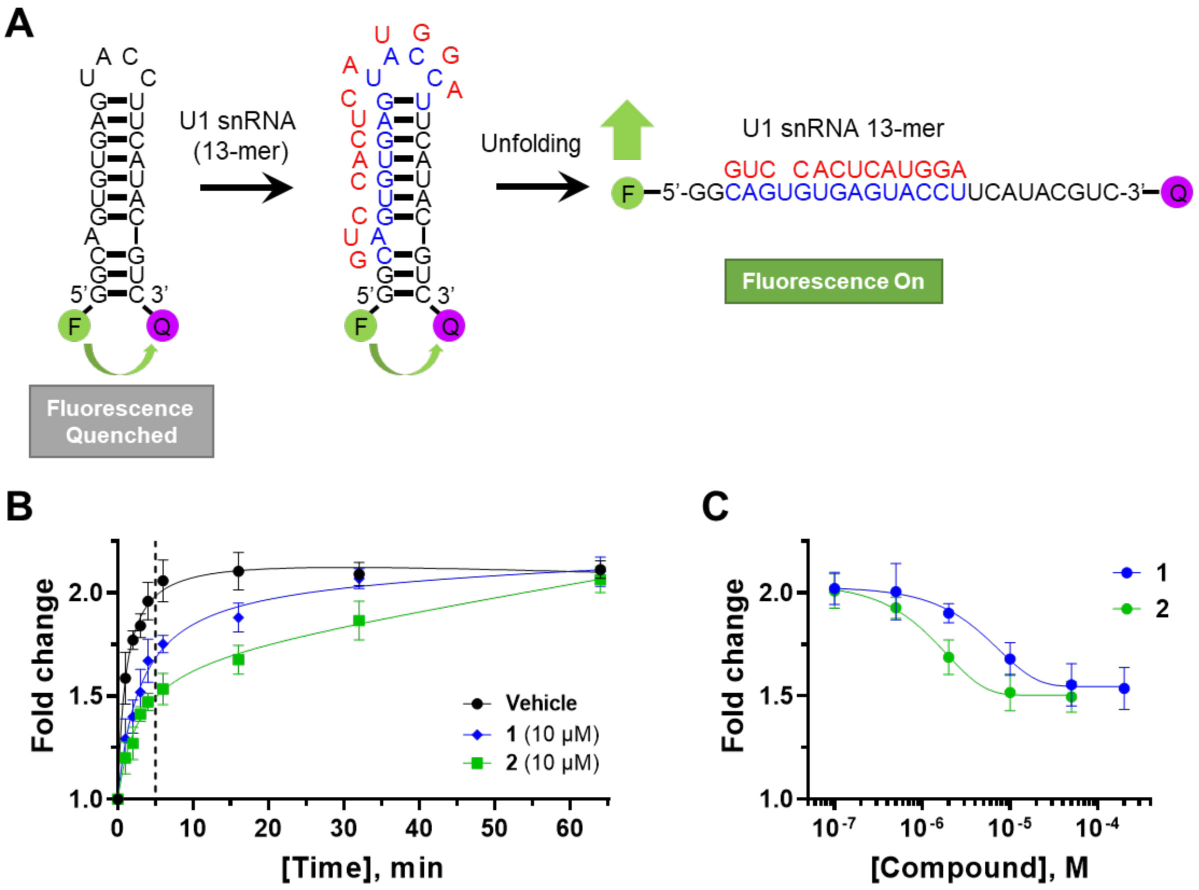

**Figure S10. Biochemical assay to assess the ability of 1 and 2 to prevent the binding of a model of U1 snRNA, which facilitates inclusion of exon 10.** (A) Schematic of FRET-based assay to assess inhibition of U1 snRNA binding of the tau SRE. (B) Effect of **1** and **2** (10 μM) on the ability of U1 snRNA to unfold a dually labeled tau RNA hairpin as a function of time (n = 4 independent replicates). (C) Fold-change in fluorescence intensity measured at t = 5 min (end-point measurement) when tau SRE was incubated with varying concentrations of **1** and **2** (n = 4 independent replicates). The highest concentration of compound **2** in this assay was 50 μM as it was not soluble at higher concentrations under assay conditions.

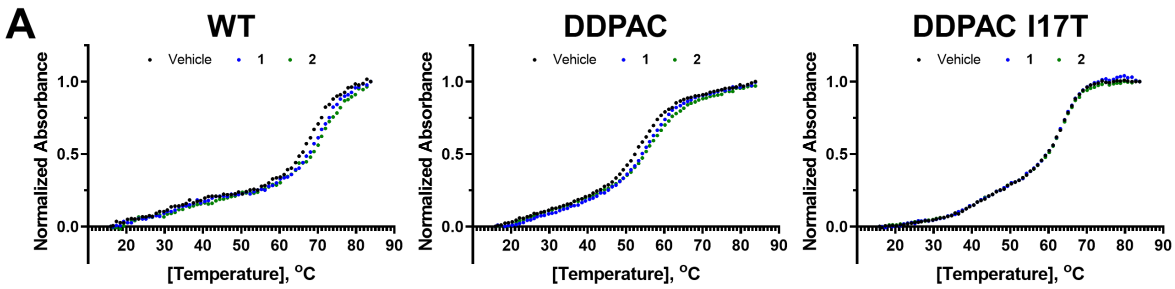

**B**

| | WT | | DDPAC | | DDPAC I17T | |
|---|---|---|---|---|---|---|
| | $T_m$/°C | ΔG/(kcal•mol$^{-1}$) | $T_m$/°C | ΔG/(kcal•mol$^{-1}$) | $T_m$/°C | ΔG/(kcal•mol$^{-1}$) |
| **Vehicle** | 68.5±0.2 | -6.5±0.3 | 53.4±0.2 | -2.7±0.1 | 64.3±0.2 | -4.0±0.1 |
| **1** | 69.1±0.1 | -7.1±0.3 | 54.6±0.1 | -3.8±0.2 | 64.6±0.2 | -4.1±0.1 |
| **2** | 69.3±0.1 | -7.6±0.1 | 54.7±0.1 | -4.3±0.2 | 64.3±0.2 | -3.9±0.1 |

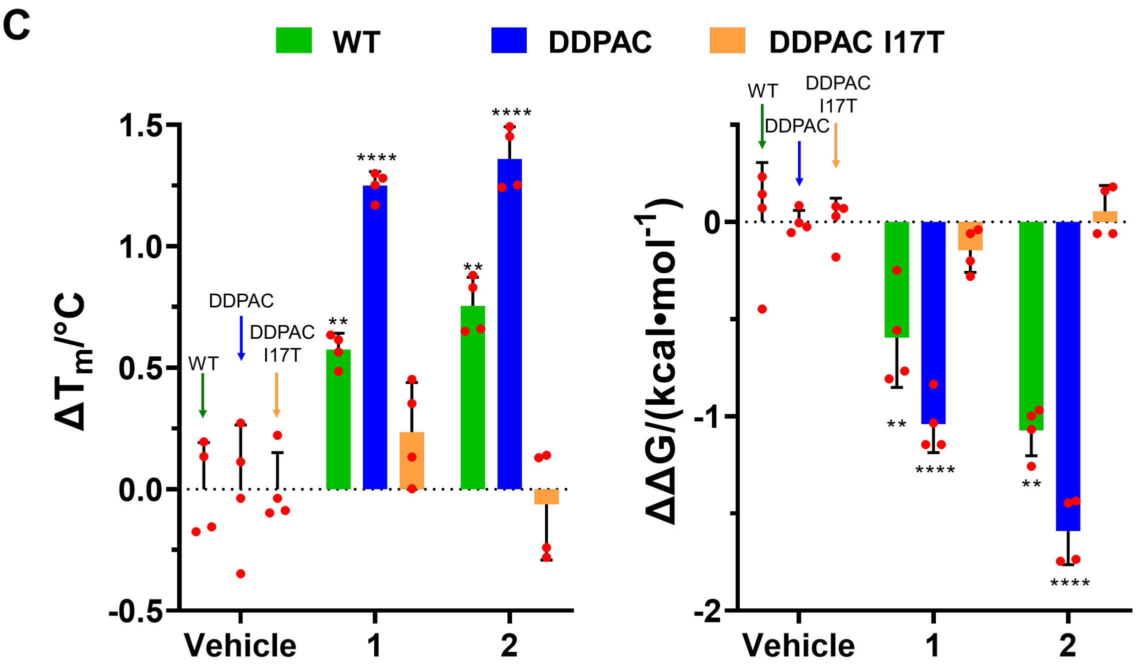

**Figure S11. The effects of 1 and 2 on the thermodynamic stability of tau RNA constructs.** (A) Representative melting curves of WT, DDPAC, and DDPAC I17T RNAs upon addition of DMSO vehicle, **1** and **2** (n = 4). (B) Table of melting temperatures ($T_m$) and difference of free energies ($\Delta G°_{37}$) (n = 4). (C) Change in $T_m$ ($\Delta T_m$) and ΔG (ΔΔG) of WT, DDPAC, and DDPAC I17T RNAs upon addition of **1** and **2** compared to DMSO vehicle (n = 4). **, $p < 0.01$; and ****, $p < 0.0001$, as determined by two-tailed Student t-test.

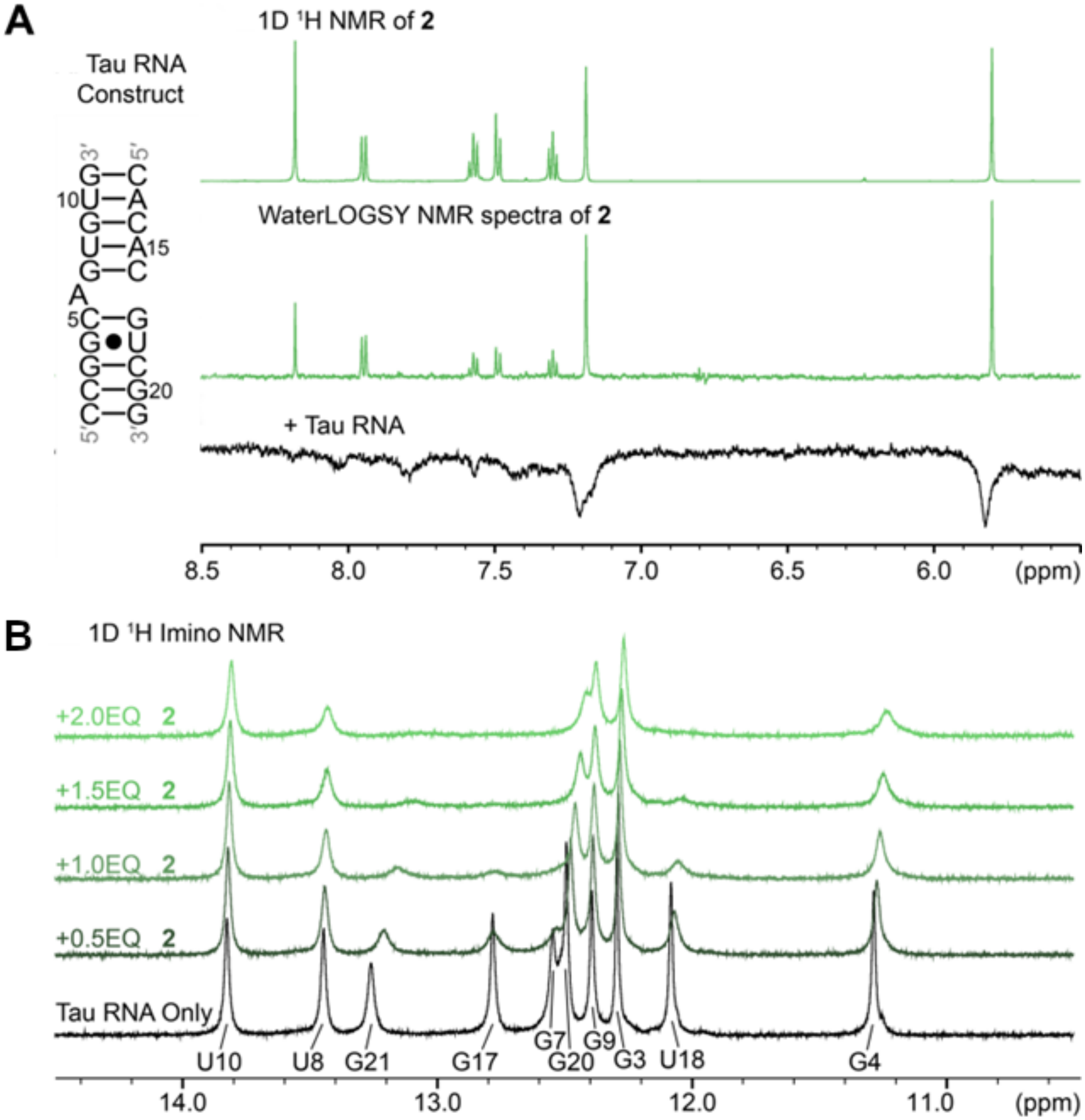

**Figure S12. 1D NMR spectral analysis of the interaction of 2 with the WT tau duplex.** (A) Secondary structure of the tau RNA duplex construct used for NMR studies (left). (Right, top) 1D $^1$H NMR spectrum of **2** (300 μM) at 298 K, focusing on the aromatic region. (Right, middle) WaterLOGSY NMR spectrum for **2** alone (300 μM; green), acquired in 95% $H_2O$ and 5% $D_2O$ at 298 K. (Right, bottom) WaterLOGSY NMR spectrum for **2** (300 μM) and the WT tau duplex (10 μM) (black), affording a compound:RNA ratio 30:1, showing the same aromatic region. (B) 1D $^1$H NMR spectrum of exchangeable imino protons in the WT tau duplex (50 μM), acquired at 10°C in the presence and absence of varying equivalents of **2**. The imino proton spectrum in the absence of **2** shows base pairing of the RNA duplex in black and allows the assignment of the bases within the RNA duplex.

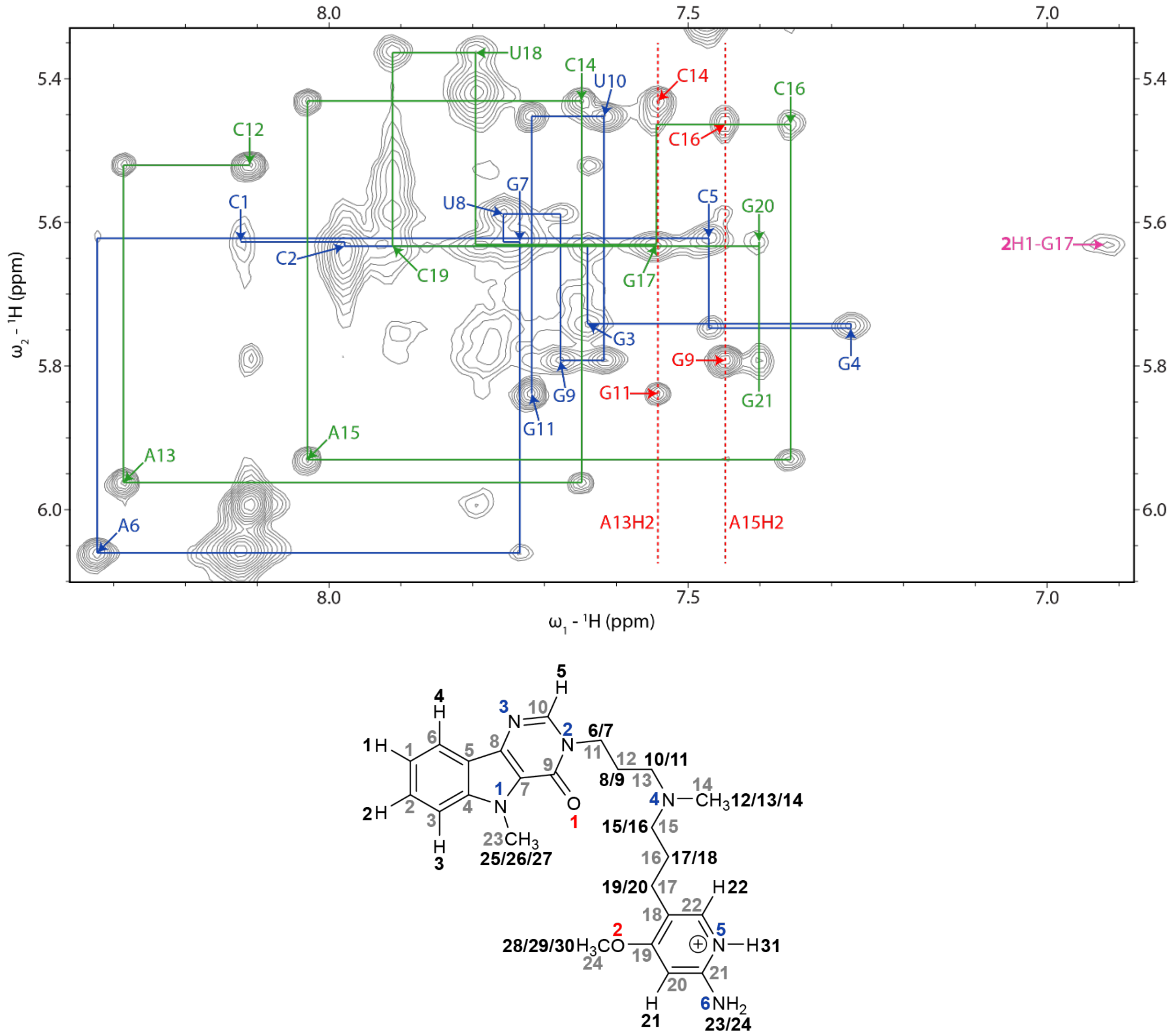

**Figure S13. 2D NMR spectral analysis of 2 and WT tau duplex.** Top, 2D $^1H$-$^1H$ $D_2O$ NOESY NMR spectrum of the tau duplex RNA and **2**, focusing on the non-exchangeable aromatic-sugar region. The NOESY walk is highlighted with lines, with distinct duplex strands shown in blue and green; AH2s are highlighted with red dashed lines, and intramolecular NOEs are shown in pink. The resulting lowest energy state of **2** bound to the RNA yielded an orientation in accordance with the single NOE observed between H5/H6 of the compound to G17H1′ (9 out of 20 different orientations created by docking). Bottom, in the chemical structure of **2** the numbering scheme for NMR study is shown. Black numbers correspond to hydrogens, gray correspond to carbons, blue correspond to nitrogens and red corresponds to oxygens.

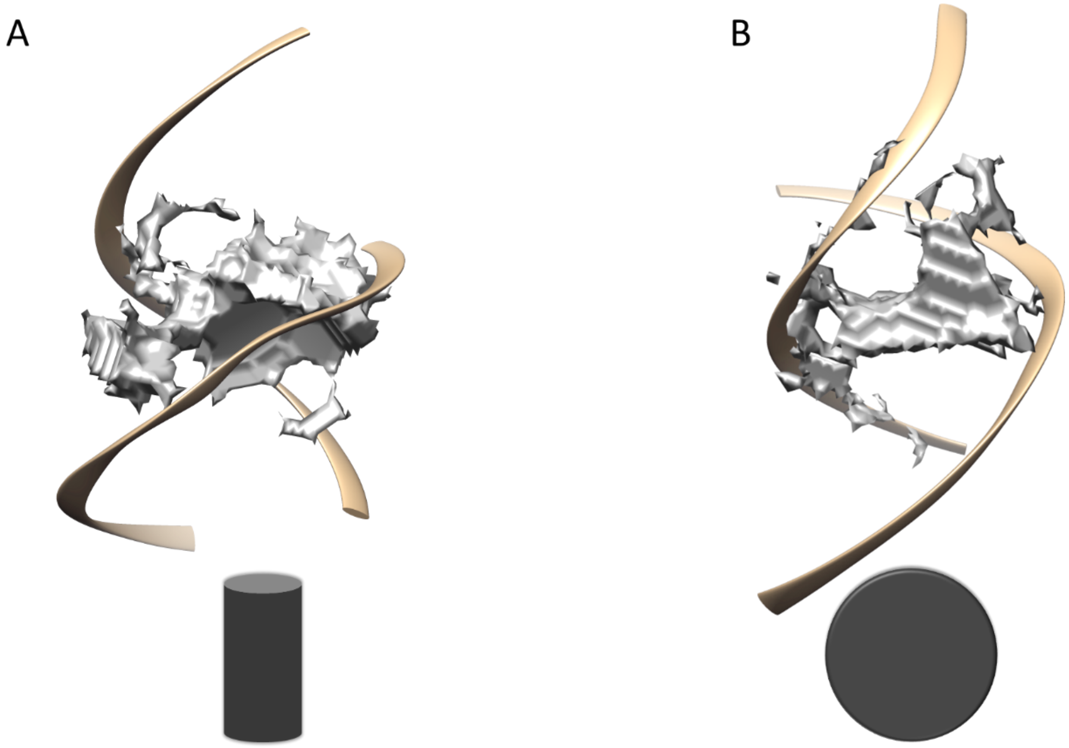

**Figure S14. Topology of the binding pocket induced by 2 (A) and 1 (B).** The binding pocket provided by the flipped-out A-bulge adopts a rod-shaped topology induced by **2** (A) and sphere-shape topology induced by **1** (B). Note: the A-bulge is positioned behind the volumetric representation of the binding pocket (grey). The RNA backbone is shown in cartoon-like mode.

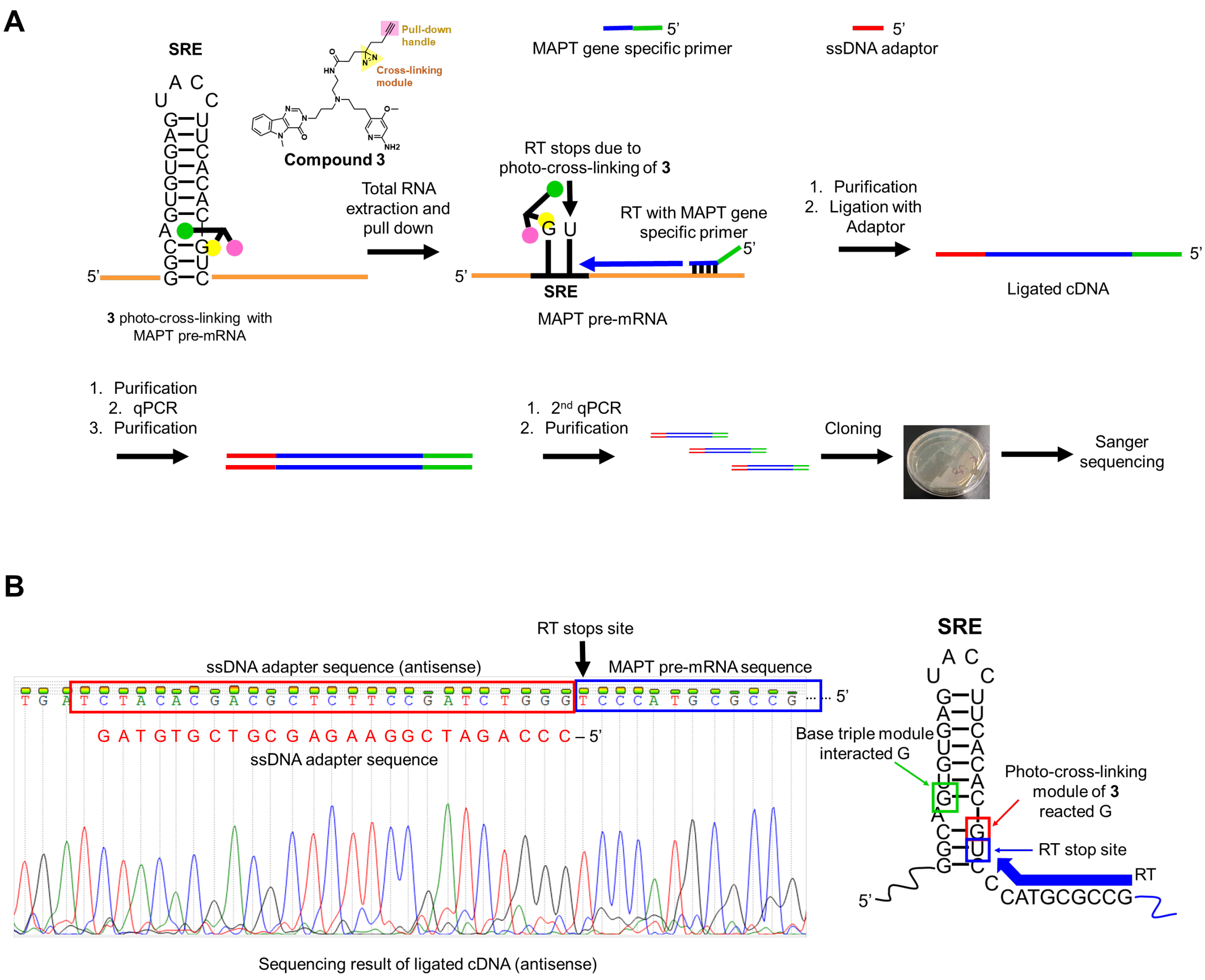

**Figure S15. Chem-CLIP-Map to map the cross-linking site of 3 in cells.** (A) Scheme of the approach to identify the binding site of the Chem-CLIP probe **3** via photo-cross-linking. (B) Representative Sanger sequencing results to identify the nucleotide cross-linked to **3**, indicated with an arrow; that is, where reverse transcription is halted (left). Secondary structure of the SRE annotated with the **2** binding site, including the G that participates in a base triple interaction (green box), the nucleotide where cross-linking occurs (red box), and the corresponding RT stop (blue box) (right). In total, 40 ampicillin-resistant colonies were selected and subjected to Sanger sequencing, 31 of which showed the indicated RT stop site (blue box).

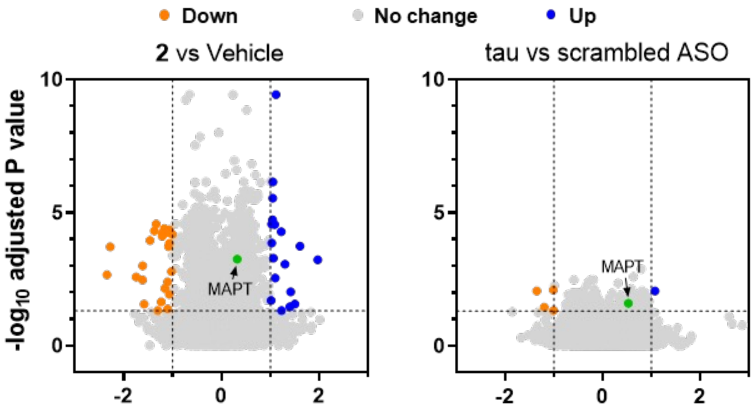

**Figure S16. Effect of 2 and an ASO directing *MAPT* splicing towards the 3R isoform in LAN5 cells, as determined by RNA-seq.** Volcano plot showing transcriptome-wide changes induced by **2** (left, 1.5 μM) as compared to vehicle (0.1% (v/v) DMSO) and tau ASO (right, 0.5 μM) as compared to the same concentration of a scrambled ASO control (n = 6 biological replicates for DMSO vehicle and **2**; n = 4 biological replicates for tau and scrambled ASO). Dotted lines represent absolute fold change >2 and a P-value < 0.05.

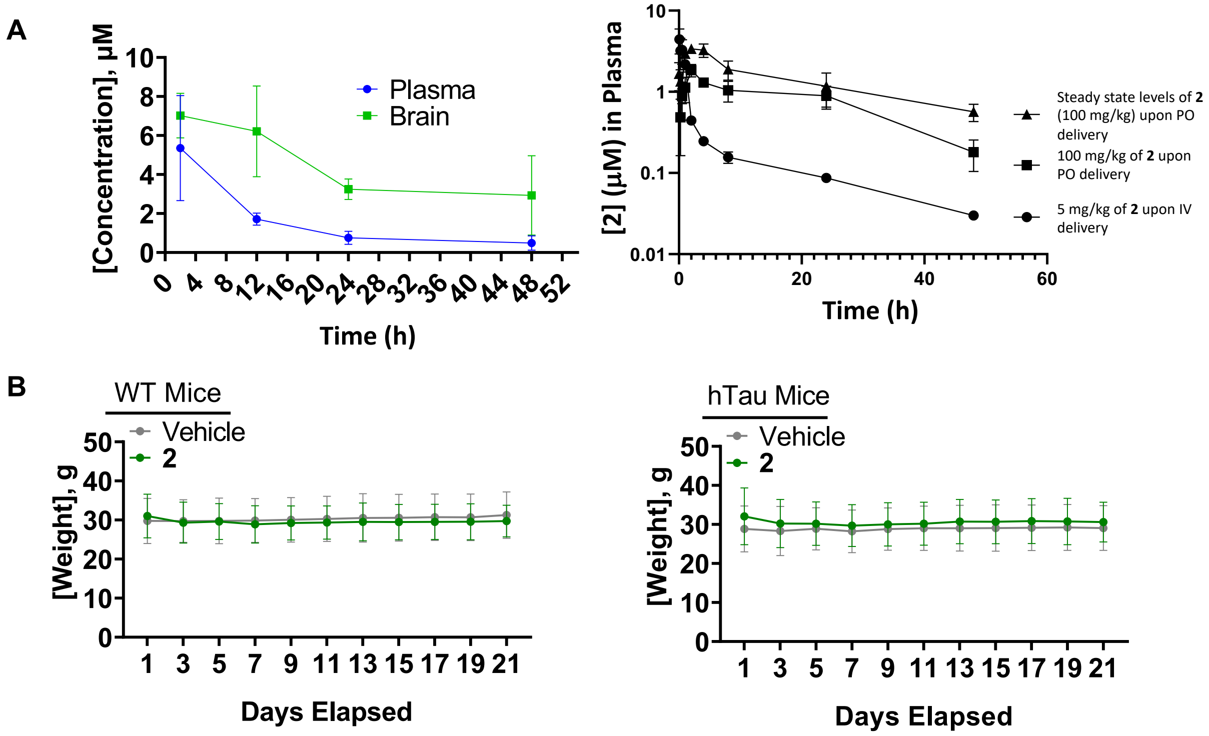

**Figure S17. Drug metabolism and pharmacokinetics (DMPK) analysis of 2 in C57BL/6 mice and the effect of 2 on the weight of WT and htau mice upon treatment.** (A) Left: Plasma and brain exposure after oral administration of a single dose of **2** (100 mg/kg), at the indicate time points (n = 3 mice per time point). These data show that equilibrium has not been fully reached at the 1 h time point as the fold difference between the concentration in brain vs. plasma is less than at longer time points. Right: Pharmacokinetics of **2** under: (i) steady state conditions (triangles; 100 mg/kg, *p.o.* delivery on Day 1 at 7 am and 5 pm, Day 2 at 7 am, and Day 3 at 7am followed by collection of data at the indicated time points); upon single delivery of 100 mg/kg *p.o* on Day 3 at 7 am followed by collection of data at the indicated time points (squares), and upon delivery of 5 mg/kg *i.v.* on Day 3 at 7 am followed by collection of data at the indicated time points (circles); the concentration of **2** was measured in plasma. Steady state studies were completed to understand plasma concentrations in therapeutic efficacy studies, in which htau transgenic mice were dosed q.o.d. (B) WT mice or htau mice treated with **2** over 3 weeks have similar weights as vehicle-treated mice (n = 7). Mice were orally administered (*p.o.*) **2** at a dose of 100 mg/kg q.o.d..

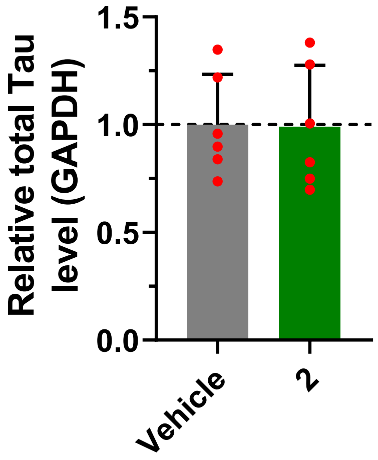

**Figure S18. Effect of 2 on total *MAPT* levels in htau mice treated with vehicle or 2 (n = 6), as determined by RT-qPCR.** Note **2** reduced the 4R/3R *MAPT* ratio in hTau mice (Figure 4A).

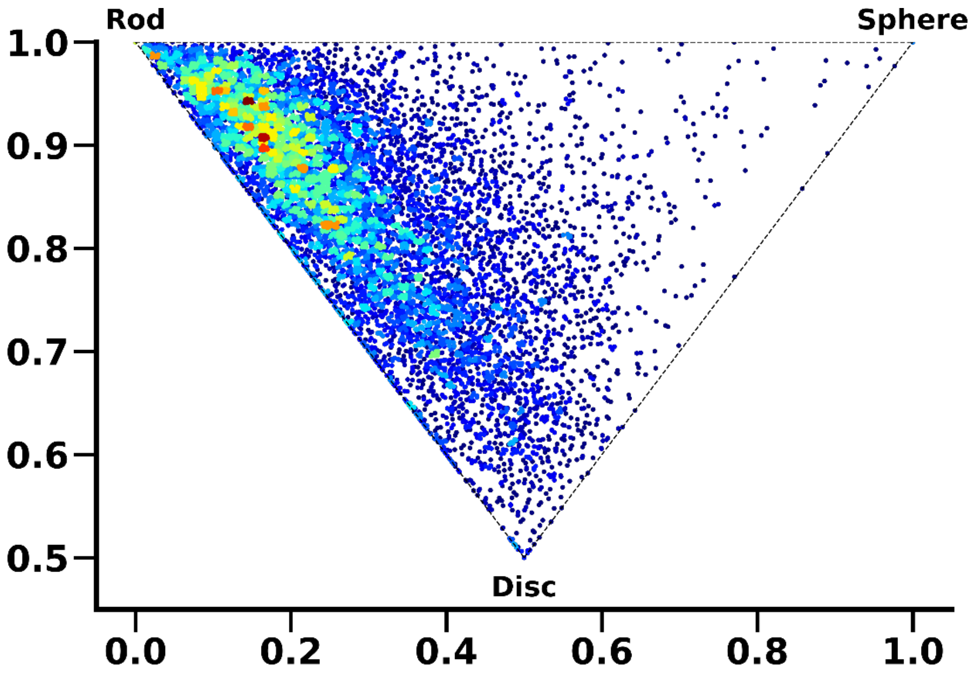

**Figure S19. Shape calculations of compounds found in DrugBank[38] using the normalized principles moment of inertia (NPR1, NPR2).** Based on the NPR values compound shape is categorized as rod-, disc- or sphere-like.[39]

**Table S1**. Calculation of CNS MPO scores and binding energy ($\Delta G°_{37}$) of compounds used in this study.

| Compound | CNS MPO Score[a] | LogP | CNS MPO Sub-score | LogD | CNS MPO Sub-score | MW (g/mol) | CNS MPO Sub-score | TPSA (Å$^2$) | CNS MPO Sub-score | HBD | CNS MPO Sub-score | p$K_a$ | CNS MPO Sub-score | Binding energy (kcal/mol)[b] |
|---|---|---|---|---|---|---|---|---|---|---|---|---|---|---|
| **1** | 4.72 | 2.34 | 1.0 | 0.17 | 1.0 | 368.48 | 0.94 | 60.93 | 1.0 | 1 | 0.75 | 9.93 | 0.03 | -8.91 |
| **2** (**1** analog) | 4.30 | 2.12 | 1.0 | -0.65 | 1.0 | 434.54 | 0.47 | 88.98 | 1.0 | 2 | 0.50 | 9.34 | 0.33 | -9.91 |
| **M1** | 4.0 | 2.46 | 1.0 | 0.0 | 1.0 | 306.37 | 1.0 | 84.02 | 1.0 | 4 | 0.0 | 11.35 | 0.0 | -6.39 |
| **S1** | 3.65 | 2.18 | 1.0 | 1.88 | 1.0 | 408.53 | 0.65 | 120.54 | 0.0 | 6 | 0.0 | 7.32 | 1.0 | -11.42 |
| **S2** (**S1** analog) | 2.74 | 2.89 | 1.0 | 2.52 | 0.74 | 528.68 | 0.0 | 145.46 | 0.0 | 7 | 0.0 | 7.41 | 1.0 | -12.97 |
| **S3** (**S1** analog) | 2.96 | 2.16 | 1.0 | 1.28 | 1.0 | 629.79 | 0.0 | 197.78 | 0.0 | 9 | 0.0 | 8.08 | 0.96 | -13.57 |
| **S4** (**S1** analog) | 2.71 | 3.03 | 0.99 | 1.72 | 1.0 | 558.71 | 0.0 | 168.68 | 0.0 | 8 | 0.0 | 8.56 | 0.72 | -12.76 |
| **S5** (**S1** analog) | 2.14 | 3.87 | 0.56 | 1.56 | 1.0 | 708.89 | 0.0 | 216.82 | 0.0 | 10 | 0.0 | 8.85 | 0.58 | -12.29 |
| **S6** (**S1** analog) | 2.12 | 3.60 | 0.70 | 3.16 | 0.42 | 648.84 | 0.0 | 170.38 | 0.0 | 8 | 0.0 | 7.49 | 1.0 | -12.35 |
| **S7** (**1** analog) | 4.46 | 2.42 | 1.0 | -0.25 | 1.0 | 448.57 | 0.37 | 74.99 | 1.0 | 1 | 0.75 | 9.32 | 0.34 | -9.15 |
| **S8** (**1** analog) | 4.31 | 1.97 | 1.0 | -0.48 | 1.0 | 434.54 | 0.47 | 85.85 | 1.0 | 2 | 0.50 | 9.31 | 0.34 | -9.19 |

[a]CNS-MPO scores were calculated using the implementation of ChemAxon. LogP: partition coefficient, LogD: distribution coefficient, MW: molecular weight, TPSA: Topological Polar Surface Area, HBD: hydrogen bond donor, pKa: most basic center. [b]lowest free energy docking pose extracted from Autodock-GPU.

**Table S2.** Free energy calculations of the first 100 clusters of **2**. Rism was used to calculate the free energy of the bound states of the first 100 clusters with a minimum of 100 structures.[40]

| Cluster # | $\Delta G_{RISM}$ | Cluster # | $\Delta G_{RISM}$ | Cluster # | $\Delta G_{RISM}$ |
|---|---|---|---|---|---|
| 16 | -20.0385 | 36 | -15.5342 | 73 | -12.6870 |
| 3 | -19.1309 | 77 | -15.5275 | 80 | -12.2771 |
| 41 | -18.9802 | 78 | -15.5185 | 90 | -12.2113 |
| 21 | -18.9724 | 29 | -15.4620 | 52 | -12.0081 |
| 6 | -18.7596 | 14 | -15.4530 | 54 | -11.9202 |
| 48 | -18.3165 | 51 | -15.3080 | 33 | -11.7872 |
| 10 | -18.1301 | 15 | -15.0693 | 99 | -11.7288 |
| 4 | -18.0512 | 0 | -15.0174 | 62 | -11.3820 |
| 100 | -17.9325 | 93 | -14.9522 | 55 | -11.1613 |
| 32 | -17.7566 | 75 | -14.9455 | 94 | -11.0715 |
| 8 | -17.5080 | 37 | -14.7530 | 83 | -10.9736 |
| 1 | -17.4271 | 40 | -14.7144 | 35 | -10.9243 |
| 17 | -17.3956 | 88 | -14.5433 | 64 | -10.1825 |
| 26 | -17.3750 | 13 | -14.3551 | 72 | -9.8956 |
| 87 | -17.3603 | 59 | -14.3140 | 18 | -9.5742 |
| 43 | -17.3524 | 31 | -14.3108 | 81 | -9.5690 |
| 95 | -17.2698 | 71 | -14.3096 | 67 | -9.5645 |
| 25 | -17.2129 | 49 | -14.0587 | 65 | -9.5006 |
| 38 | -17.1764 | 79 | -13.9413 | 39 | -9.3982 |
| 5 | -17.0453 | 28 | -13.9382 | 84 | -9.3144 |
| 7 | -16.8494 | 58 | -13.8899 | 76 | -9.2738 |
| 23 | -16.8208 | 46 | -13.8798 | 69 | -9.2691 |
| 34 | -16.4681 | 86 | -13.7486 | 45 | -9.2490 |
| 56 | -16.4535 | 66 | -13.5303 | 24 | -9.2360 |
| 9 | -16.3005 | 27 | -13.1813 | 96 | -8.8909 |
| 2 | -16.2124 | 19 | -13.0964 | 57 | -8.8818 |
| 11 | -16.2034 | 44 | -13.0564 | 74 | -8.7341 |
| 82 | -16.1283 | 47 | -13.0385 | 20 | -8.6791 |
| 50 | -15.8621 | 70 | -12.8535 | 12 | -8.6702 |
| 98 | -15.6051 | 60 | -12.8524 | 30 | -8.6700 |
| 36 | -15.5342 | 42 | -12.8402 | 68 | -8.6379 |
| 77 | -15.5275 | | | 22 | -8.5919 |
| 78 | -15.5185 | | | 61 | -8.5628 |
| 29 | -15.4620 | | | 97 | -8.5057 |
| 14 | -15.4530 | | | 53 | -8.4107 |
| 51 | -15.3080 | | | 91 | -8.3540 |
| 15 | -15.0693 | | | 85 | -8.1336 |
| 0 | -15.0174 | | | 63 | -8.0024 |
| 93 | -14.9522 | | | 92 | -7.3036 |
| 75 | -14.9455 | | | 89 | -4.8459 |
| 37 | -14.7530 | | | | |
| 40 | -14.7144 | | | | |
| 88 | -14.5433 | | | | |
| 13 | -14.3551 | | | | |
| 59 | -14.3140 | | | | |

| | |
|---|---|
| 31<br>71<br>49<br>79<br>28<br>58<br>46<br>86<br>66<br>27<br>19<br>44<br>47<br>70<br>60<br>42 | -14.3108<br>-14.3096<br>-14.0587<br>-13.9413<br>-13.9382<br>-13.8899<br>-13.8798<br>-13.7486<br>-13.5303<br>-13.1813<br>-13.0964<br>-13.0564<br>-13.0385<br>-12.8535<br>-12.8524<br>-12.8402 |

**Table S3.** Sequences of primers used in these studies. F denotes forward primer and R denotes reverse primer

| Purpose | F/R | Sequence |
|---|---|---|
| 4R tau mRNA in transfected HeLa cells | F | 5′-GAGGCGGGAAGGTGCAGATAATTAATAAGA-3′ |
| 3R tau mRNA in transfected HeLa cells | F | 5′-CAGCCGGGAGGCGGGAAGGTGCAAATAG-3′ |
| 4R and 3R tau mRNA in transfected HeLa cells | R | 5′-GCCTTATGCAGTTGCTCTCC-3′ |
| 4R tau in LAN5 cells | F | 5′-GAGGCGGGAAGGTGCAGATAATTAATAA-3′ |
| 4R tau in LAN5 cells | R | 5′-CTGGTTTATGATGGATGTTGCC-3′ |
| 3R tau in LAN5 cells and mouse | F | 5′-GAAGAATGTCAAGTCCAAGATCGG-3′ |
| 3R tau in LAN5 cells and mouse | R | 5′-GACTATTTGCACCTTCCCGC-3′ |
| Tau pre-mRNA in Chem-CLIP experiments | F | 5′-GGAAGTGGTGTGAGTGCGTACAC-3′ |
| Tau pre-mRNA in Chem-CLIP experiments | R | 5′-CACCTTCAGCCCAACTTCCAATG-3′ |
| Total tau in mouse | F | 5’-AGAAGCAGGCATTGGAGAC |
| Total tau in mouse | R | 5’-TCTTCGTTTTACCATCAGCC |
| GAPDH in mouse | F | 5’-TGCCCCCATGTTGTGATG |
| GAPDH in mouse | R | 5’-TGTGGTCATGAGCCCTTCC |
| 4R tau in mouse | F | 5′-CACTGAGAACCTGAAGCACC-3′ |
| 4R tau in mouse | R | 5′-GGACGTTGCTAAGATCCAGCT-3′ |
| GAPDH | F | 5′-GGCAAATTCAACGGCACAGT-3′ |
| GAPDH | R | 5′-GGGTCTCGCTCCTGGAAGAT-3′ |
| β-actin | F | 5′-CATGTACGTTGCTATCCAGGC-3′ |
| β-actin | R | 5′-CTCCTTAATGTCACGCACGAT-3′ |
| 18S rRNA | F | 5′-GTAACCCGTTGAACCCCATT-3′ |
| 18S rRNA | R | 5′-TCCAATCGGTAGTAGCG-3′ |
| htau mice Genotyping[41] | F | 5’-ACTTTGAACCAGGATGGCTGAGCCC-3’ |
| htau mice Genotyping[41] | R | 5’-CTGTGCATGGCTGTCCACTAA CCT T-3’ |

### Synthetic Experimental Procedures

**Abbreviations.** AcOH, acetic acid; Boc, tert-butyloxycarbonyl; Cbz, benzyloxycarbonyl; DCM, dichloromethane; DIEA, diisopropylethylamine; DMF, *N*, *N*-dimethylformamide; HATU, hexafluorophosphate azabenzotriazole tetramethyl uronium; HPLC, high performance liquid chromatography; MeOH, methanol; NIS, *N*-Iodosuccinimide; TFA, trifluoroacetic acid; TLC, thin layer chromatography

**General methods.** Reagents and solvents were purchased from commercial sources and used without further purification. Reactions were monitored by thin layer chromatography (TLC, Agela Technologies) or by LC-MS. Bands on TLC were visualized under UV light (254 nm). Compound **2** was purified by Isolera One Flash Chromatography System (Biotage) using pre-packed C18 column (spherical 20-35 μm, Agela Technologies). Preparative HPLC purification for **3** was performed by HPLC (Waters 2489 and 1525) using a SunFire® Prep C18 OBD™ 5 μm column (19 × 150 mm) with a 5 mL/min flow. NMR spectra were collected on a 400 UltraShield™ (Bruker) (400 MHz for $^1$H and 100 MHz for $^{13}$C) or Ascend™ 600 (Bruker) (600 MHz for $^1$H and 150 MHz for $^{13}$C). Chemical shifts are reported in ppm relative to tetramethylsilane (TMS) for $^1$H and residual solvent for $^{13}$C as internal standards. Coupling constants (J values) are expressed in Hz. High resolution mass spectra of all compounds were recorded on a 4800 Plus MALDI TOF/TOF Analyzer (Applied Biosystems) with α-cyano-4-hydroxycinnamic acid matrix and TOF/TOF Calibration Mixture (AB Sciex Pte. Ltd) or on an Agilent 1260 Infinity LC system coupled to an Agilent 6230 TOF (HR-ESI) with a Poroshell 120 EC-C18 column (Agilent, 50 mm × 4.6 mm, 2.7 μm).

The preparation, synthesis, and characterization of **M1**, **1**, and **4** have been previously reported.[18]

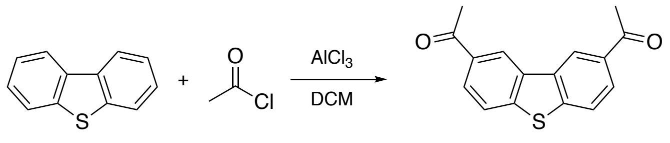

**F1** **F2**

**Scheme 7:** Synthesis of **F2**.

**Synthesis of 1,1'-(dibenzo[*b*,*d*]thiophene-2,8-diyl)bis(ethan-1-one) (F2)**: To a solution of $AlCl_3$ (32 g, 246 mmol) in dichloromethane (120 mL) was added acetyl chloride (22 mL, 307 mmol) at 0 °C. The solution was stirred for 10 min at 0 °C. A solution of dibenzothiophene **F1** (11.3 g, 61 mmol) in dichloromethane (50 mL) was added to the solution dropwise at 0 ˚C. The ice bath was removed, and the mixture was stirred at room temperature for 2 hours. The mixture was poured into ice bath and stirred for 2 hours. The product was extracted by dichloromethane, and the organic layer was washed with water until the aqueous layer was neutral. The organic solution was washed with brine and dried over sodium sulfate. The product was purified by silica gel column chromatography (dichloromethane 100%) to give diketone **F2** as a white solid (2.5 g, 9.5 mmol, 15%). $^1$H NMR (400 MHz, DMSO-$d_6$): $\delta$ 9.20 (d, $J$ = 1.6, 2H), 8.23 (d, $J$ = 8.4, 2H), 8.11 (dd, $J$ = 8.4, 1.7, 2H), 2.77 (s, 6H).

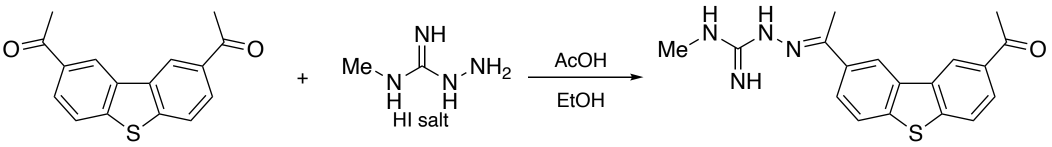

**F2** **F3**

**Scheme 8:** Synthesis of **F3**.

**Synthesis of (*E*)-2-(1-(8-acetyldibenzo[*b*,*d*]thiophen-2-yl)ethylidene)-*N*-methylhydrazine-1-carboximidamide (F3)**: A mixture of **F2** (268 mg, 1 mmol) and 1-amino-2-methylguanidinium iodide (216 mg, 1 mmol) in ethanol (5 mL) and acetic acid (5 mL) was stirred for 6 h at 80 °C. After evaporation of solvents, the crude was washed with water, saturated aqueous $NaHCO_3$, water, methanol, dichloromethane, and ether to give **F3** as a white solid (250 mg, 0.74 mmol, 74%). $^1$H NMR (400 MHz, DMSO-$d_6$): $\delta$ 10.40 (s, 1H), 9.13 (d, $J$ = 1.3, 1H), 9.02 (d, $J$ = 1.5, 1H), 8.33 (dd, $J$ = 8.6, 1.8, 1H), 8.21 (d, $J$ = 8.4, 1H), 8.15 (d, $J$ = 8.6, 1H), 8.11 (dd, $J$ = 8.4, 1.7, 1H), 8.08-7.66 (3H), 2.97 (m, 3H), 2.76 (s, 3H), 2.49 (s, 3H, overlapped with solvent peak).

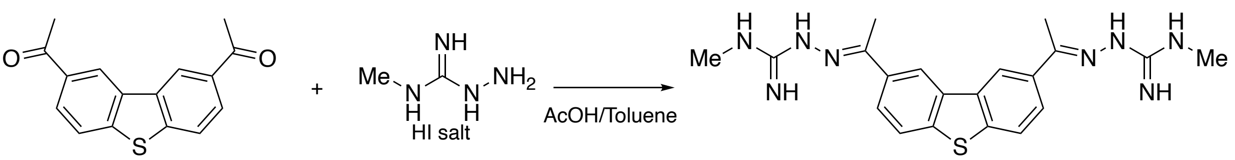

**F2** **S1**

**Scheme 9:** Synthesis of **S1**.

**Synthesis of (2*E*,2'*E*)-2,2'-(dibenzo[*b*,*d*]thiophene-2,8-diylbis(ethan-1-yl-1-ylidene))bis(*N*-methylhydrazine-1-carboximidamide) (S1)**: A mixture of **F2** (26.8 mg, 0.1 mmol) and 1-amino-2-methylguanidinium iodide (21.6 mg, 0.1 mmol) in toluene (1 mL) and acetic acid (1 mL) was stirred for 6 h at 80 °C. After evaporation of solvents, the crude was washed with water, saturated aqueous $NaHCO_3$, water, methanol, dichloromethane, and ether to give **S1** as a white solid (16%). $^1H$ NMR (400 MHz, $CD_3OD$): $\delta$ 8.75 (s, 2H), 8.13 (m, 2H), 7.91 (m, 2H), 3.00 (s, 6H), 2.51 (s, 6H).

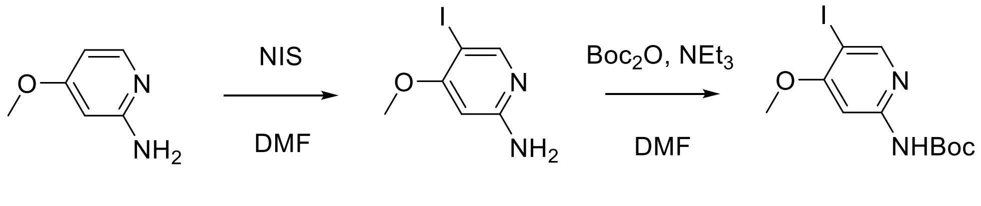

**A5** **A6** **A7**

**Scheme 3:** Synthesis of intermediate **A7**.

**Synthesis of tert-butyl (5-iodo-4-methoxypyridin-2-yl)carbamate (A7):** To a solution of 4-methoxypyridin-2-amine (**A5**) (25.0 g, 201 mmol) in DMF (400 mL), NIS (54.4 g, 225 mmol) was added at 0 °C. The mixture was stirred at room temperature overnight and then concentrated in vacuo. Next, DCM/ saturated $NaHCO_3$ (v/v, 400 mL) was added, and the resultant solids were separated by filtration. The resultant DCM/saturated $NaHCO_3$ mixture was extracted with DCM (400 mL × 3) and concentrated in vacuo. This material was washed with $H_2O$ (200 mL) to give 5-iodo-4-methoxypyridin-2-amine (**A6**) (33.0 g). To a suspension of **A6** (15.0 g, 60.0 mmol) and $NEt_3$ (13.4 mL, 66.0 mmol) in DCM (300 mL), $Boc_2O$ (14.4 g, 66.0 mmol) in DCM (150 mL) was added at 0 °C. The mixture was stirred at room temperature overnight. Additional reagents ($NEt_3$ (2.43 mL, 12.0 mmol) and $Boc_2O$ (2.62 g, 12.0 mmol)) were then added and the mixture was stirred at room temperature for 5 h. Then, 400 mL of silica gel was added, and the mixture was concentrated in vacuo. The mixture was purified by column chromatography (Silica gel 1.0 L, 10% ethyl acetate in DCM), and the resultant product was washed with hexane (8x v/w) and filtered to give tert-butyl (5-iodo-4-methoxypyridin-2-yl)carbamate (**A7**) (8.64 g, 27% (2 steps)). $^1H$ NMR (400 MHz, $CDCl_3$) δ 9.51 (s, 1H), 8.46 (s, 1H), 7.67 (s, 1H), 3.98 (s, 3H), 1.57 (s, 9H) $^{13}C$ NMR (150 MHz, DMSO-$d_6$) δ 165.2, 154.9, 154.8, 153.1, 96.3, 80.4, 77.3, 56.6, 28.5 (3C); HR-MS (ESI): Calcd for $C_{11}H_{16}IN_2O_3^+$ $[M+H]^+$; 351.0200; found, 351.0217.

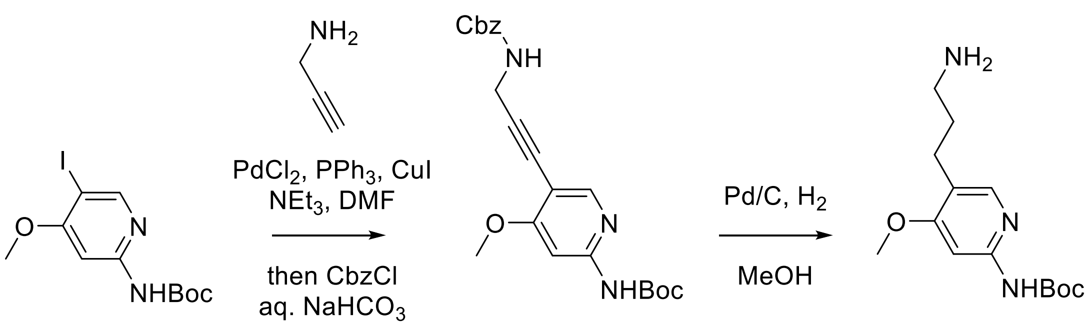

**A7** **A11** **A12**

**Scheme 6:** Synthesis of intermediate **A12**.

**Synthesis of tert-butyl (4-methoxy-5-(3-(amino)propyl)pyridin-2-yl)carbamate (A12):** A mixture of **A7** (1.50 g, 4.28 mmol), $PdCl_2$ (40.0 mg, 0.214 mmol, 5 mol %), $PPh_3$ (112 mg, 0.428 mmol, 10 mol %), CuI (81.6 mg, 0.428 mmol, 10 mol %), and propargylamine (823 mL, 12.9 mmol, 3 equiv.) in THF/DMF/$Et_3N$ (10:3:3, 16 mL) was stirred at 50 °C for 2 h. After cooling to room temperature, saturated $NaHCO_3$ solution (20 mL) and CbzCl (3.00 mL, 21.4 mmol, 5 equiv.) were added, and the mixture was vigorously stirred for 3 h. The resulting mixture was passed through a Celite pad and the filtrate was extracted with AcOEt (3 × 150 mL). The combined organic layer was washed with brine (50 mL), dried over $Na_2SO_4$ and concentrated in vacuo. The crude product was partially purified by column chromatography (Silica irregular 40-60 mm 80A 80 g, hexane/AcOEt 10–40%) to afford the coupling product **A11** (850 mg, impure), which was used without further purification. A mixture of **A11** (850 mg) and Pd/C (10%, 800 mg) in DCM/MeOH (v/v, 10 mL) was stirred at room temperature under $H_2$ (1 atm) for 1 h. The reaction mixture was filtered through a Celite pad and the filtrate was concentrated in vacuo. The crude product was purified by column chromatography (Sfär KP-Amino D 28 g, DCM/MeOH 0-20%) to afford the title amine **A12** (402 mg, 34% in 2 steps). $^1$H NMR (400 MHz, $CDCl_3$) δ 7.89 (s, 1H), 7.54 (s, 1H), 3.90 (s, 3H), 2.70 (t, J = 7.2 Hz, 2H), 2.55 (t, J = 7.4 Hz, 2H), 1.73-1.66 (m, 2H), 1.53 (s, 9H); $^{13}$C NMR (150 MHz, $CDCl_3$) δ 165.3, 152.7, 152.4, 147.6, 121.2, 94.5, 80.7, 55.3, 41.8, 33.7, 28.4, 24.6; HR-MS (ESI): Calcd for $C_{14}H_{24}N_3O_3^+$ $[M+H]^+$; 282.1812; found, 282.1776.

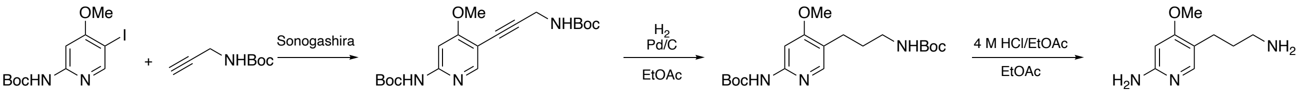

**A7** **F4** **F5** **F6**

**Scheme 10:** Synthesis of **F6**.

**Synthesis of 5-(3-aminopropyl)-4-methoxypyridin-2-amine (F6)**: **F5** was synthesized using the similar synthetic procedure for **A12** from **F4**.

**F4** (87% yield); $^1$H NMR (400 MHz, $CDCl_3$): $\delta$ 8.15 (s, 1H), 7.68-7.54 (2H), 4.80 (br, 1H), 4.20 (m, 2H), 3.95 (s, 3H), 1.57 (s, 3H), 1.53 (s, 3H).

**F5** (quantitative yield); $^1$H NMR (400 MHz, $CDCl_3$): $\delta$ 7.85 (s, 1H), 7.70-7.40 (2H), 4.59 (br, 1H), 3.90 (s, 3H), 3.12 (m, 2H), 2.53 (m, 2H), 1.72 (m, 2H), 1.53 (s, 9H), 1.45 (s, 9H).

A solution of **F5** (50 mg, 130 μmol) and 4 M HCl/EtOAc (1 mL) in ethyl acetate (5 mL) was stirred overnight at 50 °C. The solvents were evaporated to give amine **F6** as a yellow solid (40 mg, 157 μmol, quantitative). $^1$H NMR (400 MHz, $CD_3OD$): $\delta$= 7.62 (s, 1H), 6.41 (s, 1H), 4.00 (s, 3H), 2.96 (m, 2H), 2.59 (m, 2H), 1.90 (m, 2H).

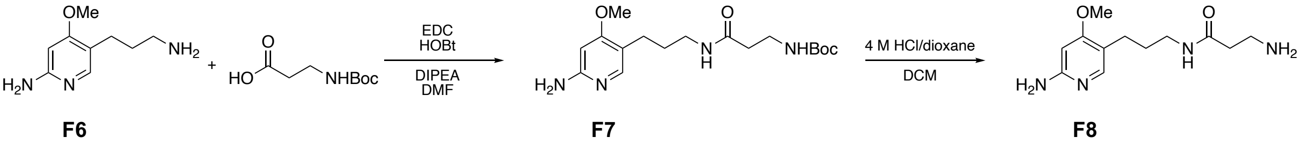

**F6** **F7** **F8**

**Scheme 11:** Synthesis of **F8**.

**Synthesis of 3-amino-N-(3-(6-amino-4-methoxypyridin-3-yl)propyl)propanamide (F8)**: A mixture of *N*-Boc-β-alanine (946 mg, 0.12 mmol), EDC (1.15 g, 0.12 mmol), HOBt (1.15g, 0.15 mmol) in DMF (0.5 mL) was stirred for 15 min at room temperature. To the mixture was added a solution of **F6** (907 mg, 0.1 mmol) and *N*,*N*-diisopropylethylamine (2.6 mL, 0.15 mmol) in DMF (0.5 mL), and the solution was stirred overnight at room temperature. The solvents were evaporated, and the crude material was extracted with DCM, washed with saturated aqueous $NaHCO_3$ and brine, and dried over $Na_2SO_4$. The product was purified by silica gel chromatography to give **F7** as a white solid (22 mg, 0.063 mmol, 63%). $^1$H NMR (400 MHz, $CD_3OD$): $\delta$ 7.53 (s, 1H), 6.17 (s, 1H), 3.84 (s, 3H), 3.20-3.10 (4H), 2.46 (m, 2H), 2.35 (m, 2H), 1.70 (m, 2H), 1.41 (s, 9H).

Boc deprotection reaction of **F7** was performed under the same condition from **F5** to **F6** as above to give **F8** as a colorless oil (508 mg, 1.6 mmol, quantitative). $^1$H NMR (400 MHz, $CD_3OD$): $\delta$ 7.60 (s, 1H), 6.40 (s, 1H), 3.98 (s, 3H), 3.26-3.13 (4H), 2.62 (m, 2H), 2.52 (m, 2H), 1.75 (m, 2H).

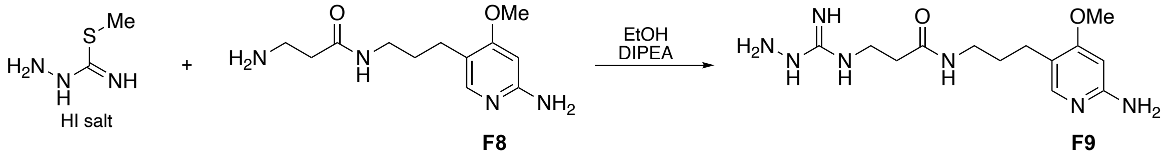

**Scheme 12:** Synthesis of **F9**.

**Synthesis of N-(3-(6-amino-4-methoxypyridin-3-yl)propyl)-3-(hydrazinecarboximidamido)-propanamide (F9)**: A mixture of methyl hydrazinecarbimidothioate hydroiodide (140 mg, 0.6 mmol), **F8** (97.5 mg, 0.3 mmol), and *N*,*N*-diisopropylethylamine (261 µL, 1.5 mmol) in ethanol (3 mL) was heated at 160 °C for 15 min by microwave. Concentration of solvent followed by HPLC purification gave **F9** as a colorless oil (TFA salt after HPLC) (83 mg, 154 µmol, 51%). $^1$H NMR (400 MHz, $CD_3OD$): $\delta$ 7.55 (s, 1H), 6.37 (s, 1H), 3.98 (s, 3H), 3.48 (m, 2H), 3.20 (m, 2H), 2.61-2.40 (4H), 1.73 (m, 2H).

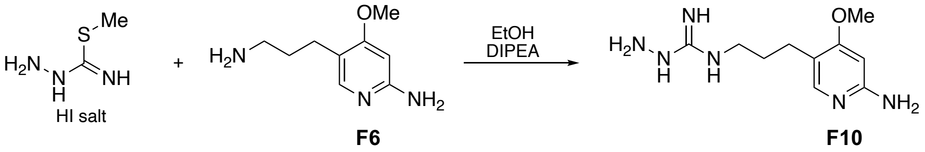

**Scheme 13:** Synthesis of **F10**.

**Synthesis of *N*-(3-(6-amino-4-methoxypyridin-3-yl)propyl)hydrazinecarboximidamide (F10)**: **F6** was performed under the same condition from **F8** to **F9** as above to give **F10** as a colorless oil (TFA salt after HPLC, 99 mg, 212 µmol, 71%). $^1$H NMR (400 MHz, $CD_3OD$): $\delta$ 7.56 (s, 1H), 6.38 (s, 1H), 3.99 (m, 3H), 3.22 (m, 2H), 2.55 (m, 2H), 1.85 (m, 2H)

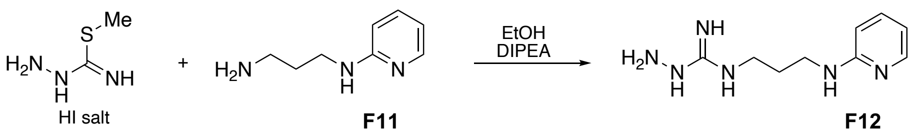

**Scheme 14:** Synthesis of **F12**.

**Synthesis of *N*-(3-(pyridin-2-ylamino)propyl)hydrazinecarboximidamide (F12)**: **F11** was performed under the same condition from **F8** to **F9** as above to give **F12** as a colorless oil (TFA salt after HPLC, 65 mg, 149 μmol, 30%). $^{1}$H NMR (400 MHz, $CD_3OD$): $\delta$= 7.93 (m, 1H), 7.86 (m, 1H), 7.07 (m, 1H), 6.91 (m, 1H), 3.52-3.33 (4H), 2.10-1.91 (2H).

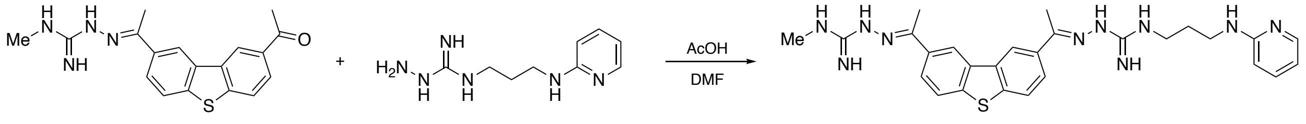

**F3** **F12** **S2**

**Scheme 15:** Synthesis of **S2**.

**Synthesis of (*E*)-*N*-methyl-2-(1-(8-((*E*)-1-(2-(*N*-(3-(pyridin-2-ylamino)propyl)carbamimidoyl)-hydrazineylidene)ethyl)dibenzo[b,d]thiophen-2-yl)ethylidene)hydrazine-1-carboximidamide (S2)**: A mixture of **F3** (13.5 mg, 40 μmol) and **F12** (19.2 mg, 44 μmol) in DMF (2 mL) and acetic acid (1 mL) was stirred overnight at 80 °C. After evaporation of solvents, the crude was purified by HPLC to give **S2** as a colorless oil (TFA salt after HPLC) (1.5 mg, 1.7 μmol, 4%). $^{1}$H NMR (400 MHz, $CD_3OD$): $\delta$ 8.87-8.76 (2H), 8.26-8.17 (2H), 8.00-7.94 (m, 2H), 7.91 (m, 1H), 7.85 (m, 1H), 7.08 (m, 1H), 6.89 (m, 1H), 3.61-3.46 (4H), 3.05 (s, 3H), 2.55 (s, 3H), 2.53 (s, 3H), 2.11 (m, 2H).

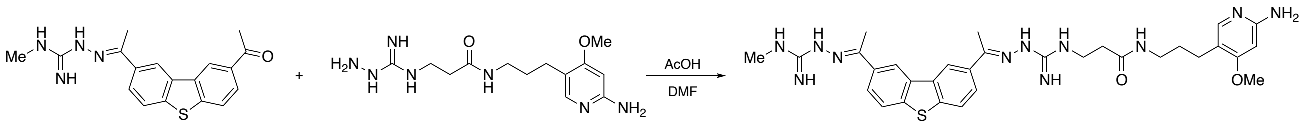

**F3** **F9** **S3**

**Scheme 16:** Synthesis of **S3**.

**Synthesis of *N*-(3-(6-amino-4-methoxypyridin-3-yl)propyl)-3-(2-((*E*)-1-(8-((*E*)-1-(2-(*N*-methyl-carbamimidoyl)hydrazineylidene)ethyl)dibenzo[b,d]thiophen-2-yl)ethylidene)hydrazine-1-carboximidamido)propenamide (S3)**: **S3** was synthesized using the similar synthetic procedure for **S2** as colorless oil (TFA salt after HPLC, 17 mg, 17.5 μmol, 35%). $^{1}$H NMR (400 MHz, $CD_3OD$): $\delta$ 8.89-8.78 (2H), 8.25-8.10 (2H), 8.02-7.91 (2H), 7.36 (s, 1H), 6.19 (s, 1H), 3.84 (s, 3H), 3.72 (m, 2H), 3.23 (m, 2H), 3.05 (s, 3H), 2.64 (m, 2H), 2.56-2.50 (6H), 2.39 (m, 2H), 1.67 (m, 2H).

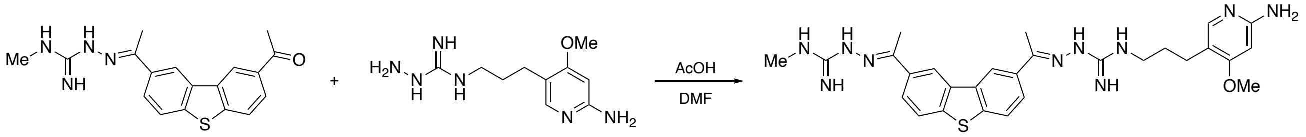

**F3** **F10** **S4**

**Scheme 17:** Synthesis of **S4**.

**Synthesis of (*E*)-N-(3-(6-amino-4-methoxypyridin-3-yl)propyl)-2-(1-(8-((*E*)-1-(2-(*N*-methyl-carbamimidoyl)hydrazineylidene)ethyl)dibenzo[b,d]thiophen-2-yl)ethylidene)hydrazine-1-carboximidamide (S4)**: **S4** was synthesized using the similar synthetic procedure for **S2** as colorless oil (TFA salt after HPLC, 7.8 mg, 8.7 μmol, 17%). $^{1}$H NMR (400 MHz, $CD_3OD$): $\delta$ 8.93-8.66 (2H), 8.32-8.06 (2H), 8.06-7.81 (2H), 7.59 (s, 1H), 6.37 (s, 1H), 3.96 (s, 3H), 3.44 (m, 2H), 3.04 (s, 3H), 2.63 (m, 2H), 1.94 (m, 2H). $^{13}$C NMR (100 MHz, $CD_3OD$): $\delta$= 171.0, 157.1, 142.8, 142.8, 136.9, 135.6, 135.0, 126.7, 126.7, 123.9, 121.6, 121.5, 119.4, 92.7, 57.4, 42.1, 29.1, 28.4, 24.9, 14.6, 14.5.

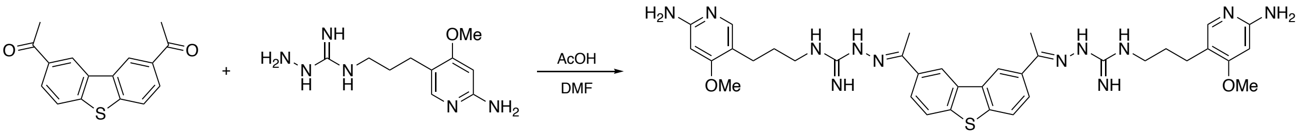

F2 F10 S5

**Scheme 18:** Synthesis of **S5**.

**Synthesis of (2*E*,2'*E*)-2,2'-(dibenzo[b,d]thiophene-2,8-diylbis(ethan-1-yl-1-ylidene))bis(*N*-(3-(6-amino-4-methoxypyridin-3-yl)propyl)hydrazine-1-carboximidamide) (S5)**: **S5** was synthesized using the similar synthetic procedure for **S2** as colorless oil (TFA salt after HPLC, 5 mg, 4.3 μmol, 17%). $^{1}$H NMR (400 MHz, $CD_3OD$): *δ* 8.81 (d, *J* = 1.5, 2H), 8.21 (dd, *J* = 8.6, 1.8, 2H), 7.97 (d, *J* = 8.6, 2H), 7.59 (s, 2H), 6.37 (s, 2H), 3.97 (s, 6H), 3.44 (m, 4H), 2.63 (m, 4H), 2.54 (s, 6H), 1.94 (m, 4H).

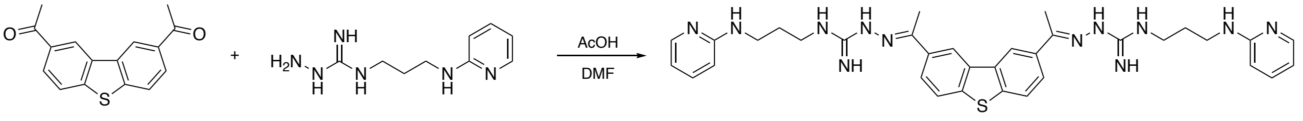

F2 F12 S6

**Scheme 19:** Synthesis of **S6**.

**Synthesis of (2*E*,2'*E*)-2,2'-(dibenzo[b,d]thiophene-2,8-diylbis(ethan-1-yl-1-ylidene))bis(*N*-(3-(pyridin-2-ylamino)propyl)hydrazine-1-carboximidamide) (S6)**: **S6** was synthesized using the similar synthetic procedure for **S2**. Colorless oil (TFA salt after HPLC) (5 mg, 4.5 μmol, 9%). $^{1}$H NMR (400 MHz, $CD_3OD$): *δ* 8.80 (d, *J* = 1.5, 2H), 8.21 (dd, *J* = 8.6, 1.8, 2H), 7.96 (d, *J* = 8.7, 2H), 7.90 (m, 2H), 7.85 (m, 2H), 7.06 (d, *J* = 9.1, 2H), 6.88 (m, 2H), 3.56 (t, *J* = 7.1, 4H), 3.50 (t, *J* = 7.0, 4H), 2.55 (s, 6H), 2.10 (m, 4H).

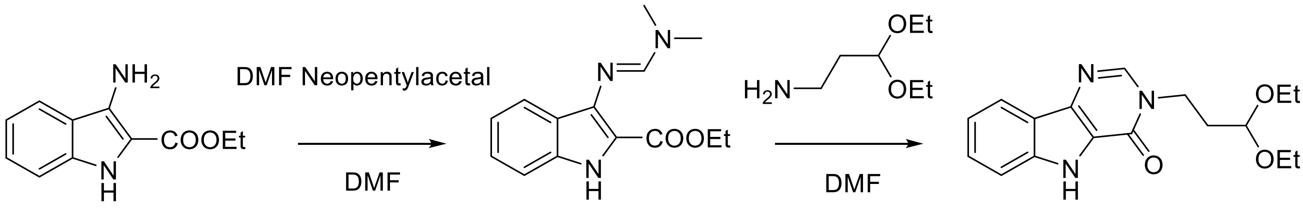

A1 A2 A3

**Scheme 1:** Synthesis of intermediate **A3**.

**Synthesis of 3-(3,3-diethoxypropyl)-3,5-dihydro-4H-pyrimido[5,4-b]indol-4-one (A3):** A mixture of ethyl 3-amino-1H-indole-2-carboxylate (**A1**) (5.00 g, 21.3 mmol) and Neopentylacetal (20.8 mL, 74.7 mmol) in DMF (75 mL) was stirred at 140 °C for 10 h. The reaction was concentrated in vacuo and azeotroped with toluene (3 × 5 mL) to give **A2**. This intermediate was added into 3,3-diethoxypropan-1-amine (20 mL), and the mixture was stirred at 100 °C for 3 h. After cooling to room temperature, the reaction mixture was concentrated in vacuo. To this mixture, $H_2O$ (20 mL) was added, and the mixture was filtered to give 3-(3,3-diethoxypropyl)-3,5-dihydro-4H-pyrimido[5,4-b]indol-4-one (**A3**) (6.00 g, 89% (2 steps)). The material was used in the next reaction without further purification. $^1$H NMR (400 MHz, MeOD) δ 8.26 (s, 1H), 8.10-8.07 (m, 1H), 7.60-7.55 (m, 1H), 7.52-7.47 (m, 1H), 7.29-7.24 (m, 1H), 4.64 (t, J = 5.2 Hz, 1H), 4.27 (t, J = 6.8 Hz, 2H), 3.68-3.60 (m, 2H), 3.55-3.44 (m, 2H), 2.19-2.10 (m, 2H), 1.11 (t, J = 7.0 Hz, 6H) $^{13}$C NMR (150 MHz, DMSO-$d_6$) δ 154.6, 144.9, 139.3, 138.0, 127.5, 122.3, 121.5, 120.7, 120.7, 113.3, 100.9, 61.2 (2C), 42.8, 33.2, 15.7 (2C); HR-MS (ESI): Calcd for $C_{17}H_{20}N_3O_3^-$ $[M-H]^-$; 314.1510; found, 314.1523.

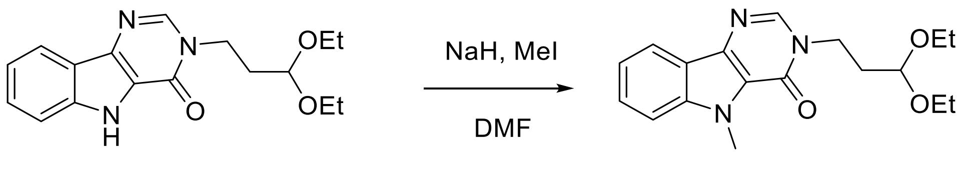

**A3** **A4**

**Scheme 2:** Synthesis of intermediate **A4**.

**Synthesis of 3-(3,3-diethoxypropyl)-5-methyl-3,5-dihydro-4H-pyrimido[5,4-b]indol-4-one (A4):** To a solution of **A3** (6.00 g, 19.0 mmol) in DMF (60.0 mL), NaH (1142 mg, 28.5 mmol) was added at 0 °C and the mixture was stirred at 0 °C for 10 min. Iodomethane (MeI, 3.53 mL, 38.1 mmol) was then added at 0 °C. The reaction mixture was stirred at 0 °C for 25 min, then $H_2O$ (100 mL) was added, and the mixture was extracted with MeOH/DCM (1/9, 3 x 100 mL) and concentrated in vacuo. The reaction mixture was purified by column chromatography (Agela Technologies, Silica, 20 g, 10% - 30% ethylacetate in hexane) to afford 3-(3,3-diethoxypropyl)-5-methyl-3,5-dihydro-4H-pyrimido[5,4-b]indol-4-one (**A4**) (4.98 g, 79 %). $^1$H NMR (400 MHz, $CDCl_3$) δ 8.15 (dd, J = 8.0, 0.90 Hz, 1H), 8.01 (s, 1H), 7.58-7.52 (m, 1H), 7.46 (d, J = 8.4 Hz, 1H), 7.34-7.29 (m, 1H), 4.57 (t, J = 5.4 Hz, 1H), 4.27 (s, 3H), 4.19 (t, J = 6.9 Hz, 2H), 3.72-3.62 (m, 2H), 3.55-3.46 (m, 2H), 2.20-2.14 (m, 2H), 1.20 (t, J = 7.1 Hz, 6H) $^{13}$C NMR (150 MHz, MeOD) δ 155.2, 144.4, 140.5, 137.3, 127.5, 121.3, 120.5, 120.2, 120.1, 109.9, 101.1, 61.5 (2C), 42.8, 32.5, 30.2, 14.1 (2C); HR-MS (ESI): Calcd for $C_{18}H_{24}N_3O_3^+$ $[M+H]^+$; 330.1812; found, 330.1827.

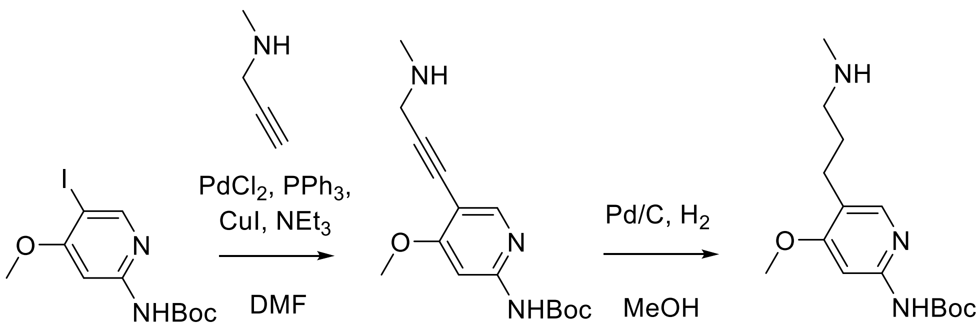

**A7** **A8** **A9**

**Scheme 4:** Synthesis of intermediate **A9**.

**Synthesis of tert-butyl (4-methoxy-5-(3-(methylamino)propyl)pyridin-2-yl)carbamate (A9):** A mixture of **A7** (3.00 g, 8.57 mmol), $PdCl_2$ (103 mg, 0.343 mmol), $PPh_3$ (180 mg, 0.685 mmol), CuI (131 mg, 0.685 mmol), $NEt_3$ (8.44 mL, 60.6 mmol) and N-methylprop-2-yn-1-amine (1.78 g, 25.7 mmol) in DMF (45 mL) was stirred at 50 °C for 1 h. After cooling to room temperature, saturated $NaHCO_3$ (100 mL) was added, and the mixture was extracted with ethyl acetate (3 × 100 mL) and concentrated in vacuo. The reaction mixture was purified by column chromatography (Biotage SNAP cartridge, KP-NH, 28g, 2% - 18% MeOH in DCM) to afford crude **A8**. Crude **A8** was further washed with hexane/$Et_2O$ (1/1, ×10 v/w) and lyophilized to dryness to give **A8** (2.13 g).

A mixture of **A8** (2.08 g, 7.12 mmol) and Pd/C (10%, 2.08 g) in MeOH (104 mL) was stirred at room temperature under $H_2$ (1 atm) for 1 h. The reaction was filtered through Celite and concentrated in vacuo. The resultant solid was washed with hexane/$Et_2O$ (1/1, x10v/w) and filtered to give tert-butyl (4-methoxy-5-(3-(methylamino)propyl)pyridin-2-yl)carbamate (**A9**) (1.56 g, 63% (2 steps)). $^1$H NMR (400 MHz, $CDCl_3$) δ 7.87 (s, 1H), 7.83 (br s, 1H), 7.52 (s, 1H), 3.90 (s, 3H), 2.60-2.51 (m, 4H), 2.43 (s, 3H), 1.78-1.69 (m, 2H), 1.53 (s, 9H) $^{13}$C NMR (150 MHz, MeOD) δ 165.5, 153.2, 152.7, 147.0, 121.1, 94.8, 80.3, 54.5, 50.8, 34.6, 28.8, 27.2, 24.5 (3C) ; HR-MS (ESI): Calcd for $C_{15}H_{26}N_3O_3^+$ [M+H]$^+$; 296.1969; found, 296.1978.

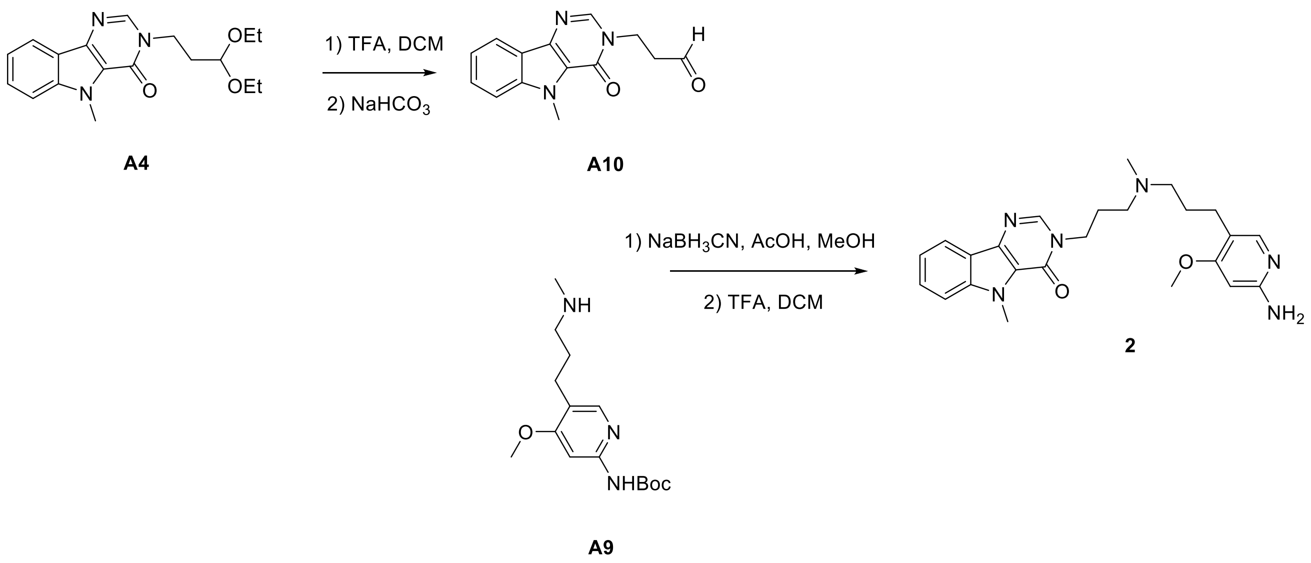

**Scheme 5:** Synthesis of **2**.

**Synthesis of 3-(3-((3-(6-amino-4-methoxypyridin-3-yl)propyl)(methyl)amino)propyl)-5-methyl-3,5-dihydro-4H-pyrimido[5,4-b]indol-4-one (2):** A mixture of **A4** (1.68 g, 5.10 mmol) in TFA/DCM (v/v, 25.7 mL) was stirred at room temperature for 30 min and then concentrated in vacuo. Next, saturated $NaHCO_3$ solution (10 mL) was added, and the mixture was extracted with DCM (3 x 20 mL) and concentrated in vacuo to give **A10**. To this product, **A9** (1.51 g, 5.10 mmol) was added in MeOH (51.0 mL), and the mixture was stirred at room temperature for 2 min. Subsequently $NaBH_3CN$ (641 mg, 10.2 mmol) and AcOH (584 μL, 10.2 mmol) were added, and the mixture was stirred at room temperature overnight. Saturated $NaHCO_3$ (20 mL) was added, and the mixture was extracted with DCM (3 × 20 mL) and concentrated in vacuo. The resultant material was dissolved in TFA/DCM (v/v, 25.7 mL), and the mixture was stirred at room temperature for 5 h and then concentrated in vacuo.

The crude material was purified by reverse-phase column chromatography (C18 column, 120 g, 2% - 100% MeOH + 0.1%TFA/$H_2O$ + 0.1%TFA, 80 ml/min). The fractions were collected and neutralized with saturated $NaHCO_3$ and extracted with DCM (3 × 50 mL). The resultant organic layer was dried over $MgSO_4$ and concentrated in vacuo. The material was then purified by column chromatography (Biotage SNAP cartridge, KP-NH, 28g, 100% ethyl acetate, 5 CV, 80 ml/min then 2% - 20% MeOH in DCM, 5 CV, 80 ml/min) to afford 3-(3-((3-(6-amino-4-methoxypyridin-3-yl)propyl)(methyl)amino)propyl)-5-methyl-3,5-dihydro-4H-pyrimido[5,4-b]indol-4-one (**2**) (1.00 g, 45% (3 steps)). $^1$H NMR (400 MHz, $CDCl_3$) δ 8.15 (d, J = 8.0 Hz, 1H), 8.06 (s, 1H), 7.71 (s, 1H), 7.58-7.52 (m, 1H), 7.46 (d, J = 8.4 Hz, 1H), 7.34-7.28 (m, 1H), 5.96 (s, 1H), 4.33 (br s, 2H), 4.26 (s, 3H), 4.16 (t, J = 6.9 Hz, 2H), 3.79 (s, 3H), 2.47 (t, J = 7.9 Hz, 2H), 2.42-2.32 (m, 4H), 2.21 (s, 3H), 2.03-1.95 (m, 2H), 1.75-1.65 (m, 2H) $^{13}$C NMR (150 MHz, $CDCl_3$) δ 165.4, 158.7, 155.5, 147.9, 143.7, 140.5, 138.2, 127.6, 121.9, 120.9, 120.9, 120.7, 117.6, 110.0, 90.3, 57.2, 54.9, 54.1, 44.7, 41.7, 31.3, 27.3, 26.8, 25.1 ; HR-MS (ESI): Calcd for $C_{24}H_{31}N_6O_2^+$ [M+H]$^+$; 435.2503; found, 435.2483.

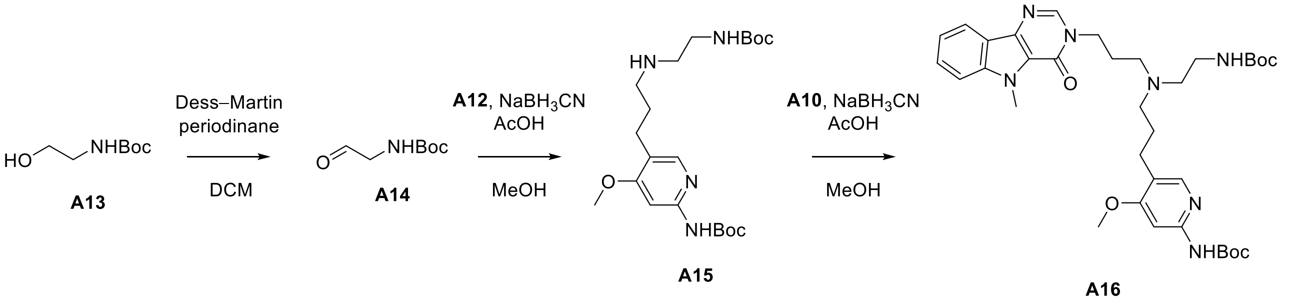

**Scheme 7:** Synthesis of intermediate **A15** and **A16**.

**Synthesis of tert-butyl (5-(3-((2-((tert-butoxycarbonyl)amino)ethyl)amino)propyl)-4-methoxypyridin-2-yl)carbamate) (A15)**: Dess-Martin periodinane (325 mg, 0.768 mmol) was added to a solution of **A13** (70.0 mg, 0.427 mmol) in DCM (5 mL), and the mixture was stirred for 1 h. The reaction mixture was quenched by addition of 10% (w/v) aqueous $Na_2S_2O_3$ (3 mL) and saturated $NaHCO_3$ solution (3 mL). The organic layer was extracted by DCM (10 mL × 2), dried over $Na_2SO_4$, and concentrated in vacuo. The crude aldehyde **A14** was used without further purification. AcOH (30.5 mL, 0.533 mmol, 1.5 equiv.) was added to a mixture of crude **A14** and amine **A12** (100 mg, 0.355 mmol) in MeOH, and the mixture was stirred for 30 min. After $NaBH_3CN$ (67.0 mg, 1.07 mmol, 3 equiv.) was added, the reaction mixture was stirred for 2 d. To the resulting mixture, saturated aqueous $NaHCO_3$ was added, and the organic layer was extracted with DCM, dried over $Na_2SO_4$, and concentrated in vacuo. The crude product was purified by column chromatography (Sfär KP-Amino D 11 g, 0-5% DCM/MeOH) to give the **A15** (55.1 mg, 37%). $^1$H NMR (400 MHz, $CDCl_3$) δ 8.59 (br s, 1H), 7.90 (s, 1H), 7.55 (s, 1H), 4.98 (br s, 1H), 3.90 (s, 3H), 3.22 (m, 2H), 2.71 (t, J = 7.3 Hz, 2H), 2.60 (t, J = 8.9 Hz, 2H), 2.54 (t, J = 8.9 Hz, 2H), 1.75-1.68 (m, 2H), 1.54 (s, 9H), 1.44 (s, 9H); $^{13}$C NMR (150 MHz, $CDCl_3$) δ 165.3, 156.1, 152.8, 152.6, 147.5, 121.0, 94.6, 80.7, 79.2, 55.3, 49.5, 49.1, 40.4, 30.0, 28.4, 28.4, 25.0; HR-MS (ESI): Calcd for $C_{21}H_{37}N_4O_5^+$ [M+H]$^+$; 425.2758; found, 425.2780.

**Synthesis of tert-butyl (5-(3-((2-((tert-butoxycarbonyl)amino)ethyl)(3-(5-methyl-4-oxo-4,5-dihydro-3H-pyrimido[5,4-b]indol-3-yl)propyl)amino)propyl)-4-methoxypyridin-2-yl)carbamate (A16)**: AcOH (1 uL, 17 mmol, 0.1 equiv.) was added to a mixture of the crude aldehyde **A10** and amine **A15** (85.4 mg, 0.201 mmol), and the mixture was stirred at room temperature for 30 min. Subsequently, $NaBH_3CN$ (102 mg, 1.61 mmol, 9 equiv.) was added, and the mixture was stirred at room temperature overnight, whereupon saturated aqueous $NaHCO_3$ (5 mL) was added, and the organic layer was extracted with DCM (10 mL x 3). The combined organic layer was dried over $Na_2SO_4$ and concentrated in vacuo. The crude product was purified by column chromatography (Sfär KP-Amino D 28 g, 30-75% hexane/AcOEt) to afford the amine **A16** (44.6 mg, 35%). $^1$H NMR (400 MHz, $CDCl_3$) δ 9.01 (s, 1H), 8.15 (d, J = 7.9 Hz, 1H), 8.01 (s, 1H),7.86 (s, 1H), 7.75 (bs, 1H), 7.58-7.53 (m, 1H), 7.51 (s, 1H), 7.46 (d, J = 7.9 Hz, 1H), 7.33-7.29 (m, 1H), 5.07 (br s, 1H), 4.26 (s, 3H), 4.15-4.10 (m, 1H), 3.89 (s, 3H), 3.19 (m, 2H), 2.55-2.47 (m ,6H), 1.99-1.94 (m, 2H), 1.73-1.63 (m, 4H), 1.53 (s, 9H), 1.43 (m, 9H); $^{13}$C NMR (150 MHz, $CDCl_3$) δ 165.3, 156.2, 155.5, 153.0, 152.9, 147.5, 143.3, 140.5, 138.2, 127.7, 121.9, 121.0, 121.0, 120.8, 110.1, 94.7, 80.7, 55.4, 53.7, 53.5, 53.3, 51.1, 45.0, 31.4, 28.5, 27.4, 27.0, 25.5; HR-MS (ESI): Calcd for $C_{35}H_{50}N_7O_6^+$ [M+H]$^+$; 664.3817; found, 664.3845.

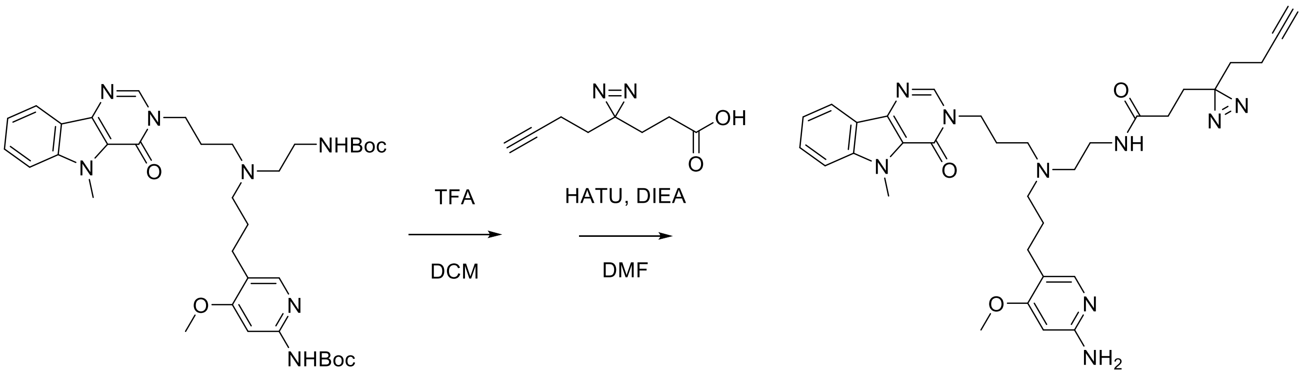

**A16** **3**

**Scheme 8:** Synthesis of **3**.

**Synthesis of N-(2-((3-(6-amino-4-methoxypyridin-3-yl)propyl)(3-(5-methyl-4-oxo-4,5-dihydro-3H-pyrimido[5,4-b]indol-3-yl)propyl)amino)ethyl)-3-(3-(but-3-yn-1-yl)-3H-diazirin-3-yl)propenamide (3)**: A mixture of Boc-amine **A16** (8.6 mg, 13.6 µmol) in DCM/TFA (v/v, 1 mL) was stirred for 2 h, and the mixture was concentrated in vacuo to give the crude product, which was used without further purification. Diazirine carboxylic acid (6.6 mg, 43 µmol) was pre-activated with HATU (18.6 mg, 49 µmol) and DIEA (15.7 mL, 98 µmol) in DMF (450 µL). To a solution of the crude amine and DIEA (6.5 mL, 41 µmol) in DMF (150 µL) the pre-activated solution (150 µL) was added, and the mixture was stirred for 4 h. An additional portion of pre-activated solution (50 µL) was added to the mixture, and the mixture was stirred overnight. The resulting mixture was diluted with MeOH, and directly purified by HPLC (SunFire Prep C18 OBD 5 um, 18 × 150 mm, 20-100% MeOH/$H_2O$ with 0.1% TFA for 60 min) to afford diazirine alkyne **3** (7.2 mg, 86%). $^1$H NMR (400 MHz, MeOD) δ 8.30 (s, 1H), 8.10 (d, J = 8.0 Hz, 1H), 7.63-7.57 (m, 3H), 7.34-7.30 (m, 1H), 6.35 (s, 1H), 4.26 (t, J = 6.6 Hz, 2H), 4.23 (s, 3H), 3.57 (br, 2H), 3.37-3.35 (m, 6H), 2.57 (t, J = 7.2 Hz, 2H), 2.31-2.27 (m, 2H), 2.25 (t, J = 2.64 Hz, 1H), 2.07 (t, J = 7.3 Hz, 2H), 2.00-1.92 (m, 4H), 1.73 (t, J = 7.5 Hz, 2H), 1.54 (t, J = 7.4 Hz, 2H); $^{13}$C NMR (150 MHz, MeOD) δ 176.4, 170.9, 157.3, 157.1, 145.3, 142.3, 139.3, 135.5, 129.5, 122.3, 121.8, 121.7, 118.6, 111.6, 93.0, 83.7, 70.5, 57.7, 54.7, 54.2, 52.0, 44.6, 36.1, 33.4,

32.9, 31.9, 30.8, 29.3, 25.6, 24.8, 24.3, 13.9; HR-MS (ESI): Calcd for $C_{33}H_{42}N_9O_3^+$ $[M+H]^+$; 612.3405; found, 612.3402.

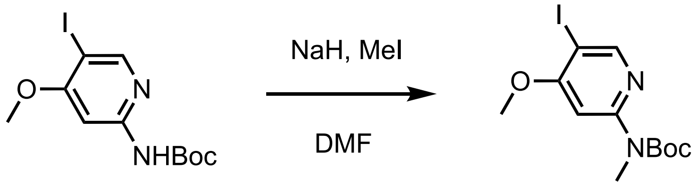

**A7** **A17**

**Scheme 9:** Synthesis of intermediate **A17**.

**Synthesis of *tert*-butyl (5-iodo-4-methoxypyridin-2-yl)(methyl)carbamate (A17):** To a solution of **A7** (992 mg, 2.83 mmol) in DMF (10.0 mL), NaH (125 mg, 60% in oil, 3.12 mmol) was added at 0 °C and the mixture was stirred at 0 °C for 10 minutes. To this mixture, MeI (212 μL, 3.40 mmol) was added at 0 °C. The mixture was stirred at 0 °C for 15 minutes, whereupon $H_2O$ (10 mL) and saturated aqueous $NaHCO_3$ (20 mL) were added subsequently, and the mixture was extracted with $Et_2O$ (50 mL x 3), dried over $MgSO_4$ and concentrated under vacuo. This crude was purified by column chromatography (Agela Technologies, Silica, 40 g, 0% - 40% ethylacetate in hexane) to afford *tert*-butyl (5-iodo-4-methoxypyridin-2-yl)(methyl)carbamate (**A17**) (938 mg, 91%). $^1$H NMR (400 MHz, $CDCl_3$) δ 8.46 (s, 1H), 7.41 (s, 1H), 3.94 (s, 3H), 3.38 (s, 3H), 1.54 (s, 9H) $^{13}$C NMR (150 MHz, MeOD) δ 165.1, 156.8, 154.3, 154.3, 102.6, 81.5, 78.7, 55.5, 33.7, 27.1 (3C) ; HR-MS (ESI): Calcd for $C_{12}H_{18}IN_2O_3^+$ $[M+H]^+$; 365.0357; found, 365.0383.

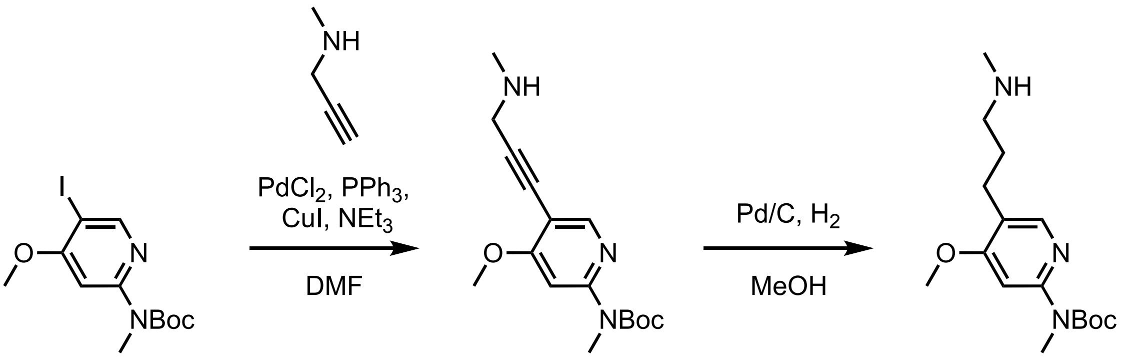

**A17** **A18** **A19**

**Scheme 10:** Synthesis of intermediate **A19**.

**Synthesis of *tert*-butyl (4-methoxy-5-(3-(methylamino)propyl)pyridin-2-yl)(methyl)carbamate (A19):** A mixture of **A17** (1.39 g, 3.83 mmol), $PdCl_2$ (46.0 mg, 0.153 mmol), $PPh_3$ (80.3 mg, 0.306 mmol), CuI (58.3 mg, 0.306 mmol), *N*-methylprop-2-yn-1-amine (793 mg, 11.5 mmol) in DMF (20 mL) was stirred at 50 °C for 1 h. After cooling to room temperature, saturated aqueous $NaHCO_3$ (50 mL) was added and the mixture was extracted with ethylacetate (50 mL x3) and concentrated under vacuo. This crude was purified by column chromatography (Agela Technologies, Silica, 20 g, 2% - 10% MeOH+5% aqueous $NH_4OH$ in DCM) to afford **A18** (1.06 g). A mixture of **A18** (437 mg, 1.43

mmol) and Pd/C (10%, 437 mg) in MeOH (15 mL) was stirred at room temperature under $H_2$ balloon (1 atm) for 1 h. The mixture was filtered through Celite pad and concentrated under vacuo. This crude material was purified by column chromatography (Biotage SNAP cartridge, KP-NH, 11g, 2% - 20% MeOH in DCM) to give *tert*-butyl (4-methoxy-5-(3-(methylamino)propyl)pyridin-2-yl)(methyl)carbamate (**A19**) (198 mg, 42% (2 steps)). $^1$H NMR (400 MHz, MeOD) δ 8.00 (s, 1H), 7.13 (s, 1H), 3.91 (s, 3H), 2.63 (t, $J$ = 7.4 Hz, 4H), 3.02 (s, 3H), 1.85-1.74 (m, 2H), 1.50 (s, 9H) $^{13}$C NMR (150 MHz, MeOD) δ 165.0, 155.5, 154.7, 146.9, 123.3, 102.6, 81.2, 54.7, 50.4, 34.2, 34.1, 28.2, 27.2, 24.4 (3C) ; HR-MS (ESI): Calcd for $C_{16}H_{28}N_3O_3^+$ [M+H]$^+$; 310.2125; found, 310.2113.

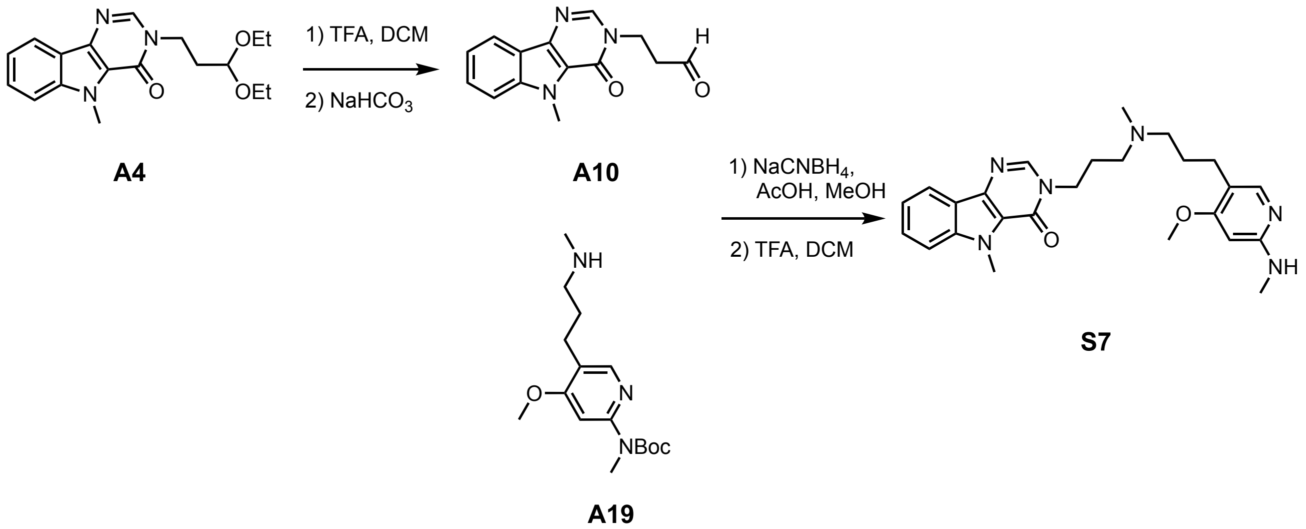

**Scheme 11:** Synthesis of **S7**.

**Synthesis of 3-(3-((3-(4-methoxy-6-(methylamino)pyridin-3-yl)propyl)(methyl)amino)propyl)-5-methyl-3,5-dihydro-4*H*-pyrimido[5,4-*b*]indol-4-one (S7)**: A mixture of **A4** (18.4 mg, 0.056 mmol) in TFA/DCM (1/1, 1.0 mL) was stirred at room temperature for 2 h. The mixture was concentrated under vacuo. To this mixture, saturated aqueous $NaHCO_3$ (1 mL) was added, and the mixture was extracted with DCM (1 mL x 3) and concentrated under vacuo to give **A10**. To this product, **A19** (25.8 mg, 0.050 mmol) in MeOH (500 µL) was added and the mixture was stirred at room temperature for 10 minutes. Subsequently $NaBH_3CN$ (6.98 mg, 0.111 mmol) and AcOH (6.35 µL, 0.111 mmol) were added and the mixture was stirred at room temperature overnight, whereupon saturated aqueous $NaHCO_3$ (1 mL) was added, and the mixture was extracted with DCM (2 mL x 3) and concentrated under vacuo. This crude was roughly purified by HPLC (20% - 80% MeOH+0.1%TFA in $H_2O$+0.1%TFA, 5 mL/min, 60 min) to afford products. The resultant material was dissolved into TFA/DCM (1/1, 2.0 mL) and the mixture was stirred at room temperature for 3 h, whereupon this mixture was concentrated under vacuo. The crude material was purified by HPLC (20% - 80% MeOH+0.1%TFA in $H_2O$+0.1%TFA, 5 ml/min, 60 min). The collected fractions were extracted with DCM/saturated aqueous $NaHCO_3$, and the resultant organic layer were concentrated to afford 3-(3-((3-(4-methoxy-6-(methylamino)pyridin-3-yl)propyl)(methyl)amino)propyl)-5-methyl-3,5-dihydro-4*H*-pyrimido[5,4-*b*]indol-4-one (**S7**) (4.9 mg, 20% (3 steps)). $^1$H NMR (400 MHz, $CDCl_3$) δ 8.16 (d, $J$ = 8.0 Hz, 1H), 8.06 (s, 1H), 7.73 (s, 1H), 7.58-7.52 (m, 1H), 7.46 (d, $J$ = 8.4 Hz, 1H), 7.34-7.28 (m, 1H), 5.82 (s, 1H), 4.48 (br s, 1H), 4.26 (s, 3H), 4.16 (t, $J$ = 6.9 Hz, 2H), 3.82 (s, 3H), 2.90 (d, $J$ = 4.9 Hz, 3H), 2.47 (t, $J$ = 7.4 Hz, 2H), 2.42-2.33 (m, 4H), 2.21 (s, 3H), 2.03-1.95 (m, 2H), 1.75-1.65 (m, 2H) $^{13}$C NMR (150 MHz, $CDCl_3$) δ 165.5, 160.1,

155.5, 147.5, 143.7, 140.4, 138.1, 127.5, 121.9, 120.9, 120.9, 120.6, 116.3, 109.9, 87.6, 57.2, 54.9, 54.1, 44.7, 41.6, 31.3, 29.4, 27.3, 26.8, 25.0 ; HR-MS (ESI): Calcd for $C_{25}H_{33}N_6O_2^+$ [M+H]$^+$; 449.2660; found, 449.2656.

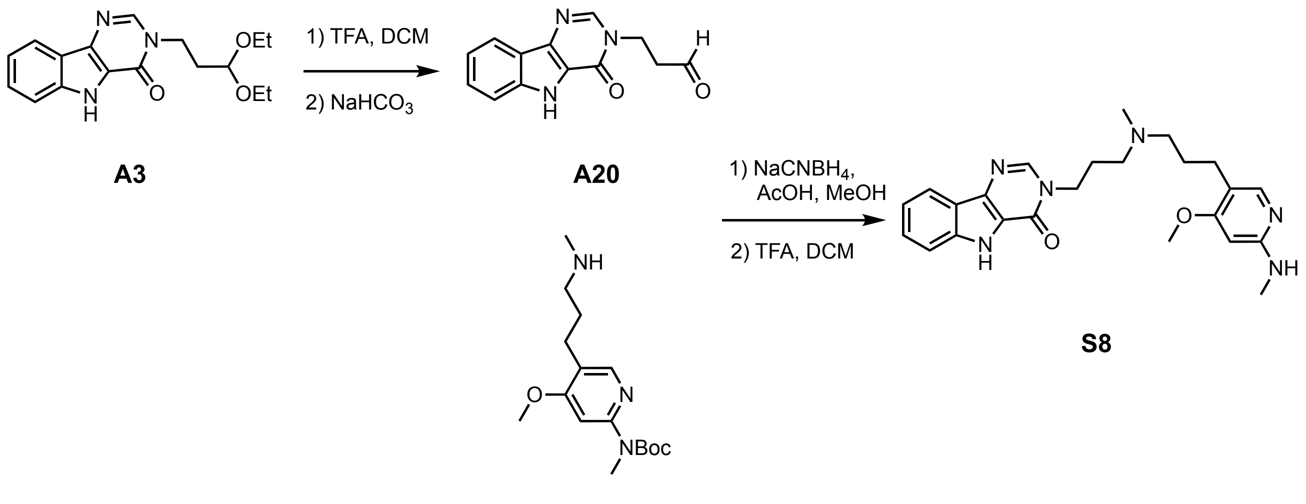

**Scheme 12:** Synthesis of **S8**.

**Synthesis of 3-(3-((3-(4-methoxy-6-(methylamino)pyridin-3-yl)propyl)(methyl)amino)propyl)-3,5-dihydro-4*H*-pyrimido[5,4-*b*]indol-4-one (S8):** A mixture of **A3** (452 mg, 1.43 mmol) in TFA/DCM (1/1, 14.3 mL) was stirred at room temperature for 30 minutes. The mixture was concentrated under vacuo. To this mixture, saturated aqueous $NaHCO_3$ (15 mL) was added, and the mixture was extracted with 30% MeOH in DCM (20 mL x 10) and concentrated under vacuo to give **A20**. To this product, **A19** (443 mg, 1.43 mmol) in MeOH (15.4 mL) was added and the mixture was stirred at room temperature for 2 minutes. Subsequently $NaBH_3CN$ (180 mg, 2.86 mmol) and AcOH (164 µL, 2.86 mmol) were added and the mixture was stirred at room temperature overnight, whereupon saturated aqueous $NaHCO_3$ (30 mL) was added and the mixture was extracted with DCM (20 mL x 3) and concentrated under vacuo. The resultant material was dissolved into TFA/DCM (1/1, 14.3 mL) and the mixture was stirred at room temperature for 5 h, whereupon this mixture was concentrated under vacuo. To this mixture, saturated aqueous $NaHCO_3$ (10 mL) was added and the mixture was extracted with DCM (30 mL x6) and concentrated. The resultant crude was purified by column chromatography (Biotage SNAP cartridge, KP-NH, 11g, 2 unit connected, 100% ethylacetate, 5CV, 30 ml/min, then 2% - 20% MeOH in DCM, 5CV, 30 ml/min) to afford 3-(3-((3-(4-methoxy-6-(methylamino)pyridin-3-yl)propyl)(methyl)amino)propyl)-3,5-dihydro-4*H*-pyrimido[5,4-*b*]indol-4-one (**S8**) (279 mg, 45% (3 steps)). $^1$H NMR (400 MHz, $CDCl_3$) δ 11.2 (s, 1H), 8.15 (d, *J* = 8.0 Hz, 1H), 8.13 (s, 1H), 7.75 (s, 1H), 7.60 (d, *J* = 8.4 Hz, 1H), 7.52-7.46 (m, 1H), 7.31-7.28 (m, 1H), 5.82 (s, 1H), 5.05 (br s, 1H), 4.27 (t, *J* = 6.8 Hz, 2H), 3.81 (s, 3H), 2.90 (d, *J* = 5.0 Hz, 3H), 2.49 (t, *J* = 7.4 Hz, 2H), 2.45-2.35 (m, 4H), 2.23 (s, 3H), 2.11-2.01 (m, 2H), 1.80-1.66 (m, 2H) $^{13}$C NMR (150 MHz, $CDCl_3$) δ 165.6, 160.2, 155.3, 147.1, 143.5, 139.5, 139.0, 127.7, 122.3, 121.6, 120.8, 120.7, 116.2, 112.7, 87.5, 57.2, 54.9, 54.1, 45.0, 41.7, 29.4, 27.3, 26.9, 25.1 ; HR-MS (ESI): Calcd for $C_{24}H_{29}N_6O_2^-$ [M-H]$^-$; 433.2358; found, 433.2353.

**Compound characterization.**

**NMR Spectra of Compound A3**

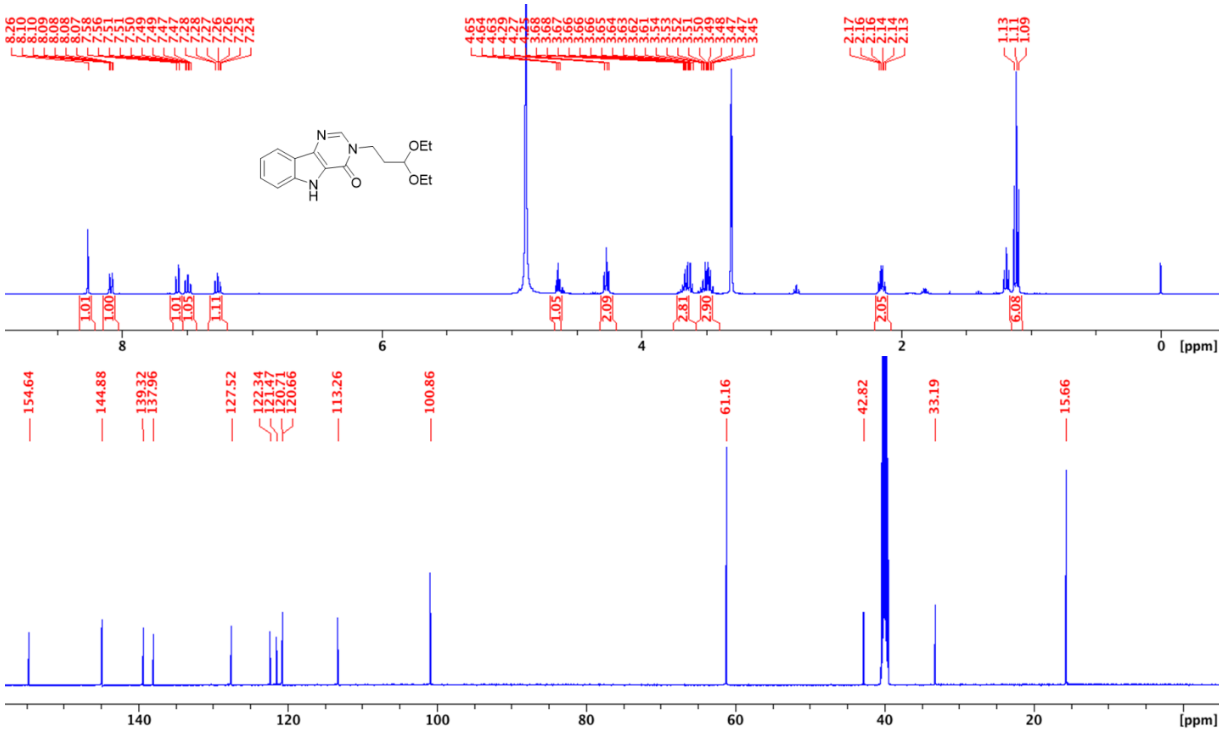

**NMR Spectra of Compound A4**

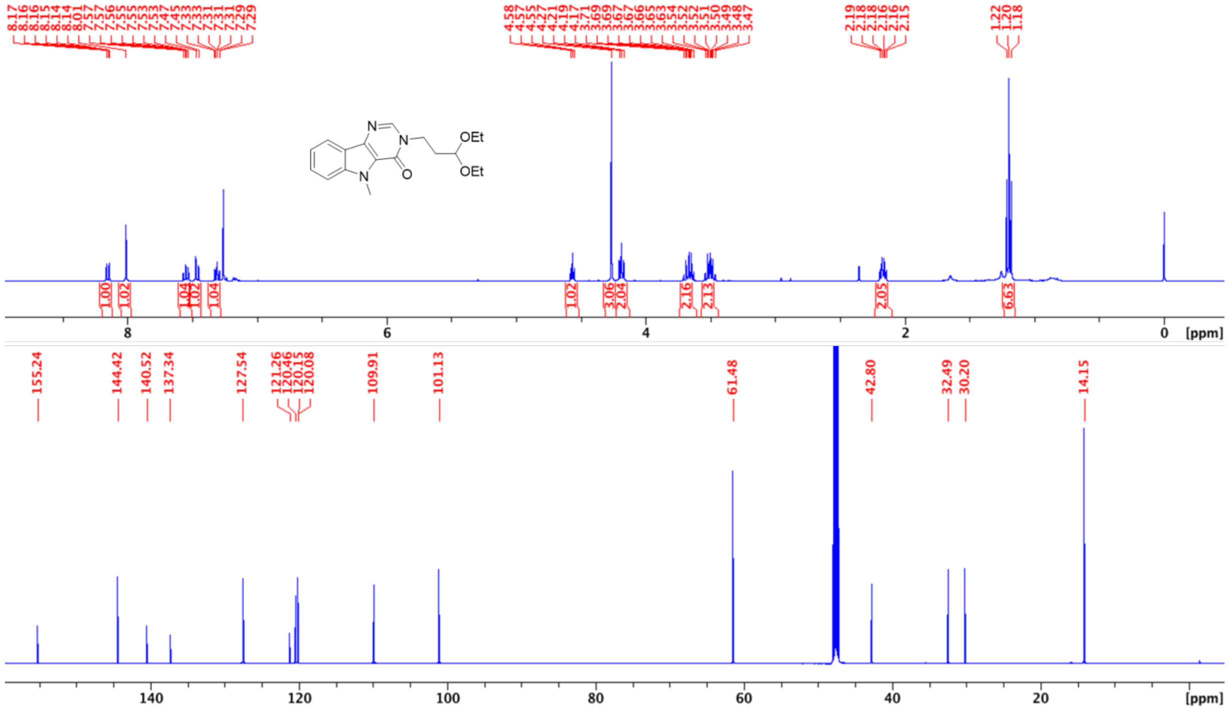

### NMR Spectra of Compound A7

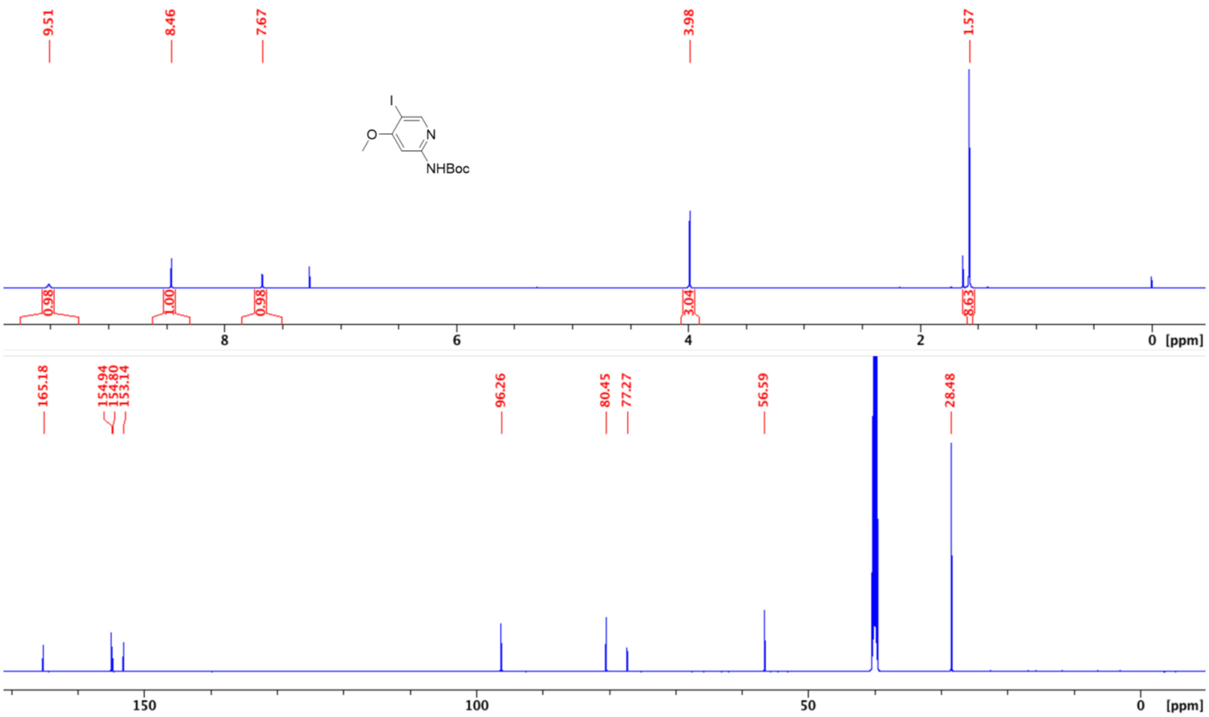

### NMR Spectra of Compound A9

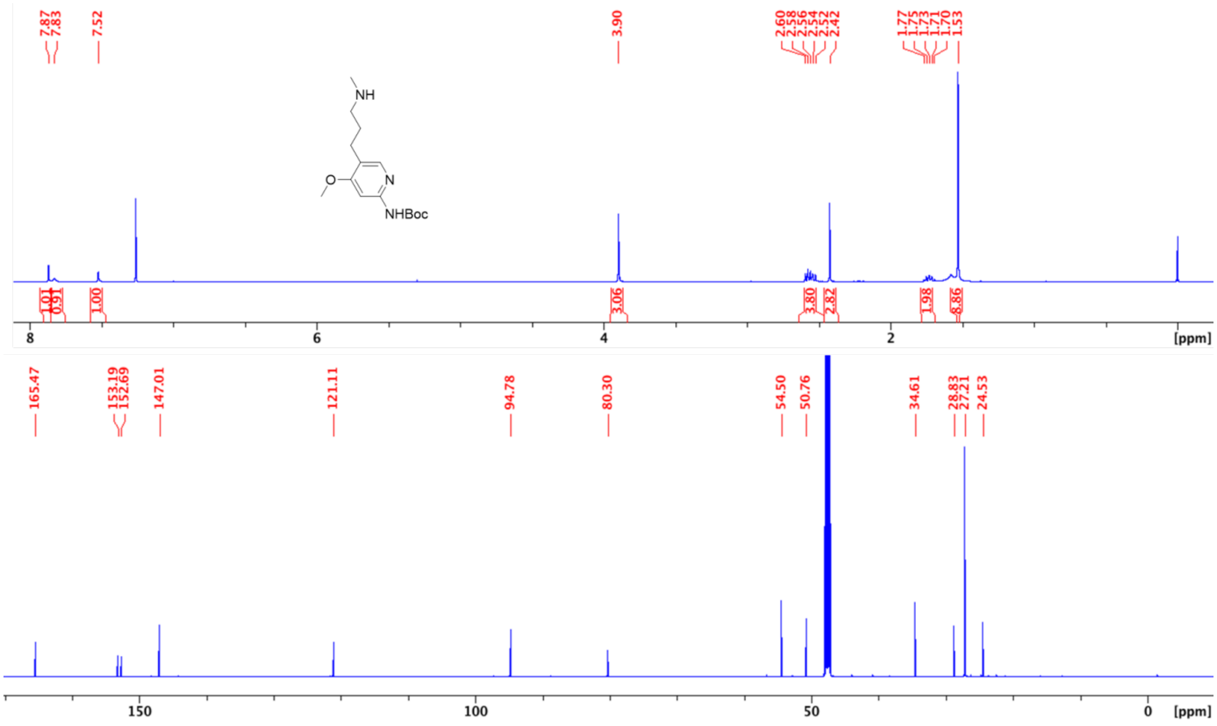

**NMR Spectra of Compound 2**

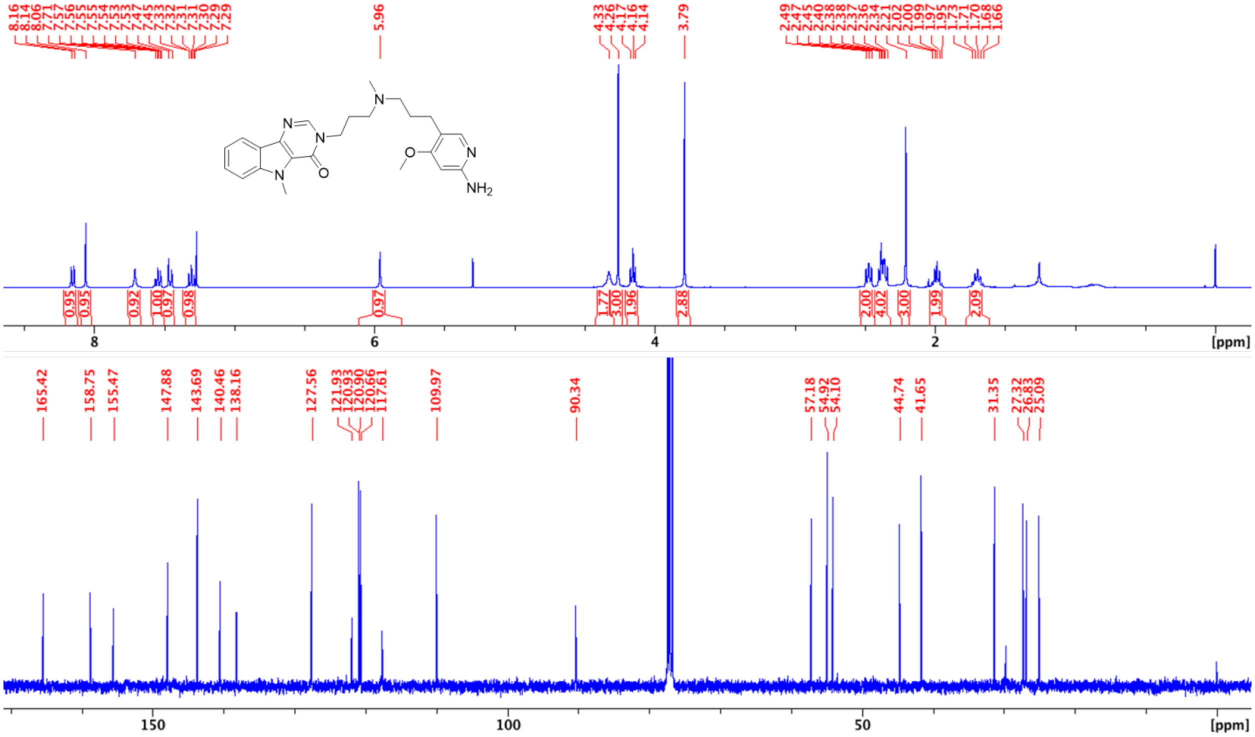

**Analytical HPLC chart of Compound 2**

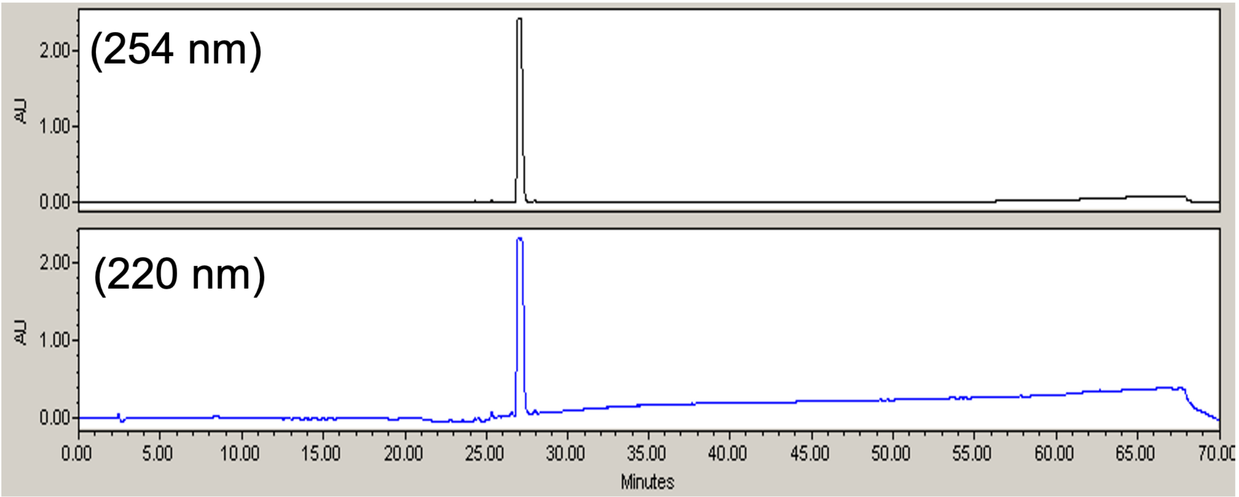

### NMR Spectra of Compound A12

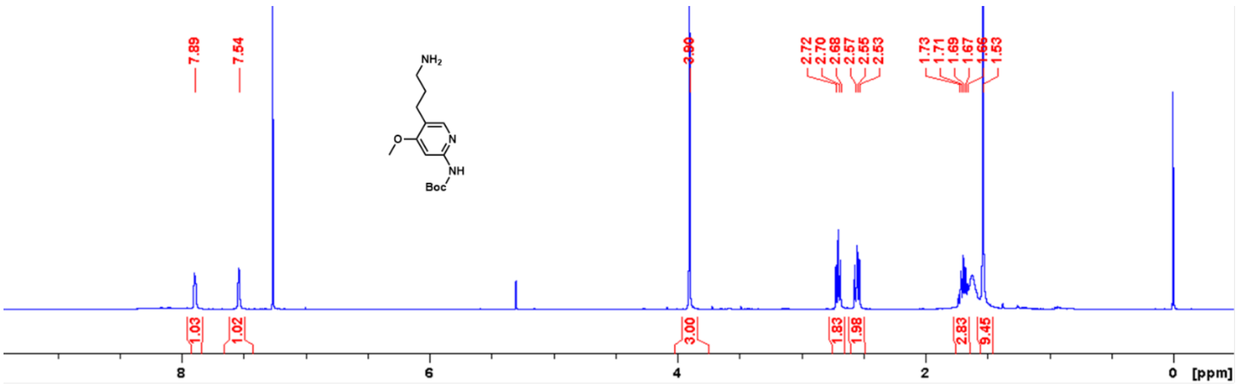

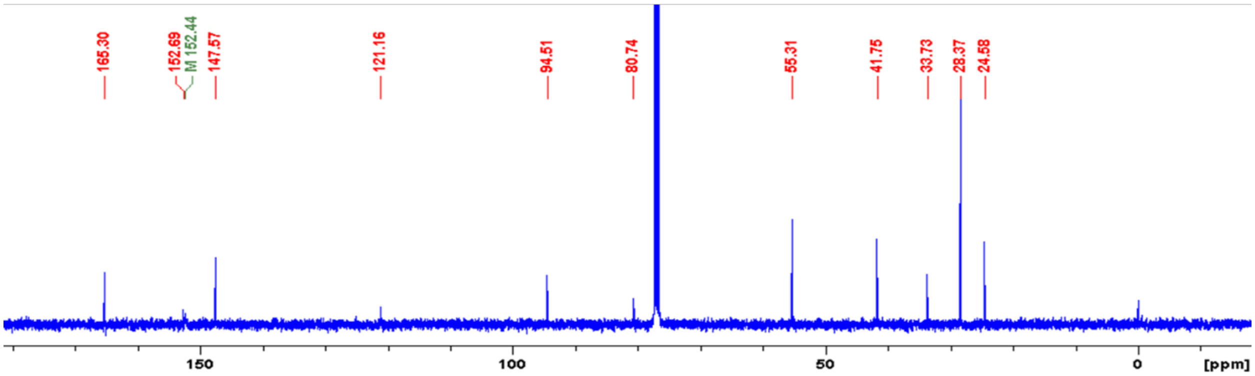

### NMR Spectra of Compound A15

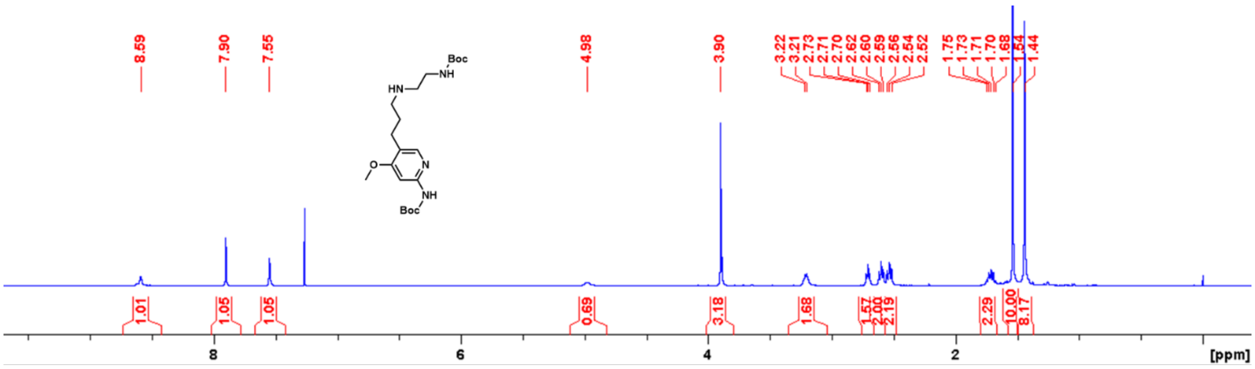

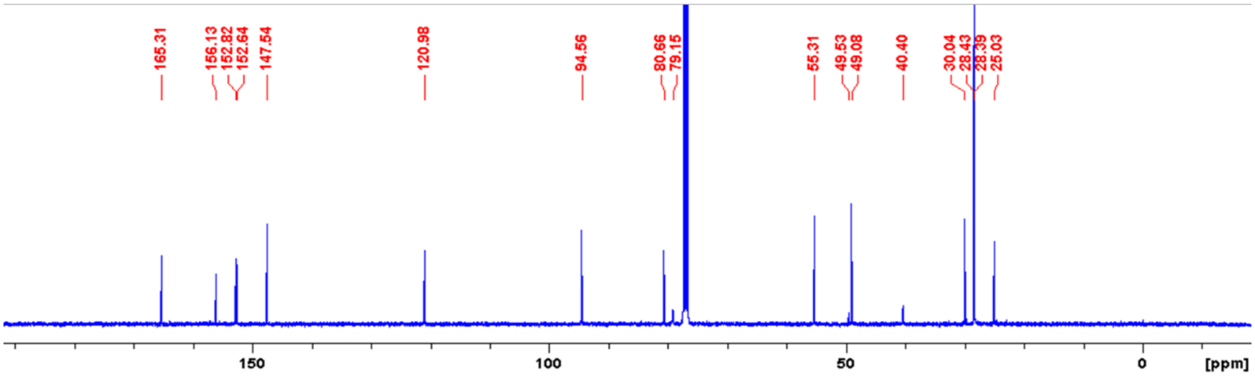

### NMR Spectra of Compound A16

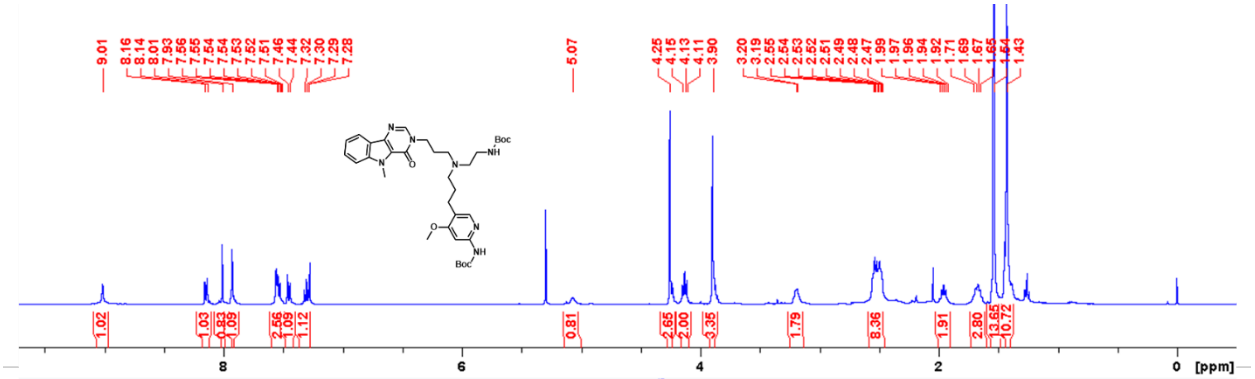

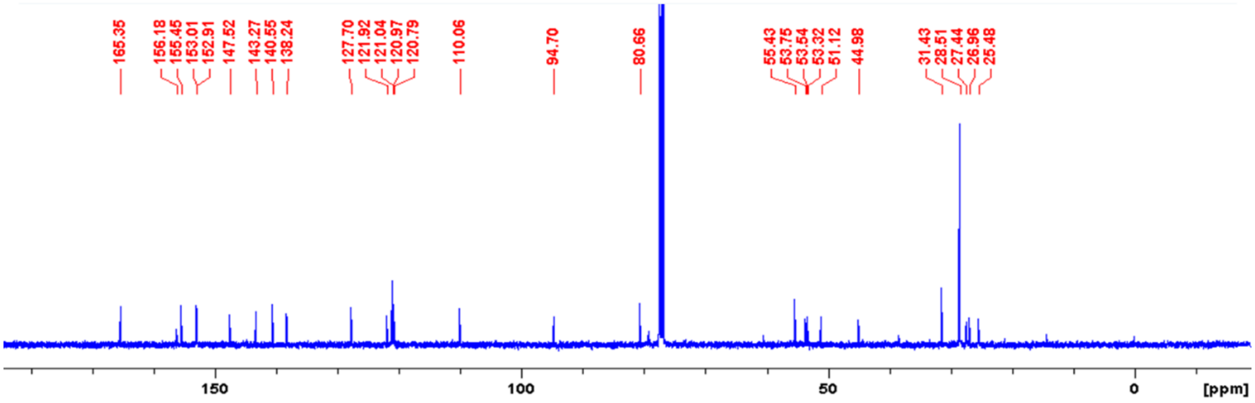

### NMR Spectra of Compound 3

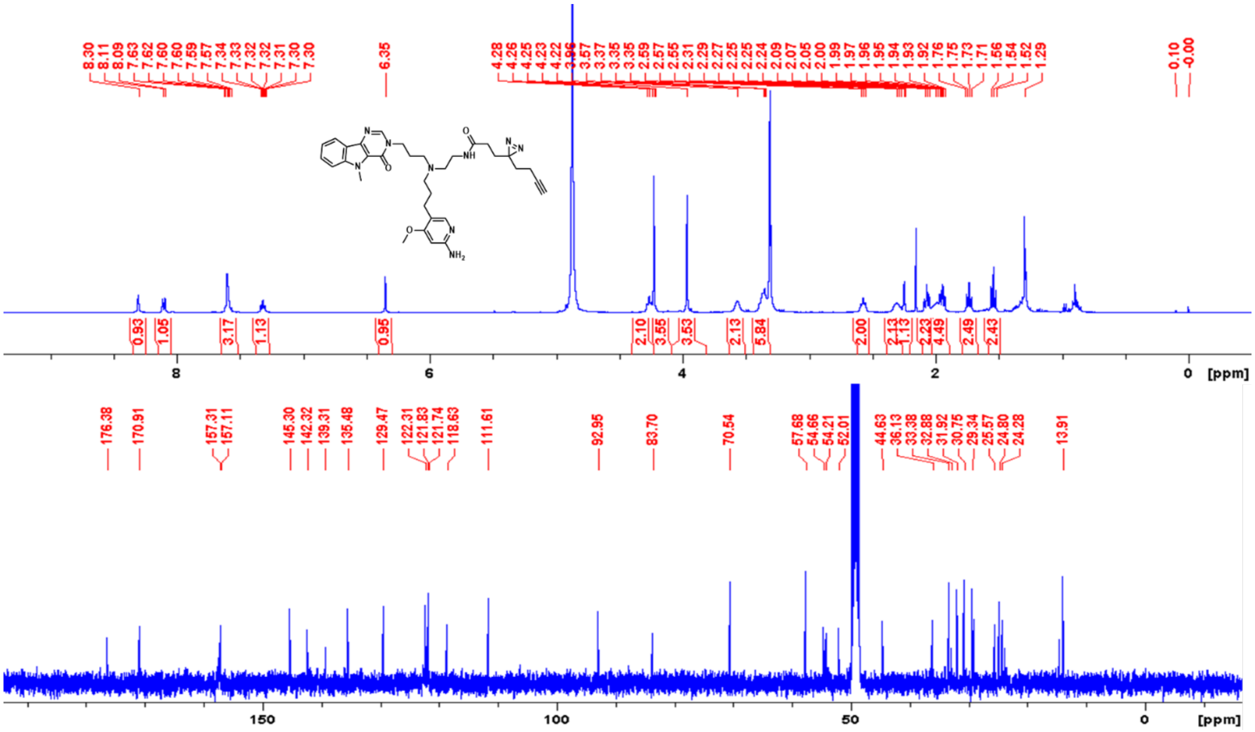

### NMR Spectra of Compound A17

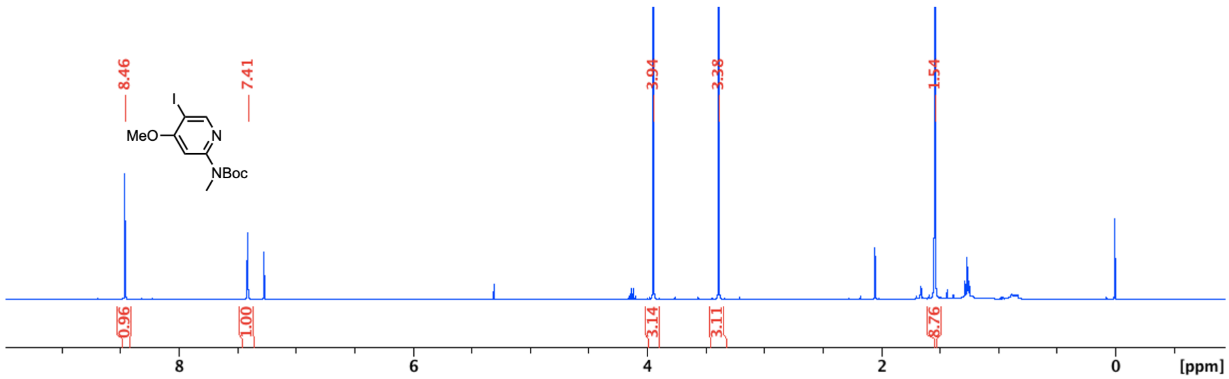

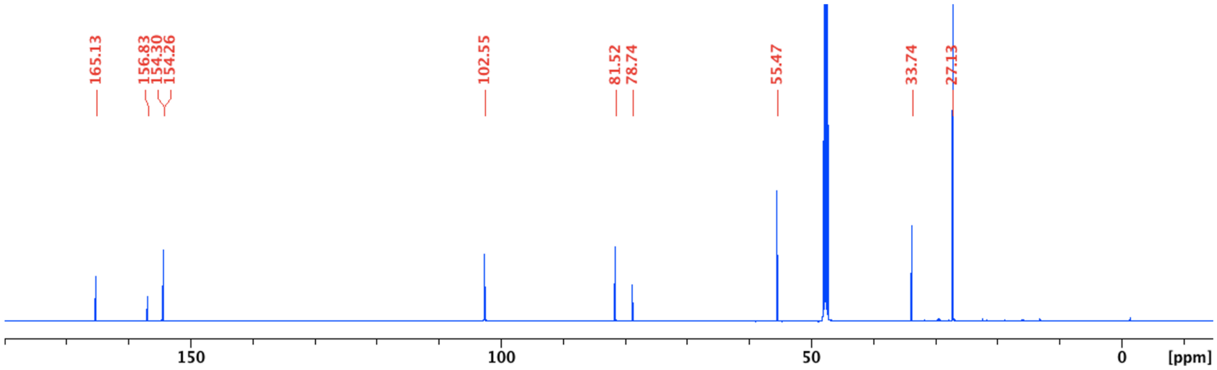

### NMR Spectra of Compound A19

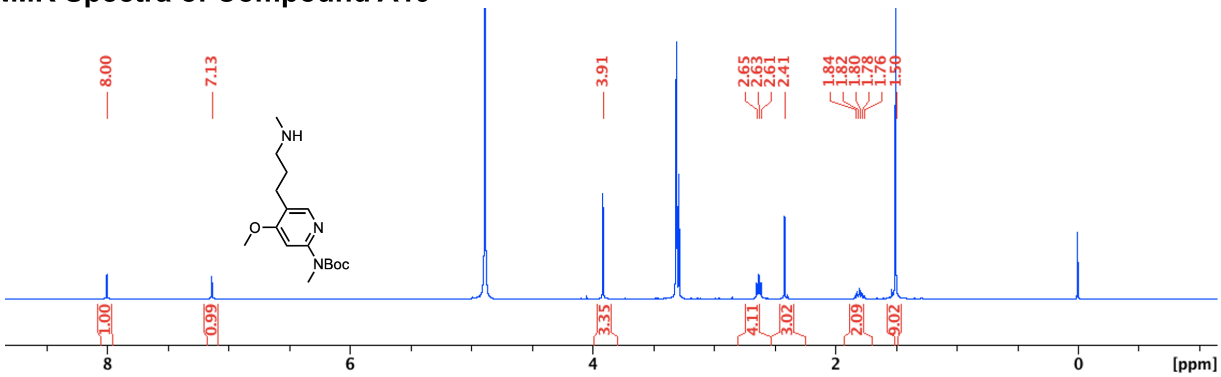

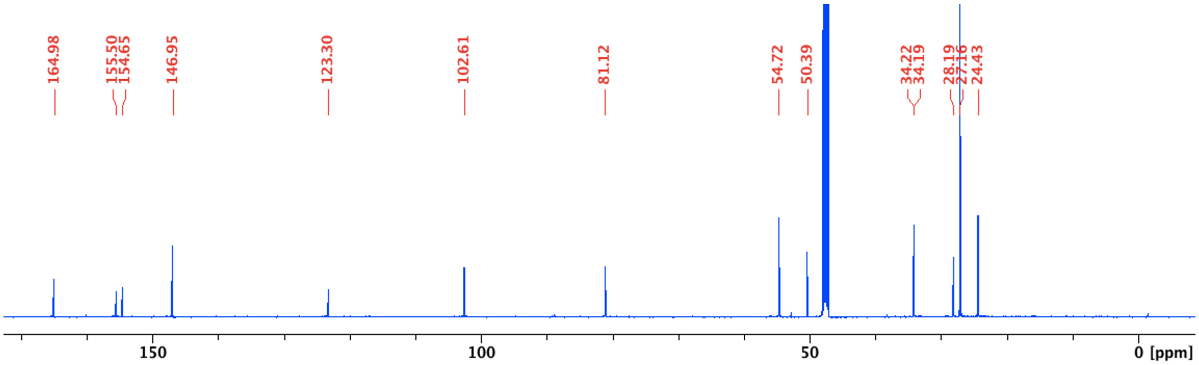

### NMR Spectra of Compound S7

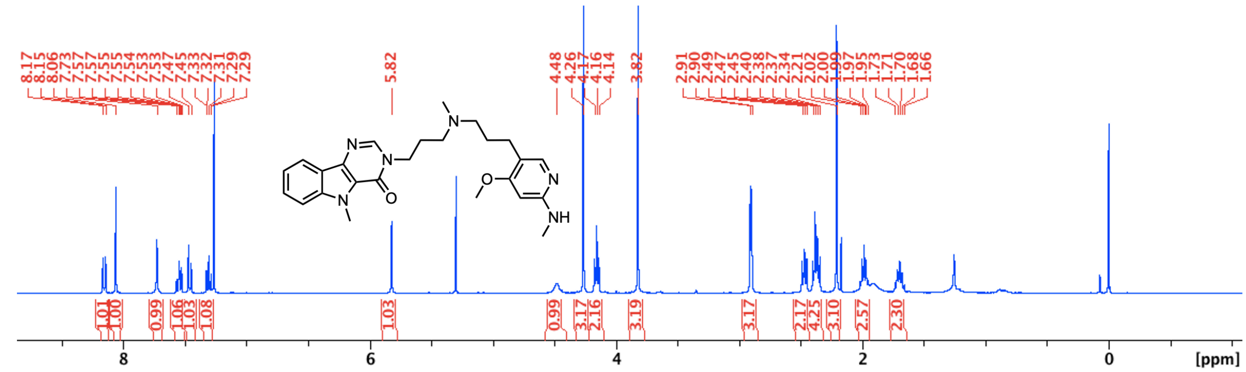

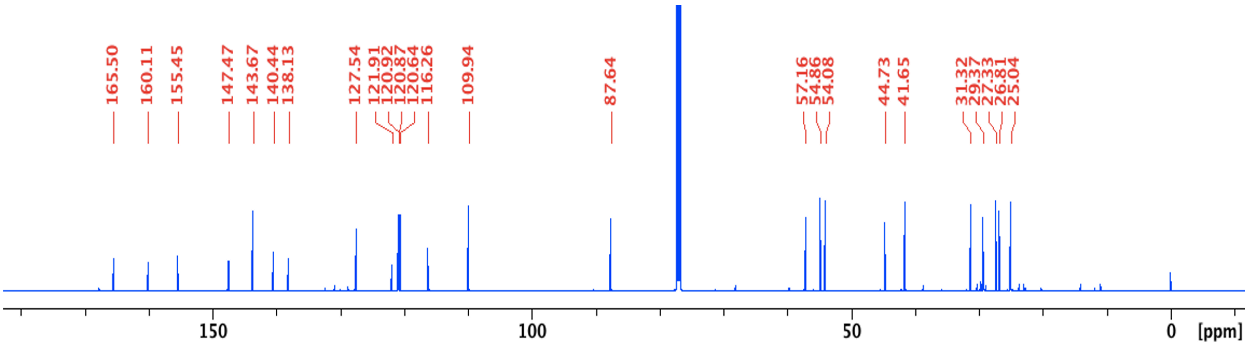

### NMR Spectra of Compound S8

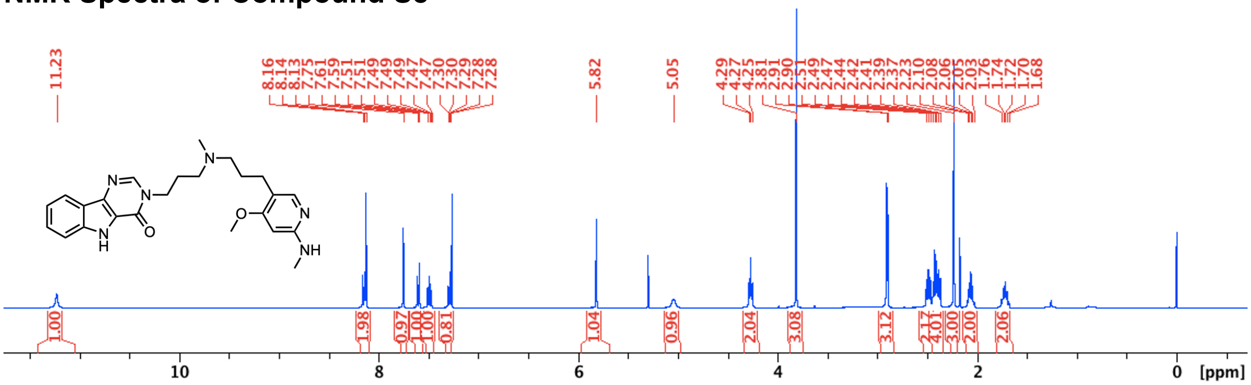

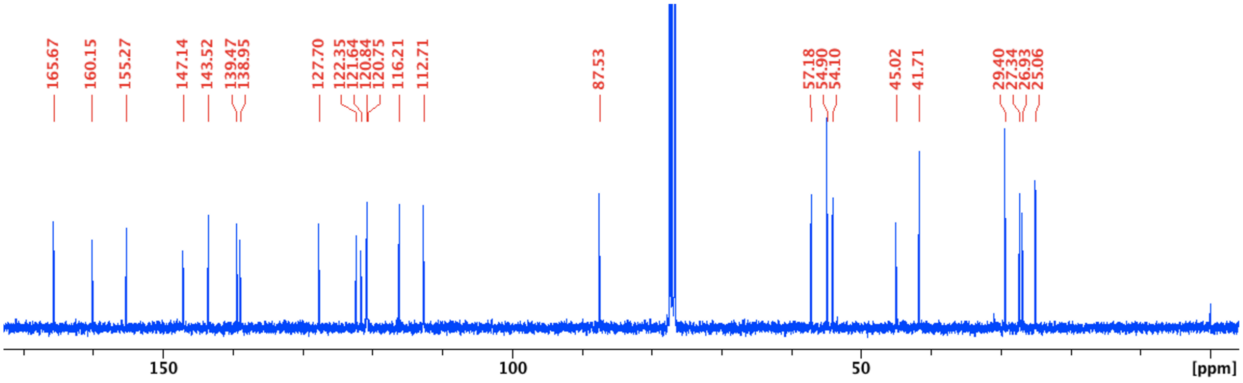

### NMR Spectra of Compound F2

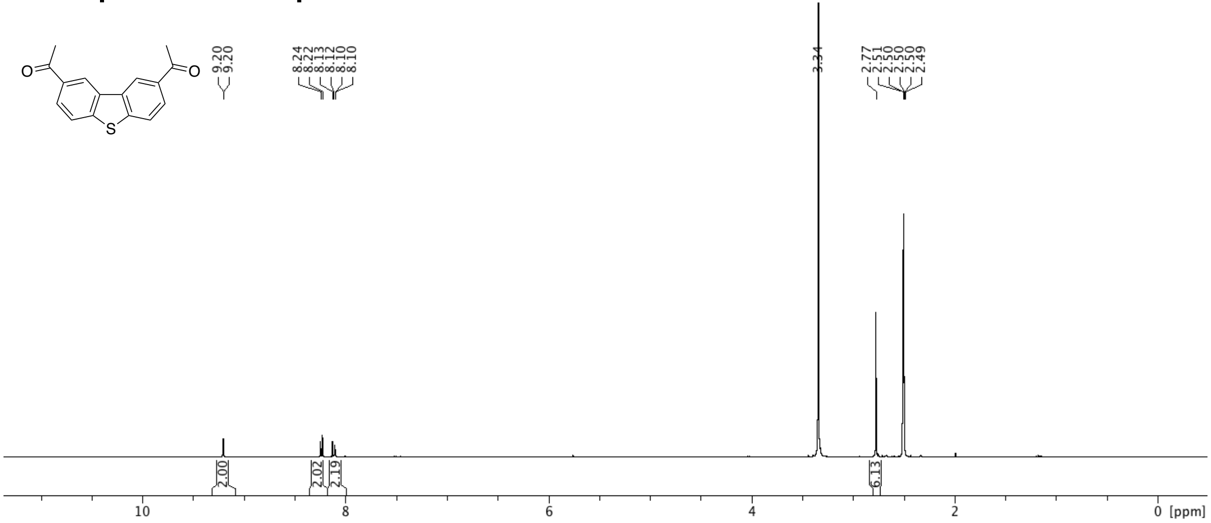

### NMR Spectra of Compound F3

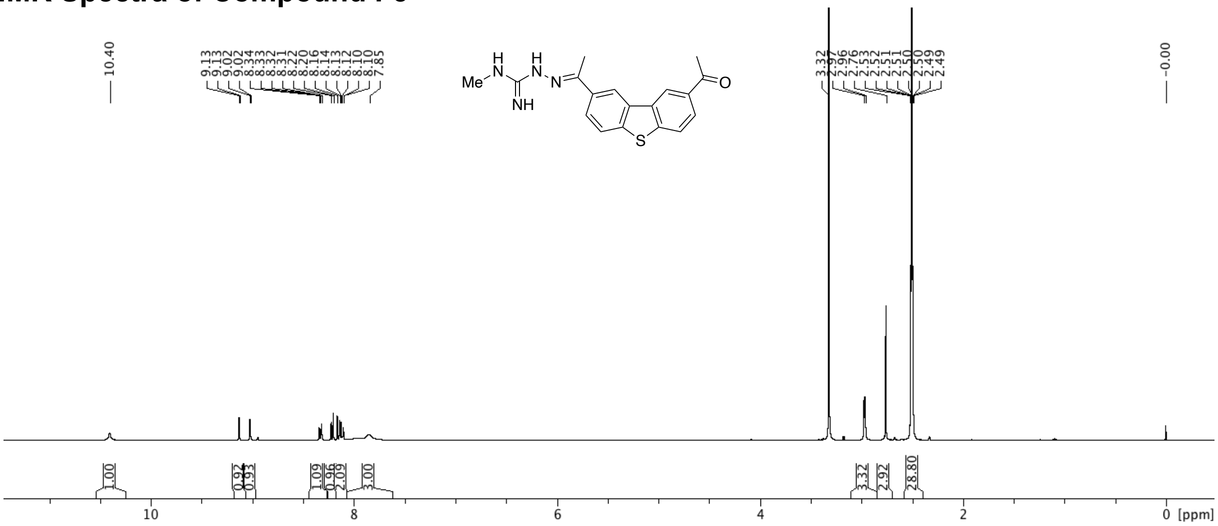

### NMR Spectra of Compound S1

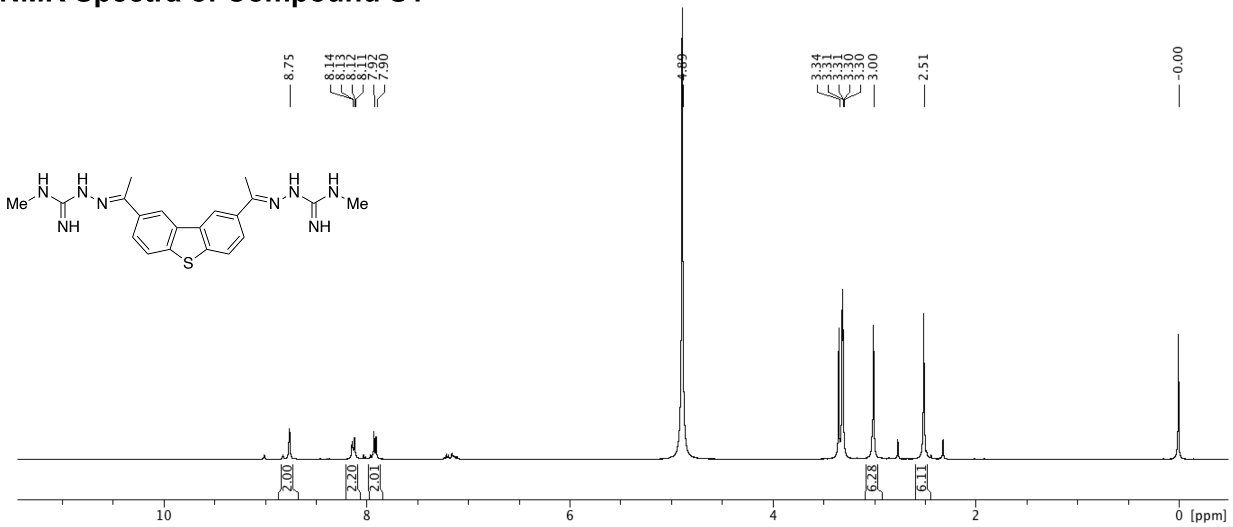

### NMR Spectra of Compound F4

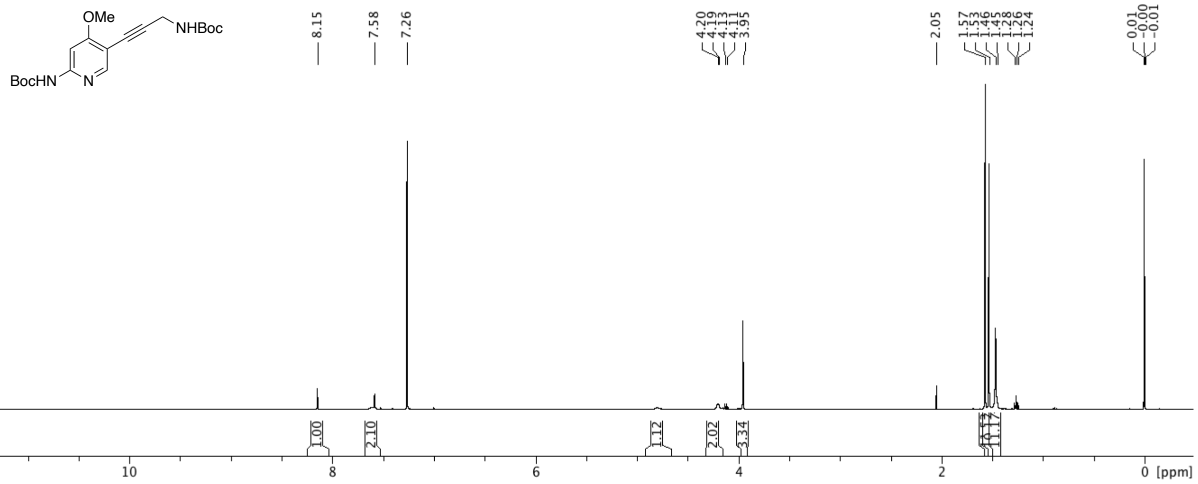

### NMR Spectra of Compound F5

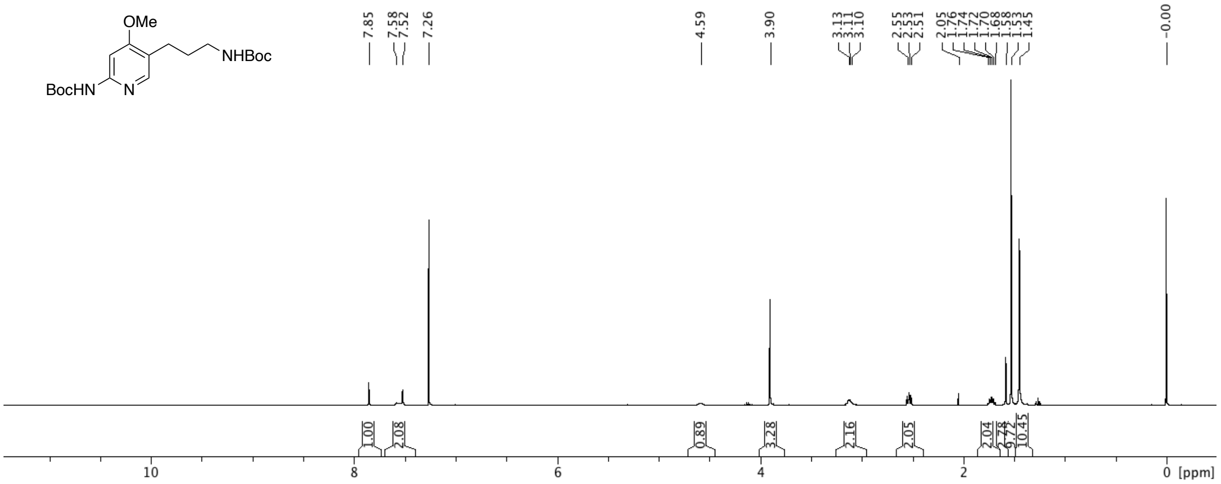

### NMR Spectra of Compound F6

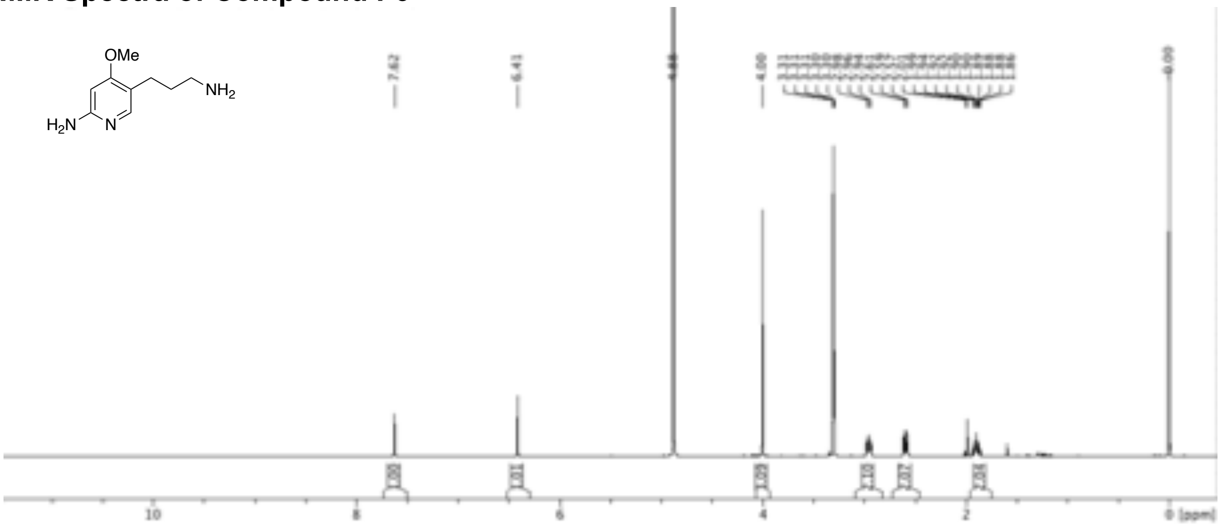

### NMR Spectra of Compound F7

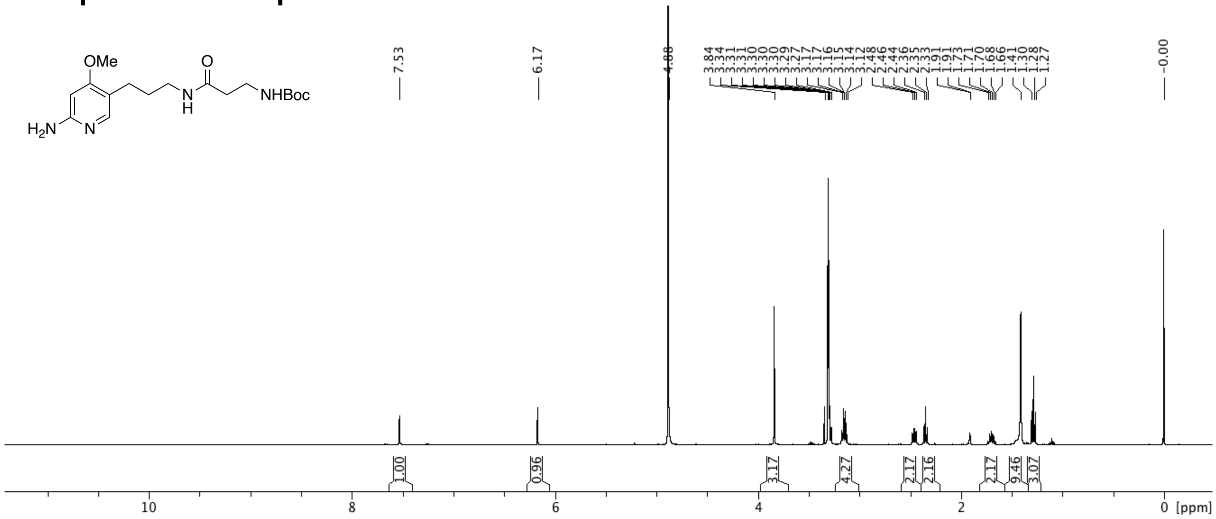

### NMR Spectra of Compound F8

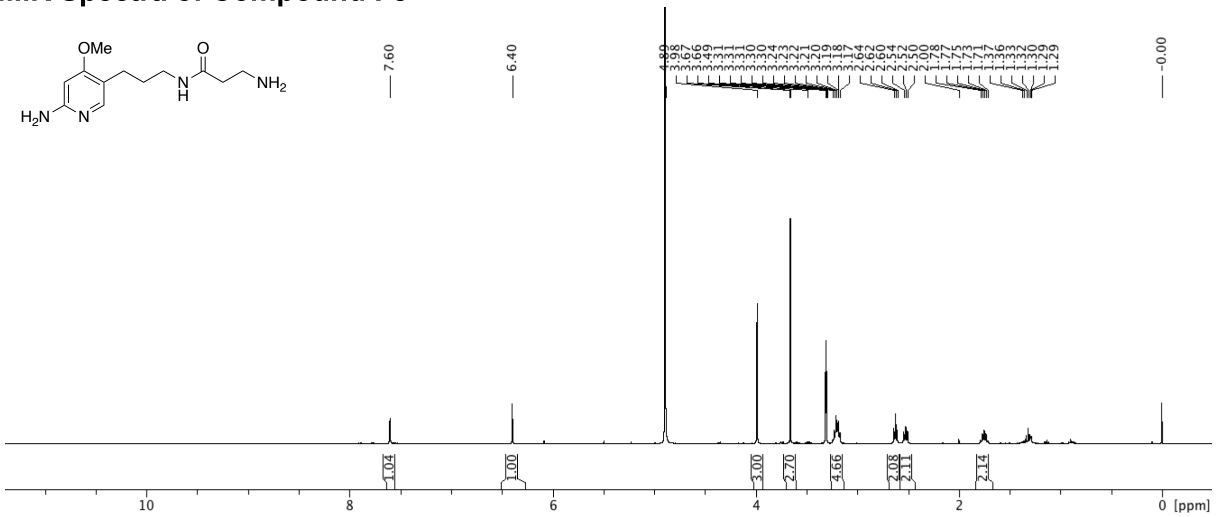

### NMR Spectra of Compound F9

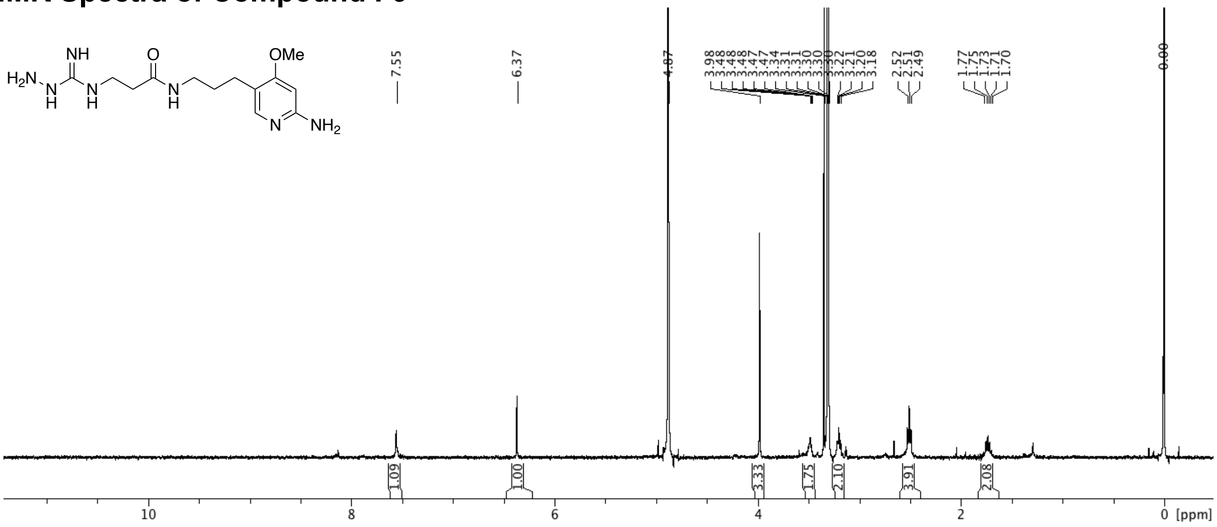

### NMR Spectra of Compound F10

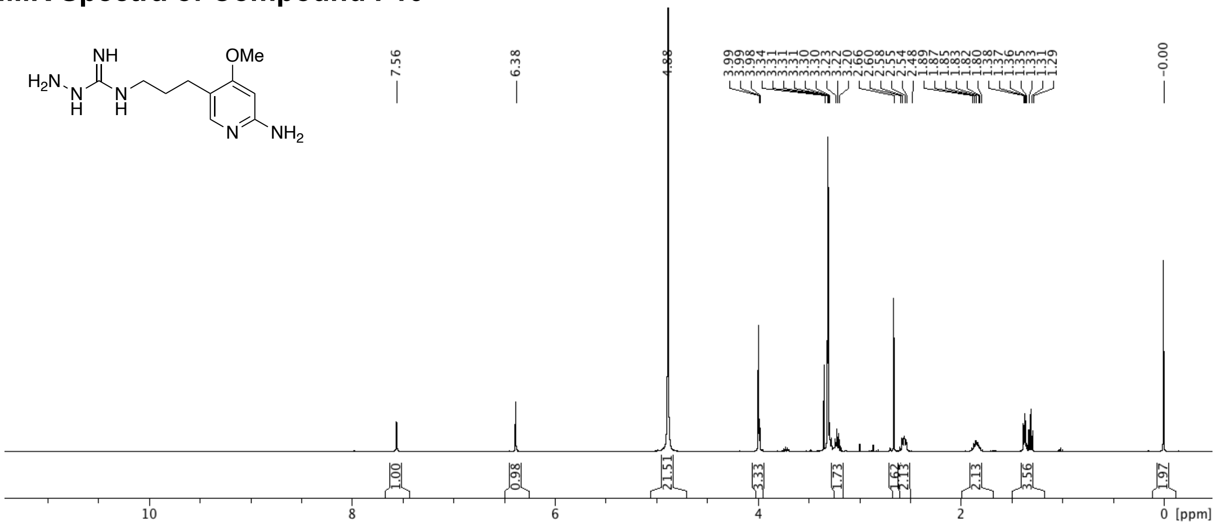

### NMR Spectra of Compound F12

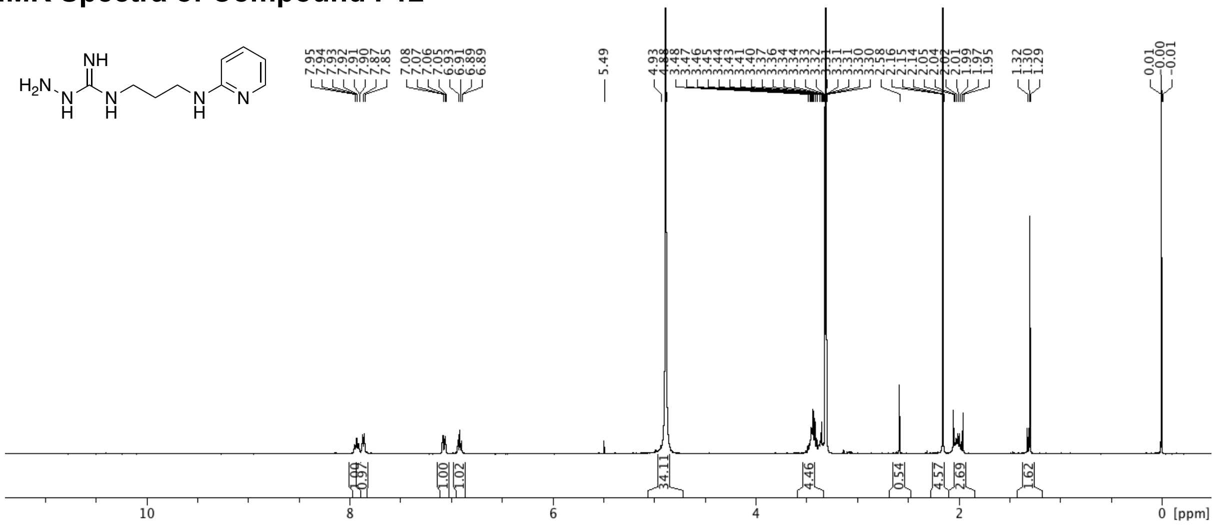

### NMR Spectra of Compound S2

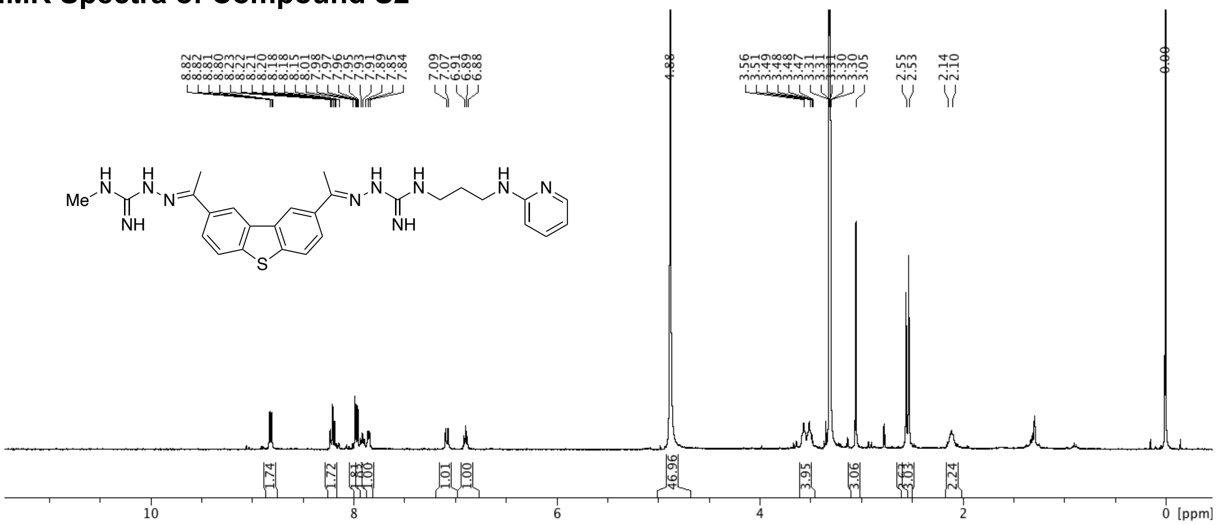

### NMR Spectra of Compound S3

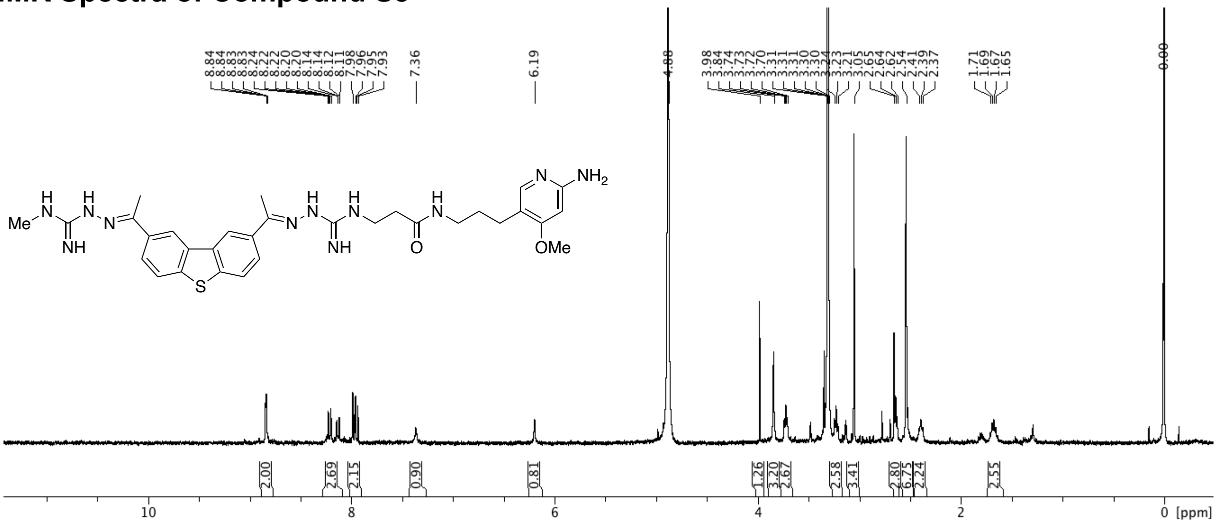

### NMR Spectra of Compound S4

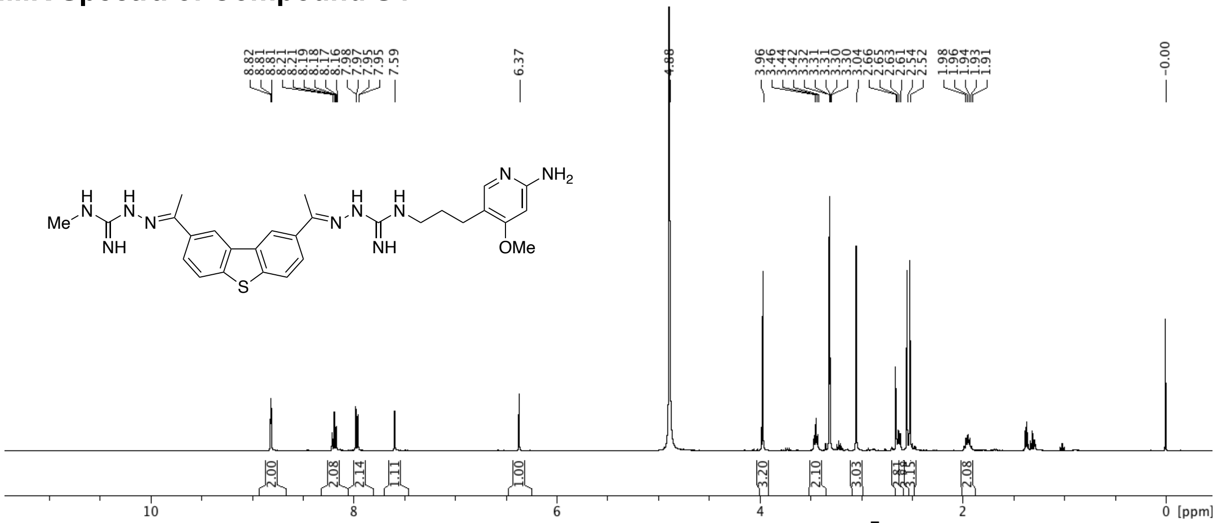

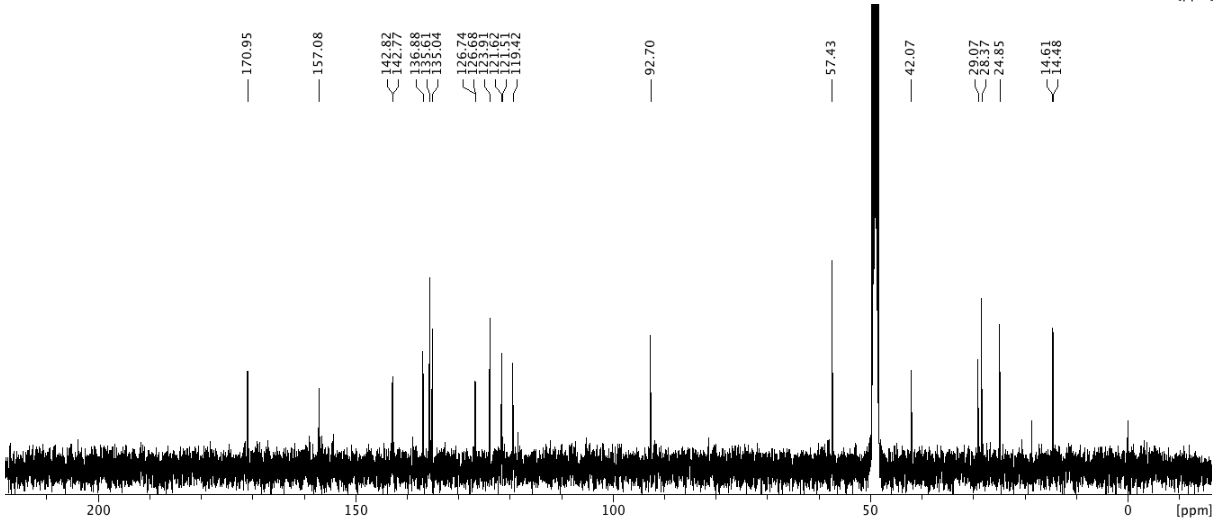

### NMR Spectra of Compound S5

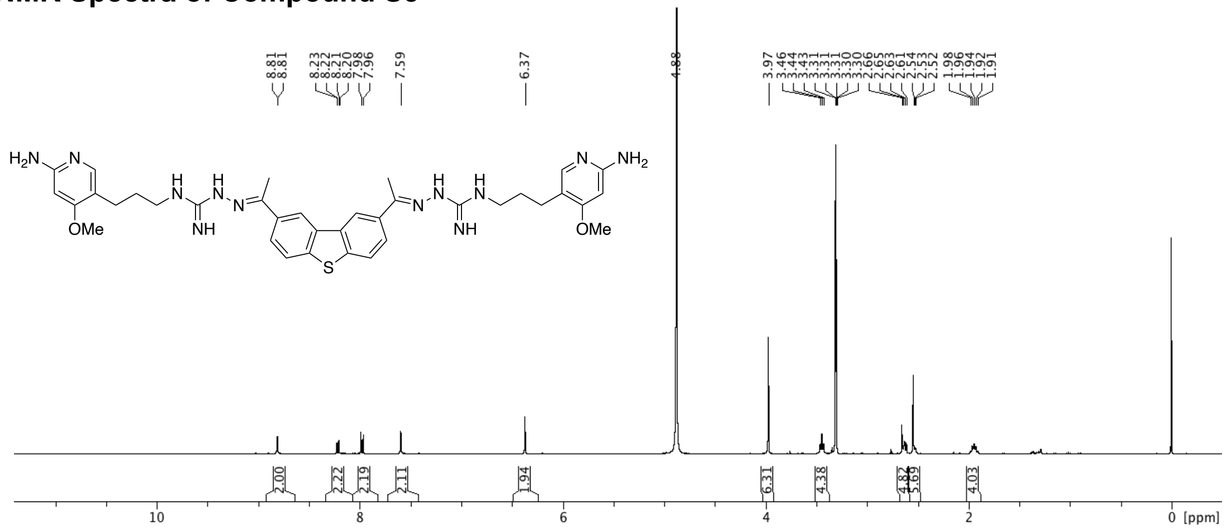

### NMR Spectra of Compound S6

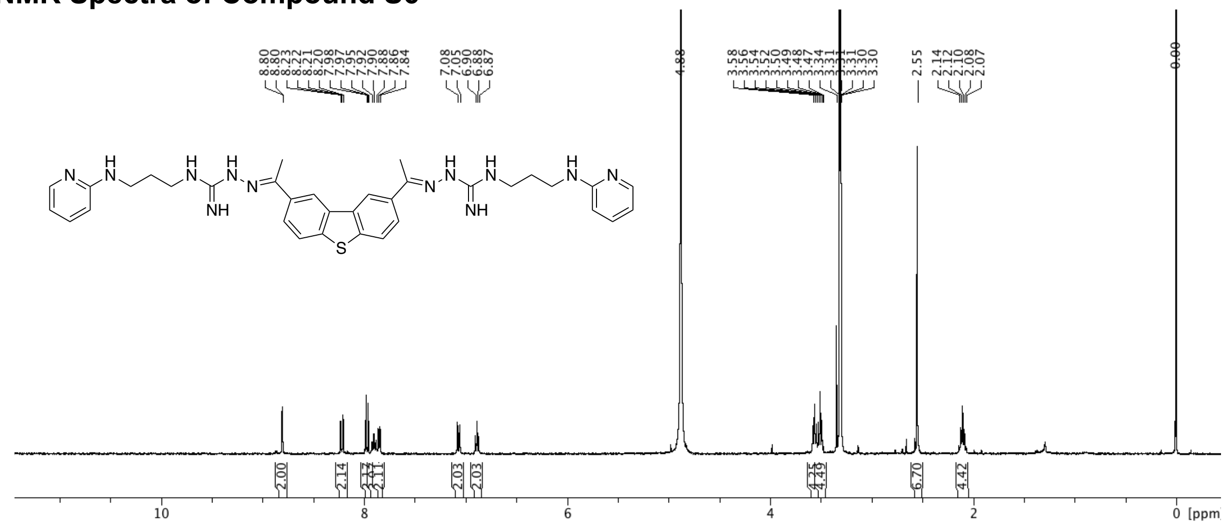

(11) Ryckaert, J. P.; Ciccotti, G.; Berendsen, H. J. C. Numerical-Integration of Cartesian

Equations of Motion of a System with Constraints - Molecular-Dynamics of N-Alkanes. *J Comput Sci* **1977**, *23* (3), 327-341.
(12) Essmann, U.; Perera, L.; Berkowitz, M. L.; Darden, T.; Lee, H.; Pedersen, L. G. A smooth particle mesh Ewald method. *J Chem Phys* **1995**, *103* (19), 8577-8593.
(13) Buneman, O. Computer Simulation Using Particles (R. W. Hockney and J. W. Eastwood). *SIAM Review* **1983**, *25* (3), 425-426.
(14) Bayly, C. I.; Cieplak, P.; Cornell, W.; Kollman, P. A. A well-behaved electrostatic potential based method using charge restraints for deriving atomic charges: the RESP model. *J Phys Chem* **1993**, *97* (40), 10269-10280.
(15) Wang, J.; Wolf, R. M.; Caldwell, J. W.; Kollman, P. A.; Case, D. A. Development and testing of a general amber force field. *J Comput Chem* **2004**, *25* (9), 1157-1174.
(16) Wang, J.; Wang, W.; Kollman, P. A.; Case, D. A. Automatic atom type and bond type perception in molecular mechanical calculations. *J Mol Graph Model* **2006**, *25* (2), 247-260.
(17) Frisch, M.; Trucks, G.; Schlegel, H.; Scuseria, G.; Robb, M.; Cheeseman, J.; Scalmani, G.; Barone, V.; Mennucci, B.; Petersson, G. Gaussian 09, Revision D. 01, Gaussian, Inc., Wallingford CT. See also: URL: http://www. gaussian. com **2009**.
(18) Chen, J. L.; Zhang, P.; Abe, M.; Aikawa, H.; Zhang, L.; Frank, A. J.; Zembryski, T.; Hubbs, C.; Park, H.; Withka, J.; et al. Design, Optimization, and Study of Small Molecules That Target Tau Pre-mRNA and Affect Splicing. *J Am Chem Soc* **2020**, *142* (19), 8706-8727.
(19) Trott, O.; Olson, A. J. AutoDock Vina: improving the speed and accuracy of docking with a new scoring function, efficient optimization, and multithreading. *J Comput Chem* **2010**, *31* (2), 455-461.
(20) O'Boyle, N. M.; Banck, M.; James, C. A.; Morley, C.; Vandermeersch, T.; Hutchison, G. R. Open Babel: An open chemical toolbox. *J Cheminform* **2011**, *3*, 33.
(21) Gasteiger, J.; Marsili, M. Iterative partial equalization of orbital electronegativity—a rapid access to atomic charges. *Tetrahedron* **1980**, *36* (22), 3219-3228.
(22) Santos-Martins, D.; Solis-Vasquez, L.; Tillack, A. F.; Sanner, M. F.; Koch, A.; Forli, S. Accelerating AutoDock4 with GPUs and Gradient-Based Local Search. *J Chem Theory Comput* **2021**, *17* (2), 1060-1073.

(23) Solis, F. J.; Wets, R. J. B. Minimization by Random Search Techniques. *Math Oper Res* **1981**, *6* (1), 19-30.
(24) Ribeiro, J.; Rios-Vera, C.; Melo, F.; Schuller, A. Calculation of accurate interatomic contact surface areas for the quantitative analysis of non-bonded molecular interactions. *Bioinformatics* **2019**, *35* (18), 3499-3501.
(25) Szulc, N. A.; Mackiewicz, Z.; Bujnicki, J. M.; Stefaniak, F. fingeRNAt-A novel tool for high-throughput analysis of nucleic acid-ligand interactions. *PLoS Comput Biol* **2022**, *18* (6), e1009783.
(26) Kovalenko, A.; Hirata, F. Three-dimensional density profiles of water in contact with a solute of arbitrary shape: a RISM approach. *Chem Phys Lett* **1998**, *290* (1), 237-244.
(27) Luchko, T.; Gusarov, S.; Roe, D. R.; Simmerling, C.; Case, D. A.; Tuszynski, J.; Kovalenko, A., Three-Dimensional Molecular Theory of Solvation Coupled with Molecular Dynamics in Amber. *J Chem Theory Comput* **2010**, *6* (3), 607-624.
(28) Johnson, J.; Case, D.; Yamazaki, T.; Gusarov, S.; Kovalenko, A.; Luchko, T. Small molecule hydration energy and entropy from 3D-RISM. *J Condens Matter Phys* **2016**, *28* (34), 344002.
(29) Gusarov, S.; Ziegler, T.; Kovalenko, A. Self-consistent combination of the three-dimensional RISM theory of molecular solvation with analytical gradients and the Amsterdam density functional package. *J Phys Chem A* **2006**, *110* (18), 6083-6090.
(30) Velagapudi, S. P.; Luo, Y.; Tran, T.; Haniff, H. S.; Nakai, Y.; Fallahi, M.; Martinez, G. J.; Childs-Disney, J. L.; Disney, M. D. Defining RNA-Small Molecule Affinity Landscapes Enables Design of a Small Molecule Inhibitor of an Oncogenic Noncoding RNA. *ACS Cent Sci* **2017**, *3* (3), 205-216.
(31) McDowell, J. A.; Turner, D. H. Investigation of the structural basis for thermodynamic stabilities of tandem GU mismatches: solution structure of (rGAGGUCUC)2 by two-dimensional NMR and simulated annealing. *Biochemistry* **1996**, *35* (45), 14077-14089.
(32) Velagapudi, S. P.; Li, Y.; Disney, M. D. A cross-linking approach to map small molecule-RNA binding sites in cells. *Bioorg Med Chem Lett* **2019**, *29* (12), 1532-1536.
(33) Schneider, C. A.; Rasband, W. S.; Eliceiri, K. W. NIH Image to ImageJ: 25 years of image analysis. *Nat Methods* **2012**, *9* (7), 671-675.
(34) Rumbaugh, G.; Sillivan, S. E.; Ozkan, E. D.; Rojas, C. S.; Hubbs, C. R.; Aceti, M.;

Kilgore, M.; Kudugunti, S.; Puthanveettil, S. V.; Sweatt, J. D.; et al. Pharmacological Selectivity Within Class I Histone Deacetylases Predicts Effects on Synaptic Function and Memory Rescue. *Neuropsychopharmacology* **2015**, *40* (10), 2307-2316.
